# Asymmetric Allostery in Estrogen Receptor-α Homodimers Drives Responses to the Ensemble of Estrogens in the Hormonal Milieu

**DOI:** 10.1101/2024.04.10.588871

**Authors:** Charles K. Min, Jerome C. Nwachukwu, Yingwei Hou, Robin J. Russo, Alexandra Papa, Jian Min, Rouming Peng, Sung Hoon Kim, Yvonne Ziegler, Erumbi S. Rangarajan, Tina Izard, Benita S. Katzenellenbogen, John A. Katzenellenbogen, Kendall W. Nettles

## Abstract

The estrogen receptor-α (ER) is thought to function only as a homodimer, but responds to a variety of environmental, metazoan, and therapeutic estrogens at sub-saturating doses, supporting binding mixtures of ligands as well as dimers that are only partially occupied. Here, we present a series of flexible ER ligands that bind to receptor dimers with individual ligand poses favoring distinct receptor conformations —receptor conformational heterodimers—mimicking the binding of two different ligands. Molecular dynamics simulations showed that the pairs of different ligand poses changed the correlated motion across the dimer interface to generate asymmetric communication between the dimer interface, the ligands, and the surface binding sites for epigenetic regulatory proteins. By examining binding of the same ligand in crystal structures of ER in the agonist versus antagonist conformers, we also showed that these allosteric signals are bidirectional. The receptor conformer can drive different ligand binding modes to support agonist versus antagonist activity profiles, a revision of ligand binding theory that has focused on unidirectional signaling from ligand to the coregulator binding site. We also observed differences in the allosteric signals between ligand and coregulator binding sites in the monomeric versus dimeric receptor, and when bound by two different ligands, states that are physiologically relevant. Thus, ER conformational heterodimers integrate two different ligand-regulated activity profiles, representing new modes for ligand-dependent regulation of ER activity.

**Significance:** The estrogen receptor-α (ER) regulates transcription in response to a hormonal milieu that includes low levels of estradiol, a variety of environmental estrogens, as well as ER antagonists such as breast cancer anti-hormonal therapies. While ER has been studied as a homodimer, the variety of ligand and receptor concentrations in different tissues means that the receptor can be occupied with two different ligands, with only one ligand in the dimer, or as a monomer. Here, we use X-ray crystallography and molecular dynamics simulations to reveal a new mode for ligand regulation of ER activity whereby sequence-identical homodimers can act as functional or conformational heterodimers having unique signaling characteristics, with ligand-selective allostery operating across the dimer interface integrating two different signaling outcomes.

## Introduction

The nuclear receptor (NR) superfamily includes some NRs that act as heterodimers partnered with retinoid X receptor (RXR) (1, 2), but the estrogen receptor-α (ER) and other steroid receptors comprise a subgroup of these transcription factors that are thought to act as homodimers (3, 4). NRs contain globular domains for DNA- and ligand-binding (DBDs and LBDs), the latter of which also contains a structurally conserved binding site for epigenetic regulatory proteins, called Activation Function-2 (AF-2) (5). By binding to the LBD, the ligand regulates the structure of ER to drive dimerization, select DNA binding sites, and control the AF-2 site and other protein interaction surfaces (6, 7), enabling it to assemble an ensemble of interacting coregulators that modify chromatin structure and regulate transcription (7–13). This unidirectional regulatory connection between ligand—through the LBD—to coregulator recruitment is thought to underlie the diverse and ligand-selective effects of estrogens on reproduction, cancer, bone, muscle, metabolism, and cognition.

While allostery between ligand binding and the coregulator binding site has been studied in the context of individual receptor monomers (8, 9, 14–21), it has not been investigated with respect to cross-dimer signaling in ER homodimers or of asymmetrical ER dimers, those only partially occupied by ligand or bound by two different ligands (3, 22). Such ER species with varied stoichiometries are likely involved in the actions of ER in vivo, in complex environments with widely varying levels of different endogenous and exogenous estrogens as during pregnancy, hormone therapy, and environmental exposures (23–26).

A barrier to studying asymmetrical ER signaling and ligand mixtures has been the lack of tools to isolate and study the relevant liganded complexes with mixtures of ligands or ligand poses. In the current work, we isolated the canonical antagonist conformers of the LBD crystallized with a series of ligands with a range of activity profiles. This designed set of flexible ligands bound to ER with different ligand conformations in each monomer of the dimer, mimicking the binding of two different ligands. Here, we observed *receptor conformational heterodimers* with coupled sequence-identical monomers showing unique allostery across the dimer interface. We performed molecular dynamics simulations (MDS) with these structures, and extended the simulations for models of monomers, single-liganded dimers, and dimers bound to both an agonist and an antagonist. We find that asymmetrical allosteric communication occurs between the sequence-identical monomers of ER conformation to integrate information from different oligomeric states and liganded species, representing new means for ligand regulation of ER activity. Our findings provide insight into the diverse modes by which ligands may regulate ER in complex *in vivo* environments, resulting in the diverse activities of natural and synthetic estrogens.

## Results

### Ligand design strategy and ligand activities

To explore new ways that ER homodimers might engage in asymmetrical interactions with resulting functional consequences, we prepared a set of ER ligands with conformational flexibility to interact with the two sequence-identical ER monomers in both different ligand orientations and receptor conformations. We based these new compounds on an oxabicycloheptene sulfonamide (OBHSN) core ligand (17, 27), to which we added a phenol and a series of other substituents, R1 or X onto the sulfonamide nitrogen (see **Fig. 1A–B; *SI Appendix*, Fig. S1A–B** for ring designations in these ligands). The E-ring, found in most ER antagonists, provides a launching point for side chains that exit the LBD between helices 3 (h3) and h11, directly displacing h12 of the AF-2 surface to block coregulator binding (8, 9) (**Fig. 1A; *SI Appendix*, Fig. S1B**). The F-ring, characteristic of these OBHSN core ligands, as well as the appended R1 and X groups, are directed at the start of the h11-h12 loop or to h11 itself and modulate the disposition of h12—and antagonist effects—encased within the ligand binding pocket to regulate surface activity indirectly, which we call indirect antagonism (**Fig. 1A–B; *SI Appendix, Fig. S1B***) (17).

**Figure 1.**
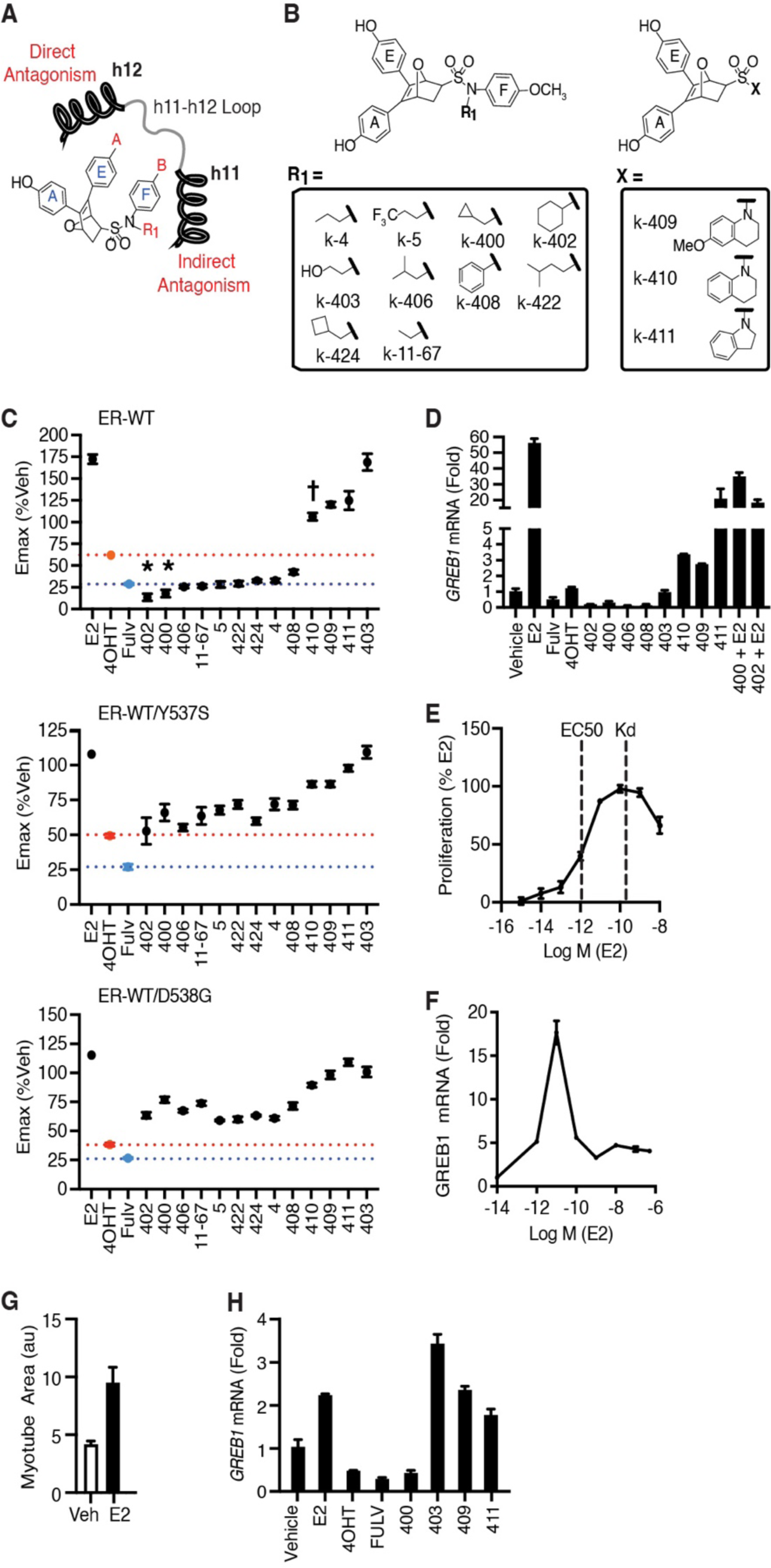
OBHSN scaffold compounds display mixed agonist-antagonist activity. **A)** Chemical structure of OBHSN scaffold and its orientation with respect to h11 and h12 of the ER LBD and noting the location of the F-ring. **B)** Chemical structures of OBHSN compounds. **C)** Maximum efficacy (Emax) from dose-response curves for compound inhibition of proliferation in WT MCF-7 or engineered ER^WT/Y537S^ and ER^WT/D538G^ cells. Datapoints are mean±SEM, n=6 from two biological replicates from dose curve data from two different cell passages. The orange and blue dots and lines represent the activity levels of 4-hydroxytamoxifen (4OHT) and fulvestrant (Fulv), respectively. *, p < 10^-6^ compared to Fulv; †, not significantly different from DMSO. Data analyzed with 1-way ANOVA with Dunnett test. **D)** Steroid deprived MCF-7 cells were treated with 1 μM ligands for 24h and analyzed by qPCR. Vehicle level of gene expression is set at 1. Values are mean ± SD from three biological replicates. **E)** MCF-7 cells cultured in charcoal-stripped fetal bovine serum were treated with E2 for 5 days and analyzed for cell number. Kd was calculated as described in the Methods through displacement of [^3^H] E2 from recombinant ER. Values are mean ± SEM from two biological replicates. **F)** Primary female mouse skeletal muscle stem cells were differentiated into multi-nucleated myotubes in steroid free conditions and treated with E2 for 24 hrs for qPCR analysis. Mean +SEM was calculated with N = 2. **G)** Myotubes were treated with 50 pM E2 for 5 days. Tube area was measured by high content image analysis. Mean + SEM, N = 16. **H)** Gene expression studies were repeated in MCF-7 cells growing normally in full media. Data is mean + SEM of n = 5 replicates.

The newly synthesized OBHSN compounds (***SI Appendix*, Scheme S1**) showed affinities, reported as Ki values, that are in the sub-μM range of 0.0054 – 0.220 μM (***SI Appendix*, Table S1**). They were profiled for estrogen responsive luciferase activity and for inhibition of growth of MCF-7 breast cancer cells with ER-WT or MCF-7 cells engineered to have one allele of the most common constitutively activating mutations found in metastatic treatment-resistant breast cancer, ER-Y537S or ER-D538G (**Fig. 1C; *SI Appendix*, Tables S1–2**) (21). We identified ligands k-402 and k-400 as showing maximal efficacy (Emax) significantly better than fulvestrant in inhibiting proliferation of MCF-7 cells, with many of the other compounds showing efficacy equivalent to fulvestrant, the ER antagonist used to treat metastatic treatment-resistant breast cancer (**Fig. 1C**). K-403, k-411 and k-409 profiled as partial agonist/antagonists, while k-410 was neutral, showing neither growth stimulatory nor inhibitory effects. In the two mutant ER cell lines, 4-OHT (4-hydroxytamoxifen, an active metabolite of tamoxifen) and the OBHSN ligands showed similar patterns as with the WT ER but reduced antagonist efficacy (**Fig. 1C; *SI Appendix, Fig. S2A***). Potency was not correlated between cell lines, but the Y537S mutant cells showed significantly reduced potency compared to the D538G cells (***SI Appendix,* Fig. S2B–C**).

To probe the activity of the compounds to regulate transcription, we measured expression of ER target genes in MCF-7 cells. The OBHSN inhibitors downregulated expression of the ER-regulated genes, *PGR*, *GREB1*, and *TFF1*/pS2 below that of the control vehicle cells (**Fig. 1D; *SI Appendix*, Fig. S2D**). For k-400 and k-402, we cotreated with E2, which reversed their inhibitory effects (**Fig.1D, *SI Appendix*, Fig.S2D**). Compounds k-409, k-410, and k-411 were agonists, with upregulation of *GREB1* and *TFF1*, and *PGR* mRNA levels. Thus, the pharmacophores around the two substituents on the sulfonamide nitrogen of the OBHSN compounds directed a range of ER-mediated cell proliferation and transcriptional activities that we studied below with crystal structures and molecular dynamics simulations.

We also explored the concentration ranges over which E2 shows activity in cells. E2 induced cell growth at doses 2 logs below the Kd of 0.1 nM for E2 binding (**Fig. 1E**), which is similar to the Kd of 0.1–0.5 nM for receptor dimerization (28–30). By definition, at the Kd concentration ER is 50% occupied by E2, giving a ligand-bound fraction of 1-10% at 1-10 pM, with a similar fraction of ER as a dimer. Skeletal muscle is another important target tissue for ER activity (31). As with growth of MCF-7 cells treated with E2, *Greb1* expression was biphasic with an inflection point below the ligand Kd in primary female mouse myotubes (**Fig. 1F**). E2 maintained myotube diameter at a 50 pM dose (**Fig. 1G**), further supporting the physiological relevance of E2 at doses with limited ER occupancy.

The effects of low dose E2 suggest the receptor can bind two different ligands. The combination of the SERM bazedoxifene and conjugated estrogens have unique properties not seen with either treatment alone (26, 32), leading to FDA approval as Duavee for prevention of both bone loss and menopausal symptoms. To explore why k-403 was not an agonist in the gene expression studies (**Fig. 1D; *SI Appendix,* Fig. S2D**), we first showed that its agonism in proliferation is reversed by fulvestrant (***SI Appendix*, Fig.S2E**). We then tested its effects in the context of ∼ 1pM E2 found in media containing intact FBS. As expected, E2 showed less induction of gene expression, but k-403 profiled as an agonist (**Fig.1H; *SI Appendix*, Fig.S2F**). We also observed genes that showed differential responses to fulvestrant in the presence of low dose E2 (***Appendix*, Fig.S2G**), unlike the classical ER target gene *GREB1* that responded similarly with E2. Non-monotonic dose and low dose responses are common in the steroid receptors (33, 34) by unknown mechanisms (35). Based on these fundamental pharmacological principles and other studies (3, 22–26, 32), ER activity can derive from ligands bound to a monomer or to dimers as ligand/apo complexes, and respond uniquely to mixtures of ligands (25, 26, 32, 34–36), species which we characterize below.

### Crystal structures reveal formation of conformational heterodimers of both ligand and receptor

We obtained crystal structures of 8 OBHSN ligands with ER in the antagonist conformation (***SI Appendix*, Table S3**), with electron density maps for the ligands shown in ***SI Appendix*, Table S2**. The flexible elements of the OBHSN ligands extending from the nitrogen demonstrated an array of orientations and receptor interactions. The F-ring adopted three different binding modes: the F-ring facing *out* towards the h11-h12 loop, *down* in between h11 and h8, or *back* towards h8 (**Fig. 2A–C; *SI Appendix,* Table S4**). Many of the LBD dimers showed ligands in two different poses, i.e., *ligand conformational heterodimers*, with some ligands stabilizing the same receptor conformation, while other ligands generating different conformations in each subunit of the dimer, i.e., *receptor conformational heterodimers*.

**Figure 2.**
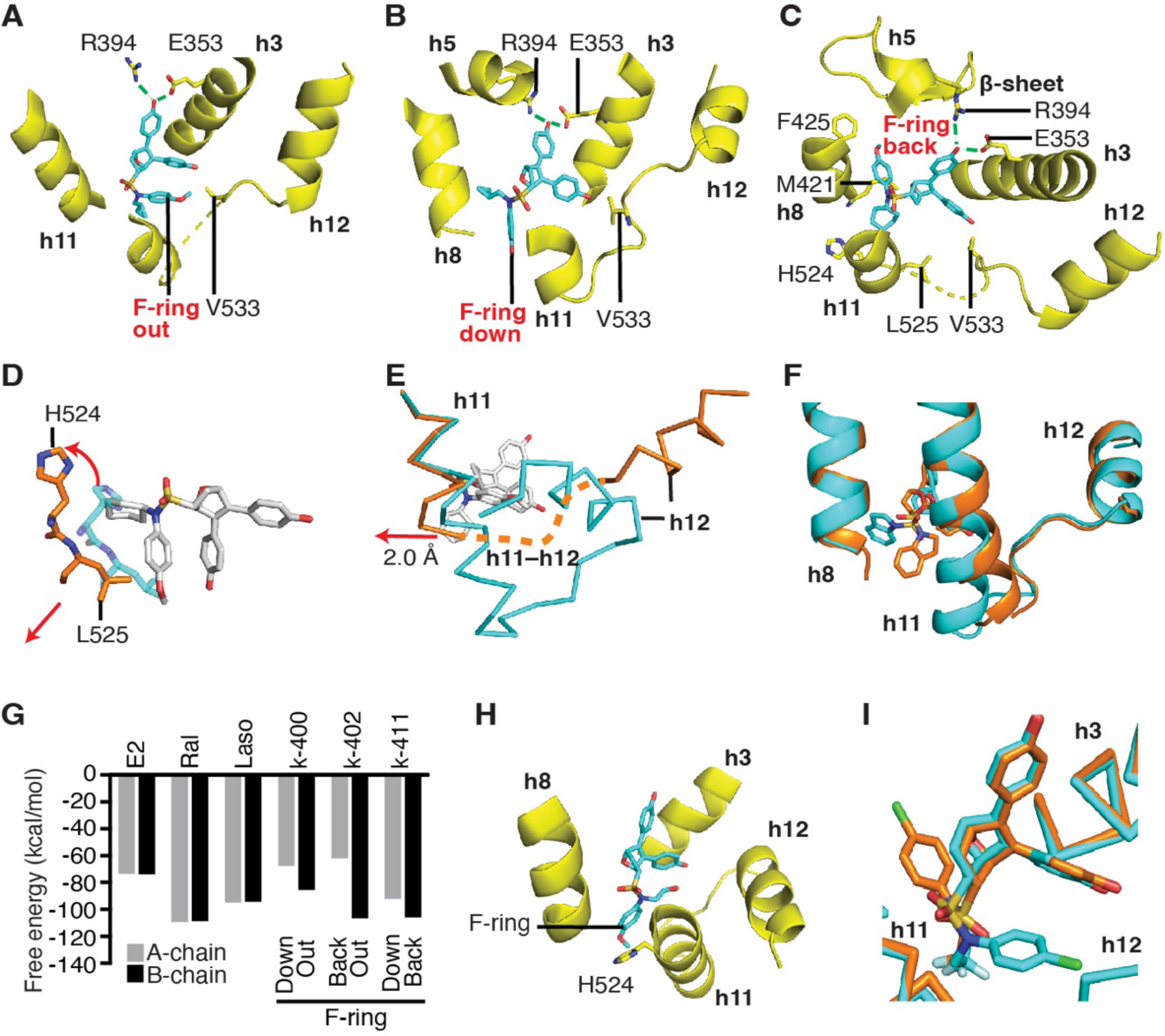
Structures of OBHSN compounds in the antagonist conformer of ER. **A)** Structure of ER LBD in complex with k-400 with F-ring facing out towards h12 **B)** In the dimer partner, k-400 adopts a different conformation, where the F-ring points down towards the C-terminus of h11. **C)** Structure of ER LBD in complex with k-402 shows the F-ring pointing back towards h8 and the β-sheet. **D)** The structure of ER bound to k-402 (orange) was superimposed on the agonist conformation structure with E2 (teal) showing the ligand induced shift in helix 11. **E)** Structure of ER LBD in complex with k-402 shows the F-ring pointing back towards h8 and the β-sheet (orange) with the C-terminus of h11 and the h11-h12 loop disordered (orange dashed line), compared to the E2 (teal) bound structure. **F)** Structure of k-411 with the ER A and B chains superimposed showing different positions of the R group and h11. **G)** MM-GBSA calculations of binding free energy of the indicated ligands bound to ER. **H)** Docking of k-403 into an agonist conformation ER structure. **I)** The structurally related full antagonist 13 showed different positions of the F-ring when crystallized with the antagonist conformation receptor (orange) and the agonist conformation receptor (blue). The chemical structure of ligand 13 corresponds to the OBHSN structure shown in Fig. 1B, with R_1_ = CF_3_CH_2_-, A = OH, and B = Cl (17).

With the antagonist k-402, the ligand bound with the F-ring out and the cyclohexane pushing on the backbone of h11 in between H524 and L525 in 3 of the 4 monomers comprising the two dimers in the asymmetric unit (**Fig. 2D**). In the fourth monomer, the positions of the F-ring and cyclohexane were switched, but still stabilized a similar shift in h11, with differences becoming apparent in molecular dynamics simulations, described below. These interactions contributed to an overall shift of h11 by 2 Å compared to the agonist conformer, destabilizing the end of h11 and the h11/12 loop so that they were not resolved in the structure, but indirectly destabilized h12 to drive antagonism (**Fig. 2D–E**). Heterogeneity of ligand conformations and receptor conformations was also found with agonist OBHSN ligands. K-410 and k-411 adopted multiple conformations across the different subunits (***SI Appendix,* Table S4**), either down or back, but here we saw two different positions of h11, representing coupling between ligand conformational heterodimers and receptor conformational heterodimers (**Fig. 2F**).

To evaluate how these different ligand binding poses relate to the activity state of the receptor, we used molecular mechanics with generalized Born and surface area solvation (MM-GBSA) methods to calculate the differential free energy of ligand binding. The antagonist conformer in which the F-ring is out, parallel to the E-ring, was most energetically favorable for the antagonists k-400 and k-402. In contrast, the backwards conformer was energetically favored over the downwards R group conformer with the agonist, k-411 (**Fig. 2G**). Thus, the outward orientation of the F-ring—with the ensuing displacement of h11—is energetically favored with the indirect antagonists but disfavored with the agonist. This supports a model in which the binding of one ligand transmits allosteric information thats predisposes a different ligand binding pose in the dimer partner. In contrast to the energy differences between the two subunits of these OBHSN conformational heterodimers, estradiol, raloxifene, or lasofoxifene bound to the ER LBD as homodimers and show no differences in binding energy between the subunits of the dimer (**Fig. 2G**).

### Reciprocal control of ligand and receptor conformations across the dimer interface with bidirectional allostery

Current dogma is that NR signal transduction flows unidirectionally, where the bound ligand acts by stabilizing an LBD conformation that modulates AF-2 activity. We found that allostery between ligand and coregulator binding sites is bidirectional, with receptor conformation also driving different ligand binding modes. The hydroxyethyl substituent on the sulfonamide of k-403 showed the F-ring group facing outward in all subunits of the two dimers modeled in the antagonist conformation structure (***SI Appendix,* Table S4**). We used flexible sidechain docking to probe how k-403 bound to the agonist conformation structure. In the full agonist conformation of the receptor, the F-ring of the agonist k-403 is preferentially positioned downwards into the pocket of space between h8 and h11 instead of out towards h12 (**Fig. 2H**), demonstrating that different receptor conformers drive different binding modes of the ligand. The crystal structure of a related full antagonist **13**, which we previously obtained with ER in the agonist conformer (25), is shown here bound to ER in the antagonist conformer (**Fig. 2I**). The structure of compound **13** has a CF_3_CH_2_-group at the R1 substituent and an ortho-Cl-group on the F-ring. The different substates of ER (i.e., agonist vs antagonist conformer) stabilized the ligand F-ring in different positions, either back into the pocket or out towards h12, demonstrating that receptor conformations can determine different ligand binding poses.

### Mapping ligand-selective allosteric communication across the dimer interface

To further characterize conformational heterodimerization and effects on h11, we superimposed the B chains onto the A chains from the dimer for each structure and then measured the distance between backbone amides or sidechains of H524 and L525 to compare the positioning of h11 between the receptor monomers (**Fig. 3A**). E2, raloxifene, and lasofoxifene showed almost identical monomers, with distances < 0.5 Å. In contrast, most of the OBHSN structures showed conformationally heterogeneous positions of h11 compared with E2 (**Fig. 3A**). With k-409, k-410 and k-411, we observed that one chain shifted away from h12 (***SI Appendix,* Fig. S3A**, blue), representing an indirect antagonist conformation by pulling on the h11-12 loop and destabilizing h12. However, the other chain in the dimer showed h11 shifted towards h12 (***SI Appendix,* Fig. S3A,** yellow), a conformation that is consistent with the neutral to agonist activity of these ligands. Thus, ER displayed conformational heterodimers where each monomer showed coupled ligand-receptor conformations with known, distinct activity profiles of h11 positioning.

**Figure 3.**
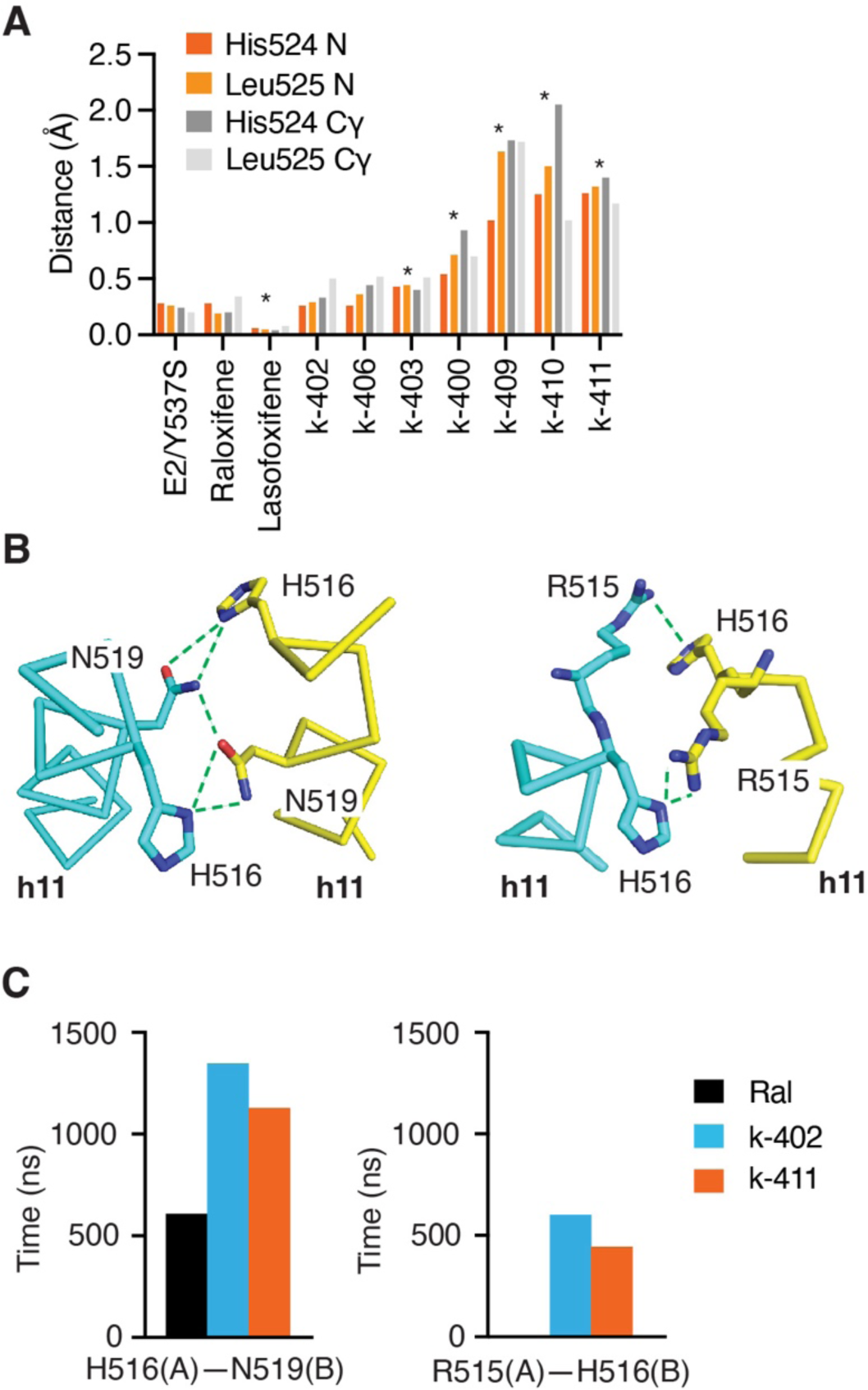
Ligand selective dimer conformers. **A)** B chains were superimposed onto the A chains of the ER structures with the indicated ligands. Distances between the indicated backbone amide (N) or sidechain (Cγ) were measured between the A and B subunits of each of the structures. * Significantly different from E2 bound structure by 1-way ANOVA. **B)** Structure of k-402-bound ER shows hydrogen bond-mediated networks across the dimeric interface for H516–N519, and R515–H516 **C)** Histogram shows the total time of H516–N519, and R515–H516 interactions during the MDS.

To further understand the mechanisms and consequences of conformational heterodimerization, we performed all atom molecular dynamics simulations (MDS). Looking at the distribution of the h11 dimer states, we found that the L525–L525 distance across the dimer interface was shorter in the k-402 and k-411 structures than with raloxifene-bound ER by ∼1.5Å (***SI Appendix*, Fig. S3B**), consistent with the shifts in h11 seen in the crystal structures. The interactions across the dimer interface include a series of intermolecular hydrogen bonds from amino acids S512 to E523, which are adjacent to the ligand contacts at G521, H524, and L525 (**Fig.3B–C; *SI Appendix*, Fig. S3C**). We measured the resident times of these contacts at 3.3 Å distance and found that H-bond interactions were highly populated through most of the simulations with k-402 and k-411, but much less so with raloxifene (**Fig. 3C**). These data support a model where the different positions of h11, transmitted through ligand contacts with G521, H524 and L525, affect H-bonds between R515, H516 and N519 across the h11 dimer interface.

### Correlation network analysis reveals ligand-selective allosteric signaling mechanisms

A correlated motion network was generated from the respective MDS to identify routes of communication between the two ligands in the dimer. Individual residues are represented as nodes in the network with edges reflecting correlated motion between different residues, and the weight of the edges corresponds to correlation value. Raloxifene communicated through a direct and symmetrical route across the dimer interface between the regions of h11 adjacent to the ligand, between G521 and Y526 (**Fig. 4A**), including a water-mediated H-bond network between N519, K520, and E523 (**Fig. 4B**).

**Figure 4.**
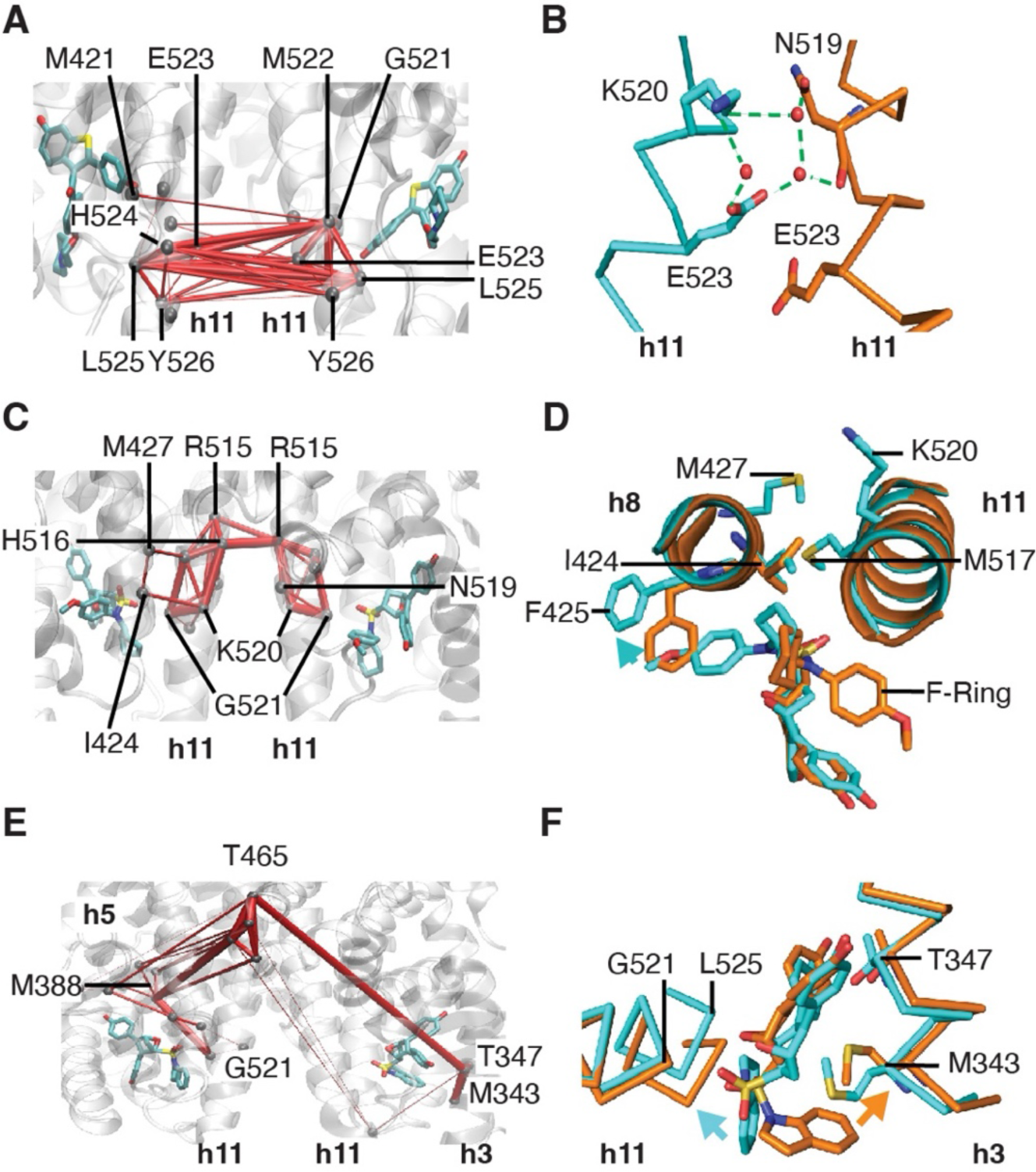
Dynamics of h11 and h12 and communication pathways between ligands. **A)** Network analysis of pathways for correlated motion between the two raloxifene ligands of each subunit (PDB: 2QXS). Red edges indicate identified pathways and weight of edges signify strength of correlation. **B)** Water bridged network observed in the crystal structure of raloxifene-bound ER. **C)** Pathways of correlated motion between the two k-402 ligands in the dimer of each subunit. **D)** A (cyan) and B (orange) subunits of k-402-bound ER are superimposed. Pathway analysis reveals allosteric signaling across h8 and h11. **E)** Pathways of correlated motion between the two k-411 ligands of each subunit. **F)** K-411 ligand conformational heterodimers have differential effects on the positioning of h11 and h3.

Pathway analysis with k-402/ER showed different communication through the center of h11. This included a region between G521 and R515, with an asymmetrical distribution of interactions where H516 in the A chain communicated with R515 in the B chain, but not vice versa (**Fig. 4C**). The F-ring in the back position elicited additional nodes involving h8 I424 and M427, which contact residues in h11 (**Fig. 4C**). The back position of the F-ring shifted the position of F425 on the opposite face of h8 from I424 and M427 (**Fig. 4D**), suggesting that this contact directs the ligand-selective allosteric network.

The agonist k-411 showed greater asymmetry of communication between ligands. This was through a more indirect route, from h11 to h5 of the coactivator binding site, and then across the dimer interface to the loop between h9 and h10, and finally to residues contacting the other ligand in h3 (**Fig. 4E**). The down position of the R group shifted G521 and showed a larger change in H524 and L525 positions at one side of the allosteric network, while shifts in the position of the E-ring altered the positioning of h3 including T437 (**Fig. 4F**). These analyses show that cross dimer signaling has ligand-selective characteristics and is asymmetrical, producing ligand and/or protein conformational heterodimers.

### Accelerated MDS identification of ligand-specific conformational substates

To probe whether the ligands are switching binding modes in situ or only upon initial binding, we measured the movement of the F-ring relative to the E-ring for ER bound to k-411 or k-402 with accelerated MDS (aMDS) (***SI Appendix*, Fig. S4**), enabling sampling of larger conformational changes that occur on longer time scales. With k-402, the A-ring to F-ring ligand distance showed two distinct, non-overlapping positions, consistent with selection of the protein/ligand conformer upon ligand binding (**Fig. 5A**). The time course of ligand motion for k-402 showed little variance in the E-ring to F-ring distance, being different but stable in the A and B chains of ER (***SI Appendix*, Fig. S4B**). Examining the total grid boundary that the ligand occupied throughout the simulation confirmed this behavior with k-402 exhibited small fluctuations of the ligand E-ring to F-ring distance that would not support switching conformers in situ (***SI Appendix*, Fig. S5A**).

**Figure 5.**
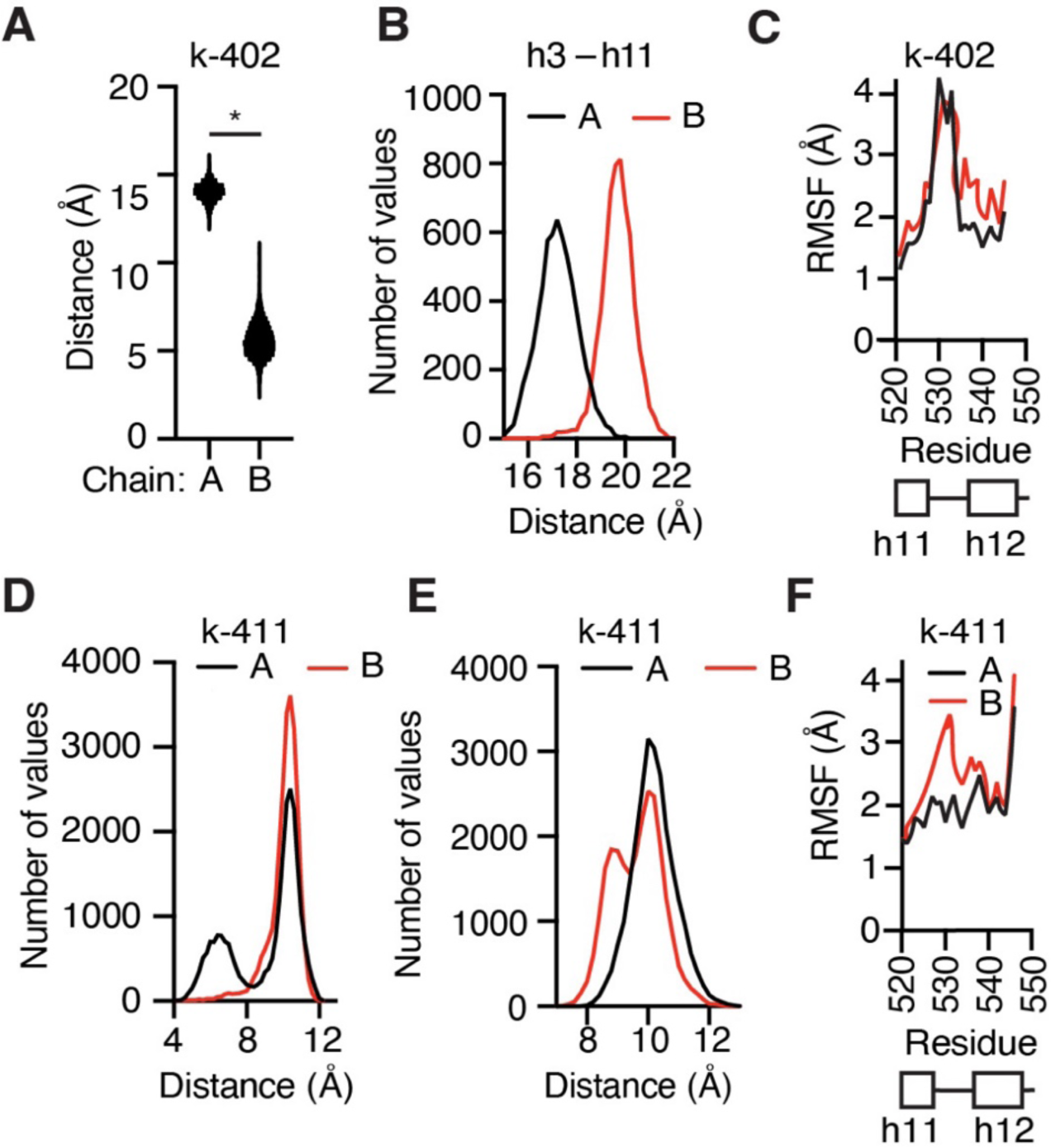
Enhanced sampling through accelerated MDS show ligand-receptor dynamics. **A)** E-ring to F-ring distance for the A and B chains of ER bound to k-402. *Student’s T-test, p < 1 x 10^-11^. **B)** Histogram distribution of the distance of h11 L525 to h3 E353 in the k-402 bound ER aMDS. **C)** Root mean square fluctuation (RMSF) of α-carbons for the A or B chains of the aMDS ER dimer bound to k-402, showing residues in the end of h11 through h12. **D)** Histogram of the E-ring to F-ring distances for k-411 in the A and B chains of ER. **E)** Histogram distribution of the distance from h8 M424 to h11 H524 in the k-411 bound ER aMDS. **F)** Root mean square fluctuation (RMSF) of α-carbons for the A or B chains of the aMDS k-411/ER dimer.

To assess monomer-selective dynamics of h11 in the conformational heterodimers, we calculated the distance between h3 and h11. With k-402 bound ER, there were substantial and largely non-overlapping differences in h11 positioning (**Fig.5B; *SI Appendix,* Fig. S5B**), where the F-ring out position of the ligand led to a more open h3–h11 position of the receptor. With k-402, but not raloxifene, there were differences in the fluctuations in h12 between chains (**Fig.5C; *SI Appendix,* Fig. S5C**), demonstrating that the differences in ligand and h11 positioning were transmitted to the AF-2 surface. The non-overlapping positions of the ligand and h11 in the dimer subunits support semi-stable receptor substates in this conformational heterodimer.

The ligand and receptor interconverted between substates with k-411. The E-ring to F-ring ligand distance was closer in the A chain (**Fig. 5D; *SI Appendix,* Fig. S5D**), which showed a distribution between two ligand positions. However, in the bimodal distribution seen in the aMDS, most of the time was spent with the ligand in the more extended conformer with the R group facing back into the pocket (**Fig. 5D**). Examining the total grid boundary that the ligand occupied throughout the simulation confirms this dynamic behavior for k-411 specifically (***SI Appendix*, Fig. S5E**). For k-411 bound to the B chain, the extended ligand conformer showed brief incursions into the rare more condensed conformer over time (***SI Appendix*, Fig. S4C**). In the A-chain, k-411 showed longer times of occupancy of both ligand conformers during the first half of the simulation. After 3 switches between binding modes, k-411 settled into a similar pattern as seen in the B chain, with the ligand largely in the extended conformer (***SI Appendix*, Fig. S4C**). This demonstrated a form of structural memory, where the receptor retained conformational aspects of the compact ligand binding form while the ligand rotates into the extended form.

With the receptor, the k-411 bound B chain showed a bimodal distribution of h11 relative to h8 (**Fig. 5E; *SI Appendix*, Fig. S5F**), switching between a more compact form and a more extended, open conformer. The RMSF with k-411 showed an overall increase in h11 dynamics in the B chain (**Fig. 5F**), consistent with the extended, bi-modal distribution of h11. The conformational stability of these two types of ligand dynamics makes physical sense: With its larger substituents, the k-402 ligand has a more stable conformation while bound to ER, while k-411 has a smaller substituent and can interconvert while still bound. Differences in ligand binding in the conformational heterodimers thus translated to differences in the key regulators of activity, including the h11-12 loop and the h12 portion of the AF-2 coactivator binding site.

### Allostery between agonist and the AF-2 coactivator binding site is altered in ligand heterodimers

To study the effects of mixed ligand complexes, we superimposed the WT ER lasofoxifene (Laso)-structure on the WT E2-bound structure to generate an E2/Laso-bound ER dimer, and then used this ligand heterodimer for comparison with the E2/E2 and Laso/Laso ligand homodimers. MDS of the mixed E2/ Laso dimer showed that Laso broadly increased dynamics in the E2-bound partner subunit, including the AF-2 surface, h11, and the SRC-2 peptide, relative to the E2/E2-bound structure (**Fig. 6A–B**). The combination of agonist and antagonist in a mixed dimer generated widespread increased dynamics in both chains in the dimer (**Fig. 6A–B; *SI Appendix*, Fig. S6A– B**). Despite the increased dynamics, the average structure showed a reduced area of the AF-2 surface in the E2-bound monomer in the mixed agonist-antagonist dimer (**Fig. 6C**). To assess the functional consequences of these changes in the AF-2 surface, we performed steered MDS to measure the amount of force required to remove the SRC-2 peptide from the surface. More work was required to remove the coactivator peptide from the E2/E2 ER dimer than from the E2/Laso ER dimer, demonstrating altered allostery in the ligand heterodimer partner (**Fig. 6D**).

**Figure 6.**
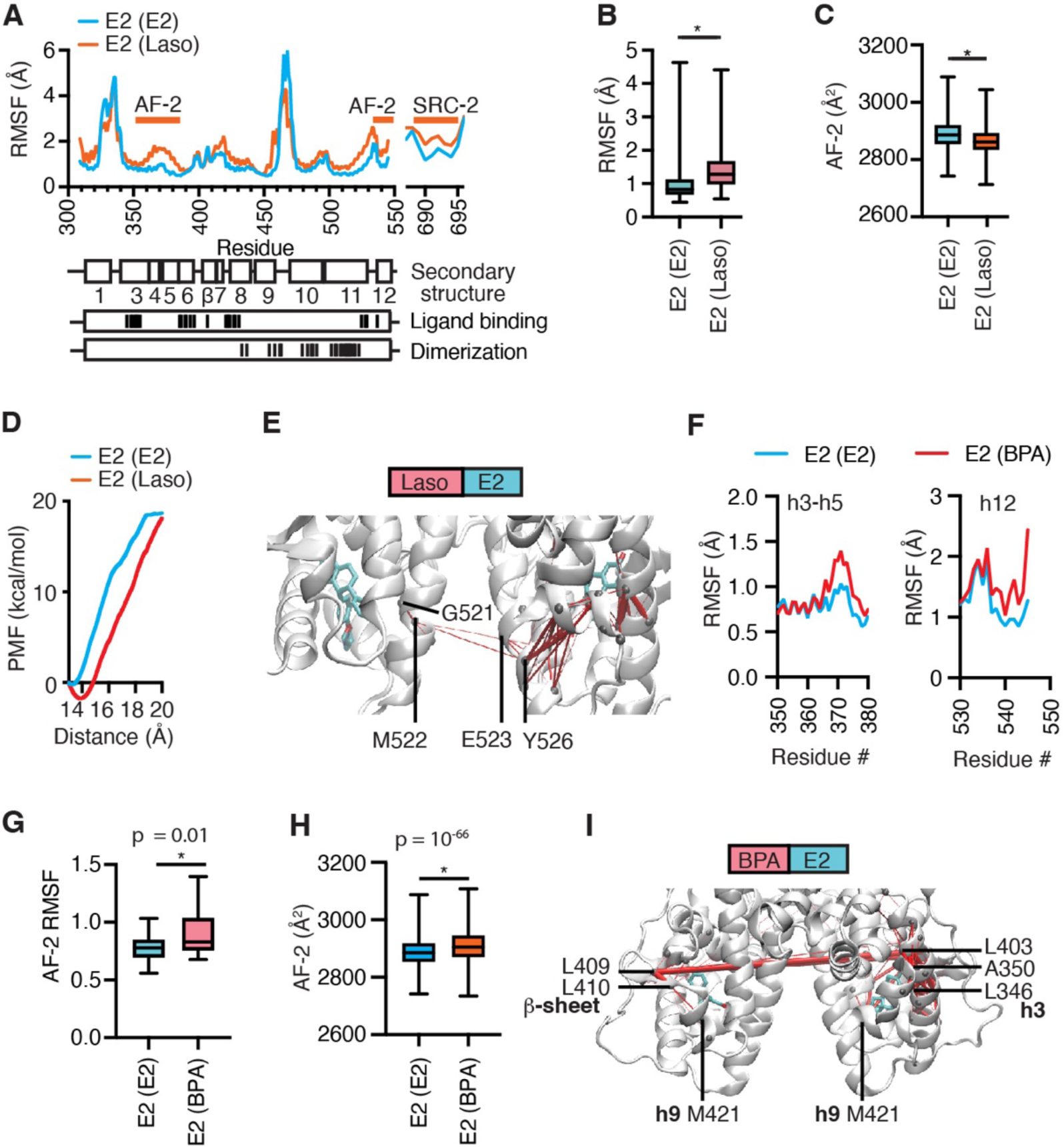
MDS of ER bound to two different ligands shows allosteric signaling across the dimer. **A)** The E2-bound WT ER-LBD (1gwr.pdb) was superimposed with the lasofoxifene(Laso)-bound ER (2ouz.pdb) to generate a mixed agonist/antagonist dimer with E2 and Laso. The RMSF of the MDS is shown for the A chain bound to E2 with the E2/E2 or E2/Laso bound dimers. **B)** Box plot of the average RMSF from the aMDS with min/max whiskers. * Student’s T-test, p = 4 x 10^-12^. **C)** Box plot of the average area of the AF-2 surface from the aMDS with min/max whiskers. Surface area was calculated from ER residues within 4 Å of the SRC-2 peptide using AMBER. * Student’s T-test, p = 1 x 10^-93^. **D)** Steered MDS was used to measure the force to remove the SRC-2 peptide from the AF-2 surface of the E2-bound LBD with either E2 or Laso-bound to the dimer partner. * Student’s T-test, p < 2 x 10^-12^. **E)** Pathways of correlated motion (suboptimal pathway analyses) between the ligands in the ER dimer were calculated from MDS. **F)** The structure of BPA with ER in the agonist conformer (3uu7.pdb) was used to generate an E2/BPA heterodimer for MDS analysis. The E2 bound side of the heterodimer was compared to the E2/E2 structure and RMSF was calculated. **G)** The RMSF was calculated for the AF-2 helices 3-5 and h12 and shown as a box plot for E2 with either E2 or BPA as the partner ligand. **H)** The surface area of the AF-2 surface residues was calculated, including helices 3-5 and h12. **I)** Pathways of correlated motion (suboptimal pathway analyses) between the ligands in the ER dimer were calculated from MDS.

Suboptimal pathway analysis of correlated motion between ligands showed the E2/E2 dimer communication was symmetrical through ligand contacts with G521 and H524 that were transmitted to R515 and H516 in the h11 part of the dimer interface (***SI Appendix,* Fig. S6C**). The E2/Laso dimer showed little allosteric communication between the two ligands (**Fig. 6E**). This contrasted with MDS of E2/bazedoxifene heterodimers, which showed asymmetrical correlated motion including communication between the h11-h12 loop in the bazedoxifene monomer and G420 in h9 of the E2 monomer (***SI Appendix,* Fig. S6D**), but little differences in RMSF. E2/4-hydroxytamoxifen liganded heterodimers showed a different pattern, with symmetrical correlation motion across dimer, but enhanced dynamics of h3–h5 of the AF-2 surface and SRC-2 peptide (***SI Appendix,* Fig. S6E**). The three antagonist/E2 heterodimers showed different patterns of asymmetrical allostery, but together demonstrate that ligand ER heterodimers have unique signaling properties.

Environmental estrogens are often present as complex mixtures with unique activity profiles (35, 36), where ligands with no effect at low dose can combine to drive uterine proliferation, for example (36). To test whether asymmetry occurred between two different agonist ligands, we generated heterodimeric ER bound to E2 and the environmental estrogen, BPA, with both h12’s in the agonist conformer and bound to SRC-2 peptides. The E2 monomer with BPA as a ligand partner showed increased dynamics of h3-5 and h12 of the AF-2 surface, and increased surface area of AF-2 (**Fig. 6G–H**). These experiments provide structural explanations for how mixtures of estrogens can have unique outcomes.

### Differential dynamics of monomeric versus dimeric ER

We performed MDS with structures from k-402 or k-411 bound to ER monomers, and to single-liganded dimers where one ligand was removed, leaving a ligand in either the A chain only or the B chain only (***SI Appendix*, Fig.S7**). Both k-402 and k-411 showed that compared to monomeric ER (black trace), the single-liganded dimers (red trace) stabilized the h11 portion of the dimer interface between residues 500–520 to a greater degree (***SI Appendix*, Fig.S7**). K-411 showed dramatic differences in h11–h12 depending upon the different binding modes of the ligand in the A versus B chains, and dimerization (***SI Appendix,* Fig.S7**), defining oligomeric status as a ligand-selective regulator of the AF-2 surface.

To probe for more pronounced effects of monomeric versus doubly liganded dimers, we ran aMDS with monomeric A or B chains from the structures of k-402 and k-411 bound ER (**Fig. 7A–D**) and compared them with the results from the two-liganded dimer receptor simulations (**Fig. 5**). With k-402 bound to both subunits in the dimeric receptor there was greater dynamics in h12 and h3 of the AF-2 surface, the SRC-2 peptide, and in the h10–h11 part of the dimer interface compared to the monomer (**Fig. 7A**). In contrast, with k-411 we observed the opposite—that monomeric ER showed greater fluctuations across the receptor compared to the dimer (**Fig. 7B**). We analyzed ligand and h11 substates in the aMDS of monomeric receptors (**Fig. 7C-D**) to compare with the aMDS analysis of the dimer (**Fig.5**). With k-402, the A chain showed a mixture of conformers in the aMDS of the monomer (**Fig.7C, red arrow**), but a single conformer in the dimer (**Fig. 5B**). With k-411, the monomeric receptor showed two overlapping populations of ligand conformers in both chains, with increased population of the more compact form of the ligand in the monomer (**Fig. 7D, red arrow**). The monomeric B chain showed that the h8-h11 distance contained an additional, expanded conformer (**Fig. 7D, blue arrow**) compared to simulation of the dimer (**Fig. 5E**). These studies demonstrate that monomeric ER has different allostery between ligand and coregulator binding sites than singly and doubly liganded dimeric ER. These analyses provide another example of signaling in the reverse direction, as dimerization changes the ensemble of ligand conformers for k-411, as well as the conformational dynamics of the surface coregulator binding site.

**Figure 7.**
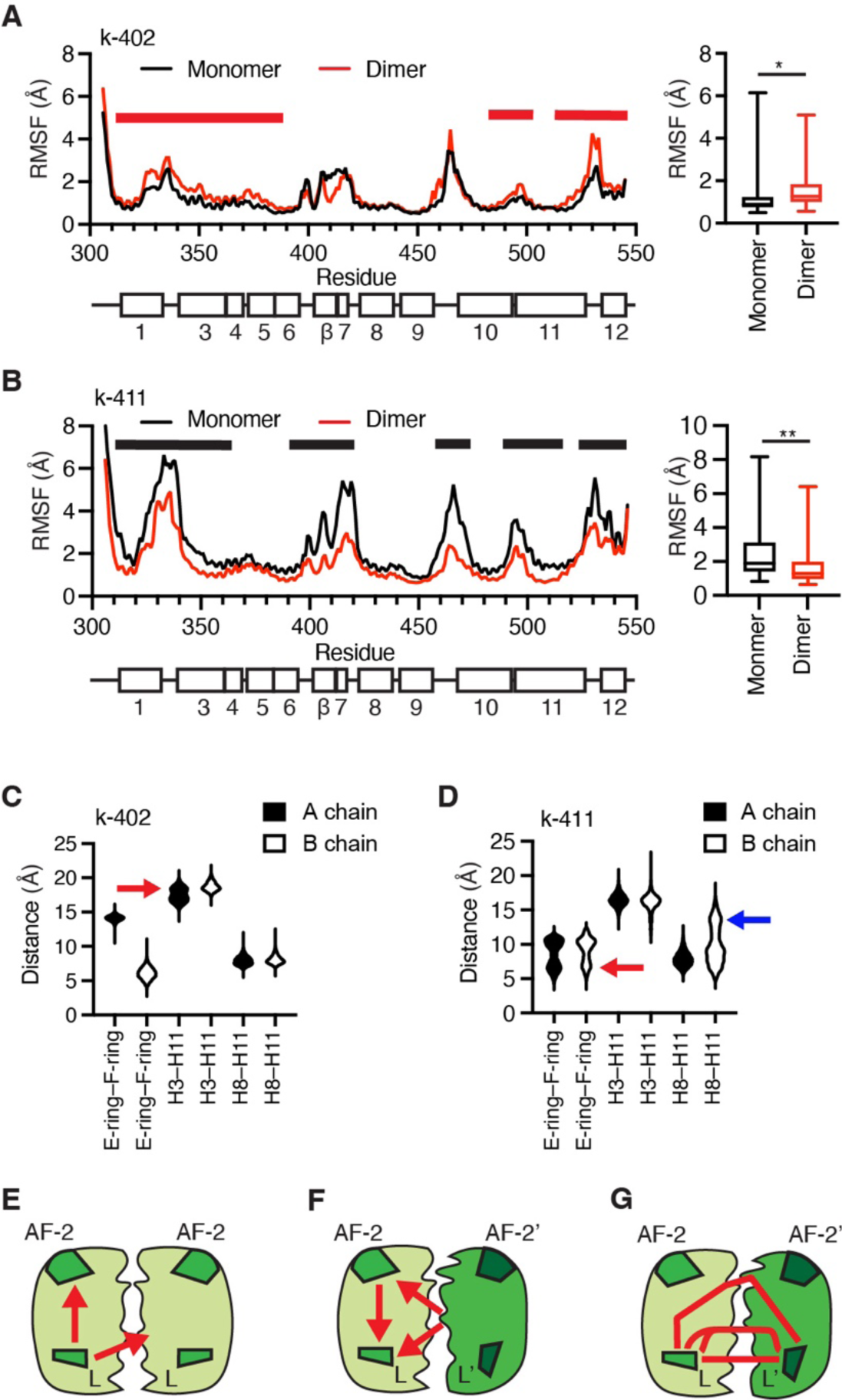
MDS of Monomeric vs Dimeric ER. **A–B)** The structures of ER bound to k-402 or k-411 were analyzed as monomers or dimers with aMDS. * Student’s T-test, p = 2 x 10^-36^; ** p= 2 x 10^-51^. Bold horizontal bars indicate regions of maximum differences; red for k-102, black for k-411. **C–D)** Monomeric receptors were analyzed with aMDS for effects on ligand and h11 conformers, as assessed by the distance from h11 H524 to h3 E353 or h11 H525 to h8 M424. Arrows indicate significant differences from dimeric receptors (shown in Fig. 5). **E)** Inside-out allostery is shown in a cartoon of the ER LBD *homodimer*, illustrating ligand (L) regulation of the coregulator binding site (AF-2) and dimerization. (red arrows). **F)** Outside-in signaling is illustrated in a cartoon of *conformational heterodimers* of ER bound to two different ligands or to the same ligand bound in two conformations, showing effects of the dimer partner (darker green) regulating ligand binding or AF-2 across the dimer interface (red arrows). **G)** Cartoon of 3 routes of correlated motion (red lines) between ligands in conformational heterodimers (detailed in Fig. 4).

## Discussion

The current model for steroid hormone action is that the receptors act as homodimers to recruit coregulators that regulate transcriptional programs (6–13). Here, we identified structural underpinnings of how ligands regulate *asymmetrical* ER complexes, including conformational heterodimers with one or two bound ligands, and monomeric receptors. When visualized with X-ray crystallography, the OBHSN ligands induced conformational heterodimers where the individual ligands and/or receptor monomers showed different ligand conformations in the dimer, mimicking the binding of two different ligands. We also found that ER in either the agonist or antagonist conformer led to altered ligand binding interactions, demonstrating that allostery and conformational heterodimerization are bidirectional (**Fig. 7E-G**). This represents a revision to ligand binding theory that has focused on unidirectional signaling, i.e., namely *inside-out signaling*, whereby ligand binding to the LBD affects the coregulator binding surface (**Fig. 7E**), as opposed to *outside-in signaling*, in which the LBD conformation alters the orientation of bound ligand (**Fig. 7F**), as we have newly demonstrated.

To understand underlying mechanics of this allosteric signaling and conformational dynamics, we performed molecular dynamics simulations (MDS) with the OBHSN-bound and other ER structures, and extended the simulations for models of monomers, single-liganded dimers, and mixed agonist/antagonists including enhanced sampling methods. We also observed outside-in signaling in the monomer versus dimer aMDS with k-411, where dimerization altered the distribution of ligand conformers. Ligand communication in protein conformational heterodimers occurred through asymmetrical coupled motion in the receptor, reflecting ligand-selective effects through the dimer interface (**Fig. 7G**). Protein conformational heterodimers presented structural differential transcriptional coregulator binding surfaces compared to heterodimers(**Fig.7F**). This illustrates a new mode by which receptor activity can be modulated, which has not been visualized by previous simulations of monomeric ER (14, 21), or conceptualized within the framework of steroid receptors as homodimer signaling molecules.

Our findings add insight into many of the diverse activities of estrogens and the conformational modes by which they act through ER. Asymmetrical ER complexes are physiologically relevant, as there is a significant population of monomeric ER that can bind with high affinity to E2 (29), as does a dimeric ER engineered to only bind one ligand (22). Apo dimers can also bind DNA, while mutations that drive dimerization of the apo ER generate some constitutive activity and are found in treatment-resistant metastatic breast cancer (37). ER ligands vary in their stabilization of dimerization (28), which we show here is a primary regulator of receptor dynamics and conformational heterodimerization. Number and brightness microscopy data indicate that ER bound to DNA as a mixture of monomers and dimers in contrast to other steroid receptors that formed dimers and higher order oligomers (38), while cryo-EM and other data show there is asymmetry in the bound coregulator complexes (11, 39).

The modern hormonal milieu dictates that ER responds to a variety of metazoan, environmental and phytoestrogens as complex mixtures of estrogens that cover a wide concentration range. Throughout the course of the menstrual cycle, levels of the highest affinity estrogen, 17β-estradiol (E2), vary from low pM to low nM, covering the ∼200-500 pM range of the Kd (i.e. 50% bound) of ER for dimerization and E2 binding. This supports that E2 frequently occupies monomeric ER or has only one ligand bound to the dimer, species that exist in solution (3, 22), or as mixed liganded heterodimers as could occur with pregnancy estrogens.

In addition to the clear role of E2 in activating monomeric or single-liganded dimers based on its cellular concentration, there are many other examples of ligands that are present at non-saturating concentrations, such as the anti-estrogen fulvestrant (40), the naturally occurring antagonist 27-hydroxycholesterol (41), and the variety of environmental and phytoestrogens. However, it is also likely that these sub-saturating ligands can bind together with E2 in the dimer partner to generate conformational heterodimers with two ligands or two different ligand conformers (42), shown by the different gene expression profiles of anti-estrogens given with E2 compared to either ligand alone (25, 26), and the unique effects of mixtures of low dose environmental estrogens (34–36). Our work showed that different ligand-binding conformers or combinations of ligands can regulate the coregulator binding site and receptor dynamics in the ER dimer partner.

In sum, this work identified dimerization and oligomeric status as regulators of ER action, including ligand-selective conformational heterodimerization, providing intriguing new mechanisms by which ER can integrate information from multiple or low dose ligands in the hormonal milieu in vivo.

## Materials and Methods

### Cell culture

MCF7-ERα-Y537S and MCF7-ERα-D538G were a gift from Steffi Oesterreich, University of Pittsburgh Medical Center and were cultured as previously described (17). Primary skeletal muscle stem cells and myotube differentiation were isolated from the leg muscles of 5-week-old female C57BL/6 mice, cultured as previously described (43)

### Luciferase reporter assay

Steroid-deprived MCF7 cells were transfected with a 3xERE-luc reporter and treated with compounds as previously described (44).

### Cell Proliferation Assay

Cells were suspended in steroid-free media supplemented with 10% charcoal-stripped FBS and placed in 384-well plates. The next day cells were treated with compounds and assessed for cell growth 5 days later using CellTiterGlo (Promega).

### RNA isolation and real-time PCR

Real-time PCR was performed using SYBRgreen PCR Master Mix (Quantabio) as described(45).

### Macromolecular x-ray crystallography

The ERα-L372S/L536S mutant LBD was purified crystallized, and the structures solved as previously described (46) using Globalphasing, (47), PHENIX software suite version 1.20(48). COOT was used for model building (49) Images were generated with PyMOL (Schrodinger).

### Classical and accelerated molecular dynamics simulations

Dimers with two ligands were generated by superpositioning the two relevant structures and combining A and B chains from the different structures using COOT (49). Missing portions of ER LBD were modeled in COOT by using other ER crystal structures with the same conformation trapping mutation and crystal packing environment. The Modeller46 extension (50) within UCSF Chimera (51) was used to fill in all other missing parts. 500ns production simulations were run in triplicates for each complex using AMBER.

For accelerated MD, a short 10ns classical MD simulation was used to compute average dihedral and potential energies as reference for the aMD parameters. A dual-boost approach was used by applying independent energy thresholds to enhance sampling (52). The aMD parameters were defined in terms of *E* and α, where *E* represents the total boost energy level and α is a tuning parameter for the acceleration potential. *E*_total_, *E*_dihedral_, α_total_, and α_dihedral_ were calculated using the average *E*_total_ and *E*_dihedral_ from the 10ns classical MD simulation, as detailed in AMBER Advanced Tutorial 22 (https://ambermd.org/tutorials/advanced/tutorial22/).

### Simulation analysis

Trajectory postprocessing was performed with CPPTRAJ (53). The autoimage command was used to recenter the trajectory coordinates and the strip command was used to remove water and ion molecules. The resulting trajectories were analyzed using CPPTRAJ and Bio3D (54). Suboptimal paths between designated “source” and “sink” nodes were drawn with the Bio3D cnapath() function, as previously described (55). The ligands on both sides of the ER-LBD dimer were designated as the source and sink nodes of the path construction. One hundred paths were collected for each pair of nodes that were analyzed. Suboptimal paths between residue sites were visualized as edges in VMD. Betweenness centrality was calculated with Bio3D.

### Potential binding energy calculation

Calculating the potential binding energy of each monomer with MM-GBSA module without flexible residues using Schrodinger Inc. software.

### Statistical Analyses

Statistics were calculated using analysis of variance (ANOVA) with multiple comparisons, or Student’s t-test, as appropriate, using GraphPad Prism 9.5.1 software.

## Acknowledgments

Grants from the National Institutes of Health (R01CA275142 to KWN, and R01CA220284 to KWN, TI, BSK, and JAK), the Frenchman’s Creek Fellowship (to KWN lab), and the Breast Cancer Research Foundation (BCRF-083 to BSK and BCRF-084 to JAK and BSK). Sunping Nie and Xujian Ji of School of Life Sciences, Hubei University, contributed to the synthesis of these compounds. The authors declare no conflicts of interest. Abbreviations: aMDS, accelerated molecular dynamics simulation; DBD, DNA-binding domain; ER, estrogen receptor-α; LBD, ligand-binding domain; MDS, molecular dynamics simulation; NR, nuclear receptor; OBHSN, oxabicycloheptene sulfonamide; PMF, potential mean force, RMSF, root mean square fluctuation; SERM, selective ER modulator.

## Data deposition

PDB coordinate files are deposited as 8VZ0, 8W07, 8VZP, 8VZQ, 8VZ1, 8VYX, 8VYT, 8W03. RNA-seq data set is available at GSE231397.

## References

1. B. M. Forman, K. Umesono, J. Chen, R. M. Evans, Unique response pathways are established by allosteric interactions among nuclear hormone receptors. Cell 81, 541–550 (1995).

2. D. J. Kojetin et al., Structural mechanism for signal transduction in RXR nuclear receptor heterodimers. Nat Commun 6, 8013 (2015).

3. M. A. Miller, A. Mullick, G. L. Greene, B. S. Katzenellenbogen, Characterization of the subunit nature of nuclear estrogen receptors by chemical cross-linking and dense amino acid labeling. Endocrinology 117, 515–522 (1985).

4. J. W. Schwabe, L. Chapman, J. T. Finch, D. Rhodes, The crystal structure of the estrogen receptor DNA-binding domain bound to DNA: how receptors discriminate between their response elements. Cell 75, 567–578 (1993).

5. K. W. Nettles, G. L. Greene, Ligand control of coregulator recruitment to nuclear receptors. Annu Rev Physiol 67, 309–333 (2005).

6. D. M. Heery, E. Kalkhoven, S. Hoare, M. G. Parker, A signature motif in transcriptional co-activators mediates binding to nuclear receptors. Nature 387, 733–736 (1997).

7. C. E. Foulds et al., Proteomic analysis of coregulators bound to ERalpha on DNA and nucleosomes reveals coregulator dynamics. Mol Cell 51, 185–199 (2013).

8. A. M. Brzozowski et al., Molecular basis of agonism and antagonism in the oestrogen receptor. Nature 389, 753–758 (1997).

9. A. K. Shiau et al., The structural basis of estrogen receptor/coactivator recognition and the antagonism of this interaction by tamoxifen. Cell 95, 927–937 (1998).

10. J. D. Norris et al., Peptide antagonists of the human estrogen receptor. Science 285, 744–746 (1999).

11. P. Yi et al., Structural and Functional Impacts of ER Coactivator Sequential Recruitment. Mol Cell 67, 733–743 e734 (2017).

12. P. Yi et al., Structure of a biologically active estrogen receptor-coactivator complex on DNA. Mol Cell 57, 1047–1058 (2015).

13. Y. Shang, X. Hu, J. DiRenzo, M. A. Lazar, M. Brown, Cofactor dynamics and sufficiency in estrogen receptor-regulated transcription. Cell 103, 843–852 (2000).

14. Y. Li et al., A mutant form of ERalpha associated with estrogen insensitivity affects the coupling between ligand binding and coactivator recruitment. Sci Signal 13 (2020).

15. J. B. Bruning et al., Partial agonists activate PPARgamma using a helix 12 independent mechanism. Structure 15, 1258–1271 (2007).

16. S. Srinivasan et al., Ligand-binding dynamics rewire cellular signaling via estrogen receptor-alpha. Nat Chem Biol 9, 326–332 (2013).

17. S. Srinivasan et al., Full antagonism of the estrogen receptor without a prototypical ligand side chain. Nat Chem Biol 13, 111–118 (2017).

18. J. D. Stender et al., Structural and Molecular Mechanisms of Cytokine-Mediated Endocrine Resistance in Human Breast Cancer Cells. Mol Cell 65, 1122–1135 e1125 (2017).

19. N. E. Bruno et al., Chemical systems biology reveals mechanisms of glucocorticoid receptor signaling. Nat Chem Biol 17, 307–316 (2021).

20. J. Min et al., Dual-mechanism estrogen receptor inhibitors. Proc Natl Acad Sci U S A 118 (2021).

21. J. A. Katzenellenbogen, C. G. Mayne, B. S. Katzenellenbogen, G. L. Greene, S. Chandarlapaty, Structural underpinnings of oestrogen receptor mutations in endocrine therapy resistance. Nature Reviews Cancer 18, 377–388 (2018).

22. R. Paulmurugan, A. Tamrazi, T. F. Massoud, J. A. Katzenellenbogen, S. S. Gambhir, In vitro and in vivo molecular imaging of estrogen receptor alpha and beta homo- and heterodimerization: exploration of new modes of receptor regulation. Mol Endocrinol 25, 2029–2040 (2011).

23. C. D. DuSell, M. Umetani, P. W. Shaul, D. J. Mangelsdorf, D. P. McDonnell, 27-hydroxycholesterol is an endogenous selective estrogen receptor modulator. Mol Endocrinol 22, 65–77 (2008).

24. J. F. Robertson, Fulvestrant (Faslodex) -- how to make a good drug better. Oncologist 12, 774–784 (2007).

25. S. E. Wardell, D. Kazmin, D. P. McDonnell, Research resource: Transcriptional profiling in a cellular model of breast cancer reveals functional and mechanistic differences between clinically relevant SERM and between SERM/estrogen complexes. Mol Endocrinol 26, 1235–1248 (2012).

26. T. J. Berrodin, K. C. Chang, B. S. Komm, L. P. Freedman, S. Nagpal, Differential biochemical and cellular actions of Premarin estrogens: distinct pharmacology of bazedoxifene-conjugated estrogens combination. Mol Endocrinol 23, 74–85 (2009).

27. M. Zhu et al., Bicyclic core estrogens as full antagonists: synthesis, biological evaluation and structure-activity relationships of estrogen receptor ligands based on bridged oxabicyclic core arylsulfonamides. Org Biomol Chem 10, 8692–8700 (2012).

28. A. Tamrazi, K. E. Carlson, J. R. Daniels, K. M. Hurth, J. A. Katzenellenbogen, Estrogen receptor dimerization: ligand binding regulates dimer affinity and dimer dissociation rate. Mol Endocrinol 16, 2706–2719 (2002).

29. D. Sakai, J. Gorski, Estrogen receptor transformation to a high-affinity state without subunit-subunit interactions. Biochemistry 23, 3541–3547 (1984).

30. D. F. Skafar, Differential DNA binding by calf uterine estrogen and progesterone receptors results from differences in oligomeric states. Biochemistry 30, 6148–6154 (1991).

31. C. A. Unger et al., Skeletal Muscle Endogenous Estrogen Production Ameliorates the Metabolic Consequences of a High-Fat Diet in Male Mice. Endocrinology 164 (2023).

32. B. S. Komm, S. Mirkin, Incorporating bazedoxifene/conjugated estrogens into the current paradigm of menopausal therapy. Int J Womens Health 4, 129–140 (2012).

33. S. J. Quirk, J. E. Gannell, M. J. Fullerton, J. W. Funder, Mechanisms of biphasic action of glucocorticoids on alpha-lactalbumin production by rat mammary gland explants. J Steroid Biochem 24, 413–416 (1986).

34. A. C. Gore et al., EDC-2: The Endocrine Society’s Second Scientific Statement on Endocrine-Disrupting Chemicals. Endocr Rev 36, E1–E150 (2015).

35. L. N. Vandenberg et al., Hormones and endocrine-disrupting chemicals: low-dose effects and nonmonotonic dose responses. Endocr Rev 33, 378–455 (2012).

36. H. Tinwell, J. Ashby, Sensitivity of the immature rat uterotrophic assay to mixtures of estrogens. Environ Health Perspect 112, 575–582 (2004).

37. S. Irani et al., Somatic estrogen receptor alpha mutations that induce dimerization promote receptor activity and breast cancer proliferation. J Clin Invest 10.1172/JCI163242 (2023).

38. D. M. Presman et al., DNA binding triggers tetramerization of the glucocorticoid receptor in live cells. Proc Natl Acad Sci U S A 113, 8236–8241 (2016).

39. J. Osz et al., Structural basis for a molecular allosteric control mechanism of cofactor binding to nuclear receptors. Proc Natl Acad Sci U S A 109, E588–594 (2012).

40. M. van Kruchten et al., Measuring residual estrogen receptor availability during fulvestrant therapy in patients with metastatic breast cancer. Cancer Discov 5, 72–81 (2015).

41. E. R. Nelson et al., 27-Hydroxycholesterol links hypercholesterolemia and breast cancer pathophysiology. Science 342, 1094–1098 (2013).

42. J. C. Nwachukwu et al., Resveratrol modulates the inflammatory response via an estrogen receptor-signal integration network. Elife 3, e02057 (2014).

43. N. E. Bruno et al., Activation of Crtc2/Creb1 in skeletal muscle enhances weight loss during intermittent fasting. FASEB J 35, e21999 (2021).

44. J. Min et al., Dual-mechanism estrogen receptor inhibitors. Proc Natl Acad Sci U S A 118 (2021).

45. A. Bergamaschi et al., The forkhead transcription factor FOXM1 promotes endocrine resistance and invasiveness in estrogen receptor-positive breast cancer by expansion of stem-like cancer cells. Breast Cancer Res 16, 436 (2014).

46. K. W. Nettles et al., NFkappaB selectivity of estrogen receptor ligands revealed by comparative crystallographic analyses. Nat Chem Biol 4, 241–247 (2008).

47. C. Vonrhein et al., Data processing and analysis with the autoPROC toolbox. Acta Crystallogr D Biol Crystallogr 67, 293–302 (2011).

48. P. D. Adams et al., PHENIX: building new software for automated crystallographic structure determination. Acta Crystallogr D Biol Crystallogr 58, 1948–1954 (2002).

49. P. Emsley, K. Cowtan, Coot: model-building tools for molecular graphics. Acta Crystallogr D Biol Crystallogr 60, 2126–2132 (2004).

50. B. Webb, A. Sali, Comparative Protein Structure Modeling Using MODELLER. Curr Protoc Bioinformatics 54, 5 6 1–5 6 37 (2016).

51. E. F. Pettersen et al., UCSF Chimera--a visualization system for exploratory research and analysis. J Comput Chem 25, 1605–1612 (2004).

52. L. C. Pierce, R. Salomon-Ferrer, F. d. O. C. Augusto, J. A. McCammon, R. C. Walker, Routine Access to Millisecond Time Scale Events with Accelerated Molecular Dynamics. J Chem Theory Comput 8, 2997–3002 (2012).

53. D. R. Roe, T. E. Cheatham, 3rd, PTRAJ and CPPTRAJ: Software for Processing and Analysis of Molecular Dynamics Trajectory Data. J Chem Theory Comput 9, 3084–3095 (2013).

54. B. J. Grant, A. P. Rodrigues, K. M. ElSawy, J. A. McCammon, L. S. Caves, Bio3d: an R package for the comparative analysis of protein structures. Bioinformatics 22, 2695–2696 (2006).

55. X. Q. Yao et al., Dynamic Coupling and Allosteric Networks in the alpha Subunit of Heterotrimeric G Proteins. J Biol Chem 291, 4742–4753 (2016).

