## Supplementary Information for "Asymmetric Allostery in Estrogen Receptor-α Homodimers Drives Responses to the Ensemble of Estrogens in the Hormonal Milieu"

*SI Appendix for:*

**SI Appendix Contents**

**Figures S1-S5…………....2–10**
**Tables S1-S4……………..11–17**
**Scheme S1………………..18**
**Materials and Methods…19–48**
**References………………49**

**Supplemental figures, tables, and schemes**

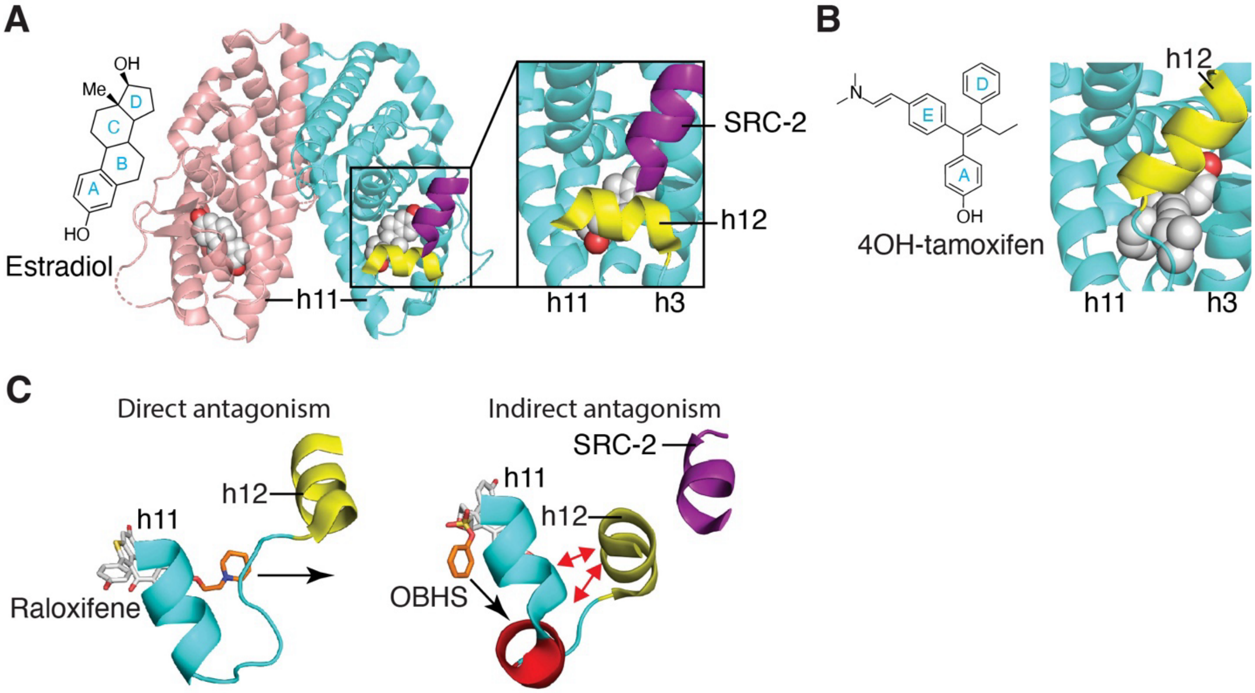

**Figure S1. Structure and pharmacology of the ER LBD.**

**A**) The ABCD ring nomenclature for steroids. Structure of the ER LBD in the agonist conformation, shown as ribbon, with h12 colored yellow and the SRC-2 peptide colored burgundy. Estradiol is shown as space filled.

**B)** Structure of the ER LBD bound to 4-hydroxytamoxifen, shown as space-filled. The receptor is shown as ribbons with h12 colored yellow, docked into the SRC-2 binding site in the antagonist conformation.

**C)** Two mechanisms of allosteric regulation. Helix 11 and helix 12 are shown as ribbons. ER bound to raloxifene shows h12 in the antagonist conformation, where the ligand side chain extends from inside the pocket and directly disrupts the agonist position of h12, which we call ***direct antagonism***. With the oxabicyclic heptene sulfonate (OBHS) ligand, the phenyl side group shifted h11 to indirectly perturb the agonist conformer by disrupting the packing between h11 and h12. This ***indirect antagonism*** can direct a range of graded partial agonist responses. This depends on how far the ligand shifts h11 to control the relative dynamics of h12 between the agonist and antagonist conformers.

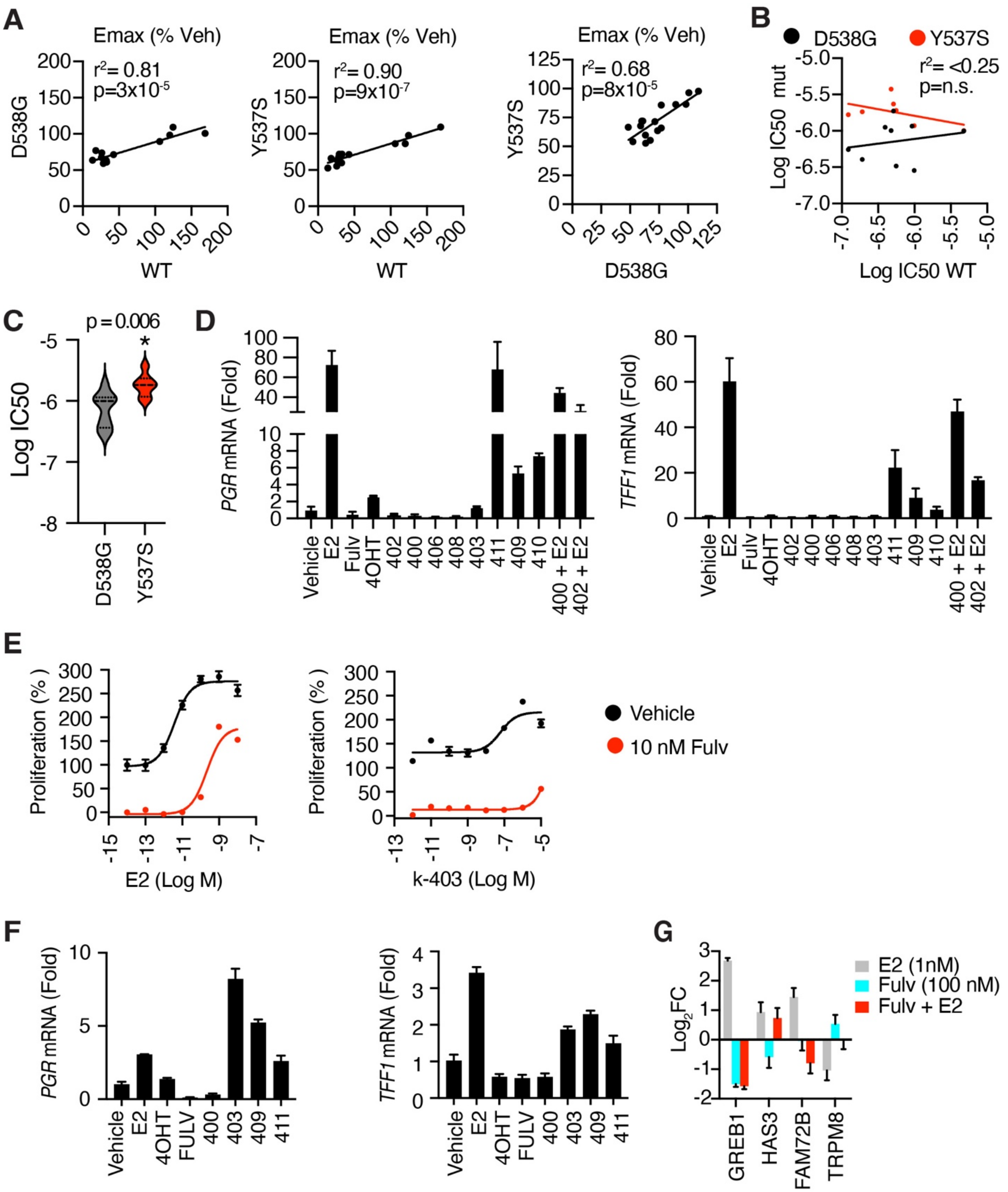

**Figure S2. Ligand pharmacology.**

**A)** The maximal efficacy (Emax) of the ligands for effects on cell growth is shown for MCF-7, MCF-7$^{D538G}$ and MCF-7$^{Y537S}$ cells. The ligands showed very similar patterns across cell lines, as shown by high Pearson correlation r values, but reduced Emax in the mutants.

**BC)** EC50/IC50 values for WT MCF-7 versus ER mutants. D538G cells showed significantly higher IC50 than Y537S cells. **B)** Each point represents potency from one compound. r, Pearson

correlation not significant for both datasets. **C)** Violin plot of data from **B)** analyzed for significance with Student's t-test.
**D)** Steroid deprived MCF-7 cells were treated with 1 μM ligands for 24h and analyzed by qPCR. Vehicle level of gene expression is set at 1. Values are mean ± SD from three biological replicates.Fulv, fulvestrant; 4OHT, 4-hydroxytamoxifen.
**E)** MCF-7 cells were treated with vehicle or 10 nM fulvestrant to suppress proliferation. Cells were co-treated with increasing doses of E2 or k-403. Data are mean+SEM of 3 replicates.
**F)** Gene expression studies were repeated in MCF-7 cells growing normally in full media. Data is mean + SEM of n = 5 replicates.
**G)** MCF-7 cells were grown for 72 hrs in charcoal stripped FBS and treated with the indicated ligands for 24 hrs. RNA isolated for RNA-seq. Data are mean + SEM of n = 3 replicates.

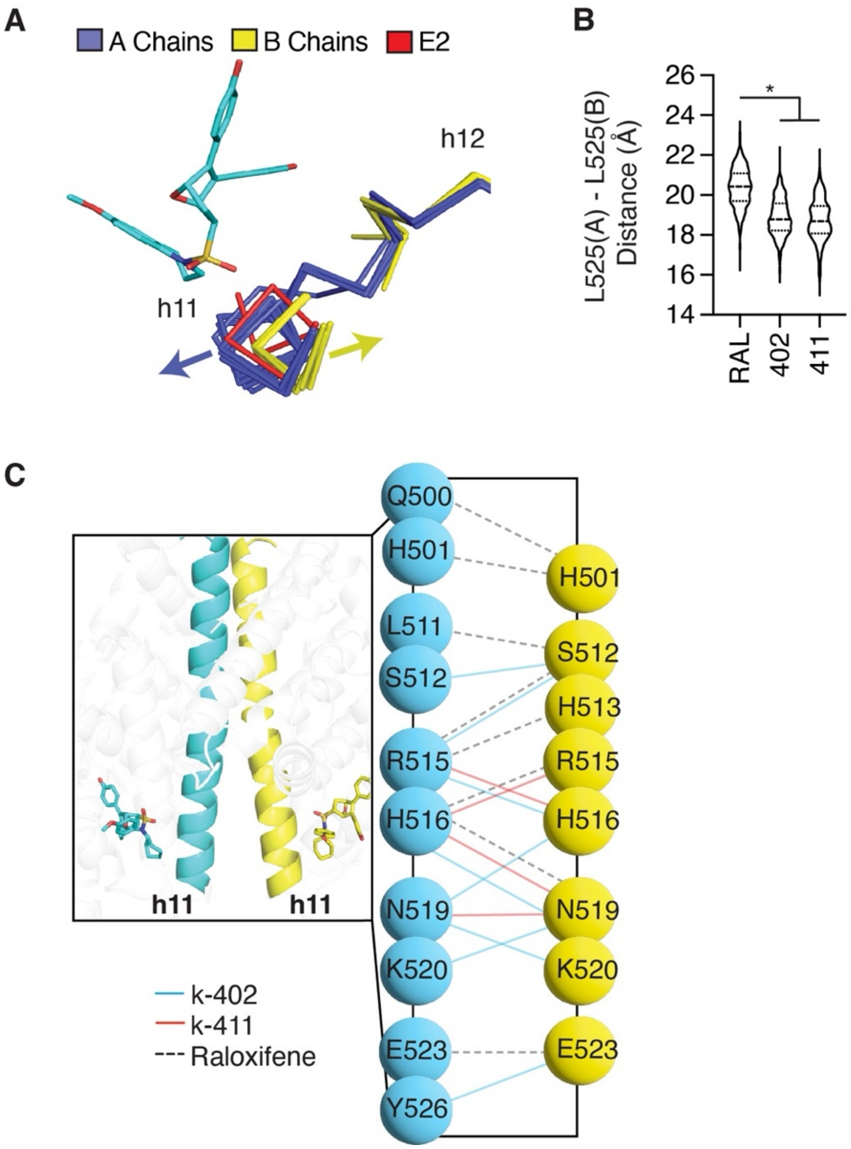

**Figure S3. Helix 11 conformation and analysis of interdimer H-bonds during MDS.**

**A)** H11 positions of each subunit of the neutral/partial agonists, k-409, k-410, and k-411, were superimposed alongside the h11 position of E2-bound ER. The difference in h11 position between the A and B subunits correlate with the disordered nature of the h11-12 loop when h11 is pushed out.

**B)** Violin plot of the distances between L525 across the dimer interface throughout the MD simulations. *One-way ANOVA, $p = 10^{-98}$.

**C)** Contacts less than 3.3 Å between h11 of individual subunits during the MDS were calculated.

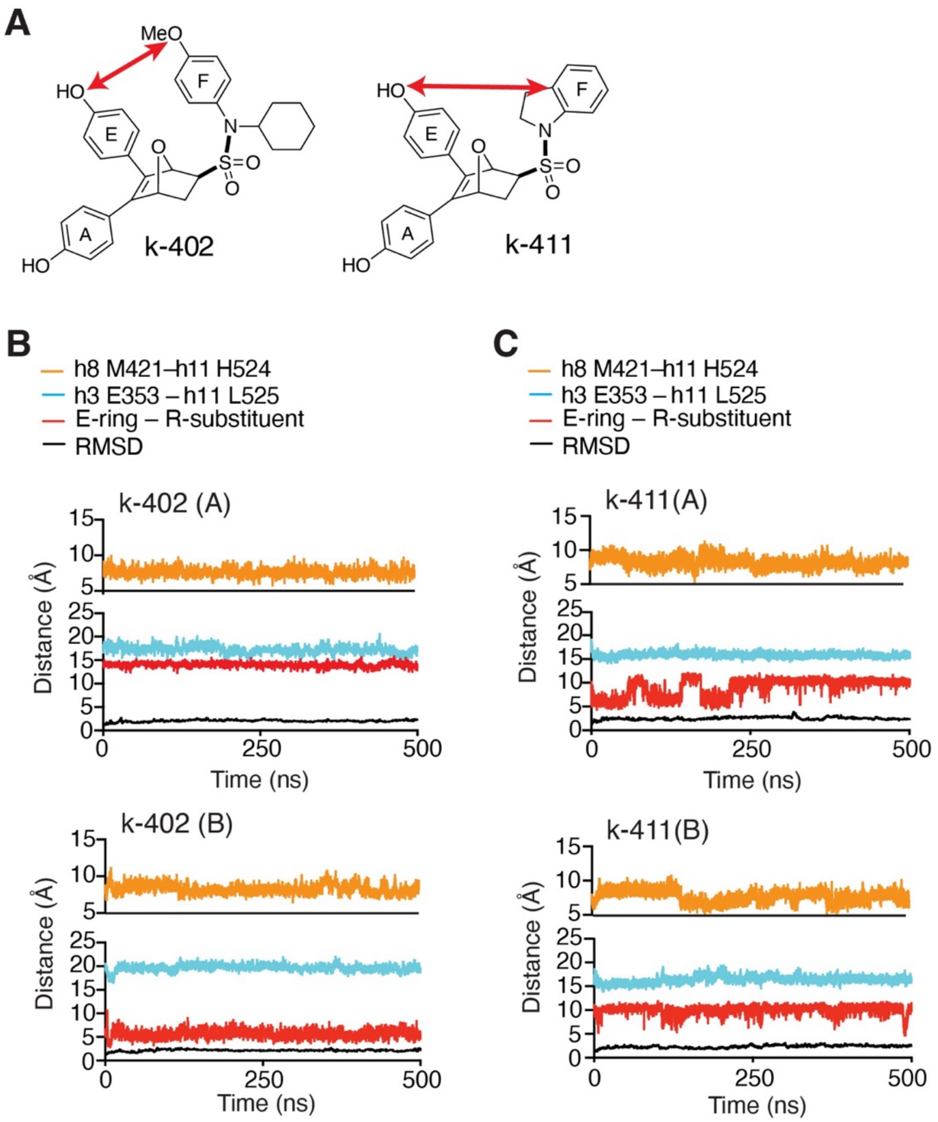

**Figure S4. Enhanced sampling through accelerated MDS show ligand-receptor dynamics time course.**

**A)** Distance analysis of the E-ring to F-ring distance from aMDS of dimers in complex with the indicated ligands for the A and B chains of the dimer.

**B–C)** Distances were measured from the E-ring to F-ring distance and the indicated residues in h11 to either h3 or h8. The RMSD trace indicates that sampling was performed with stable simulations.

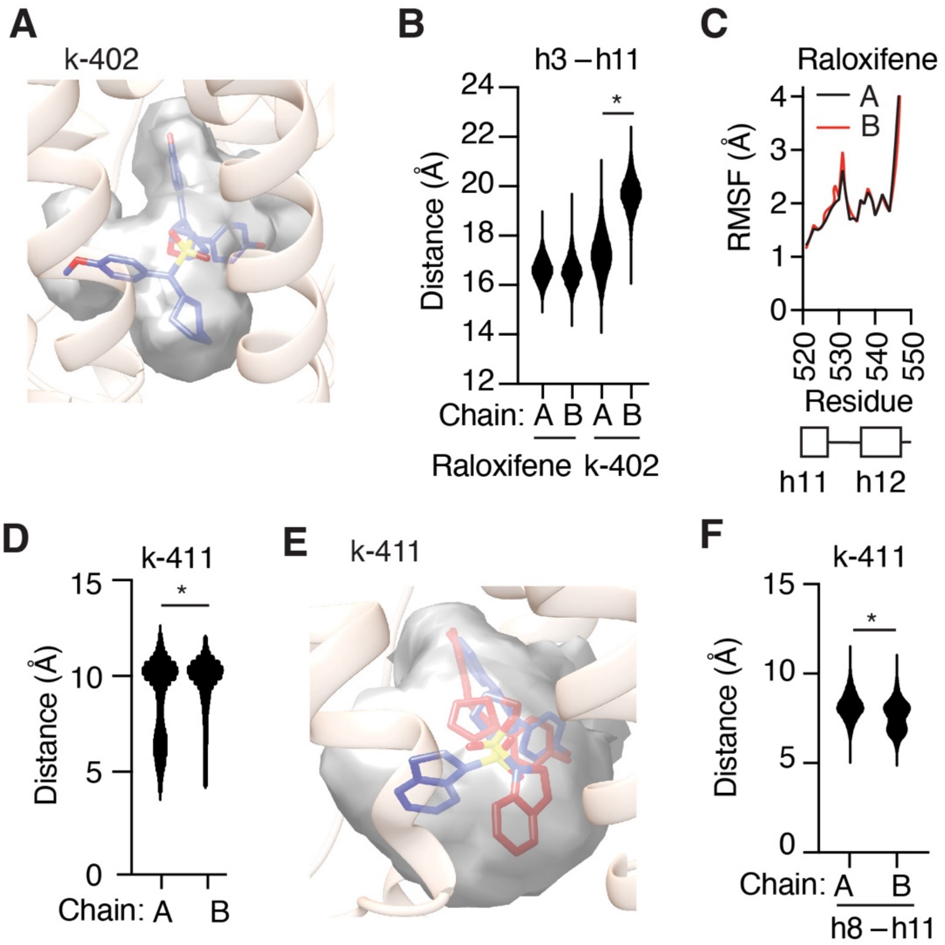

**Figure S5. aMDS analysis of conformational substates**

**A)** Volume occupancy of k-402 throughout the simulation.

**B)** Violin plot of the distance from h3 E353 to h11 L525 during aMDS of ER.

**C)** Root mean square fluctuation (RMSF) of α-carbons from the average for the A or B chains of the aMDS of the raloxifene/ER dimer, showing residues in the end of h11 through h12.

**D)** Violin plot of the E-ring to F-ring distance for the A and B chains of ER bound to k-411.

**E)** Volume occupancy of k-411 throughout the simulation with the two subunits superposed and the ligands colored blue or red.

**F)** Violin plot of the distance from h8 M424 to h11 H524 for k-411/ER during the aMDS.

*Student's T-test, $p < 1 \times 10^{-11}$.

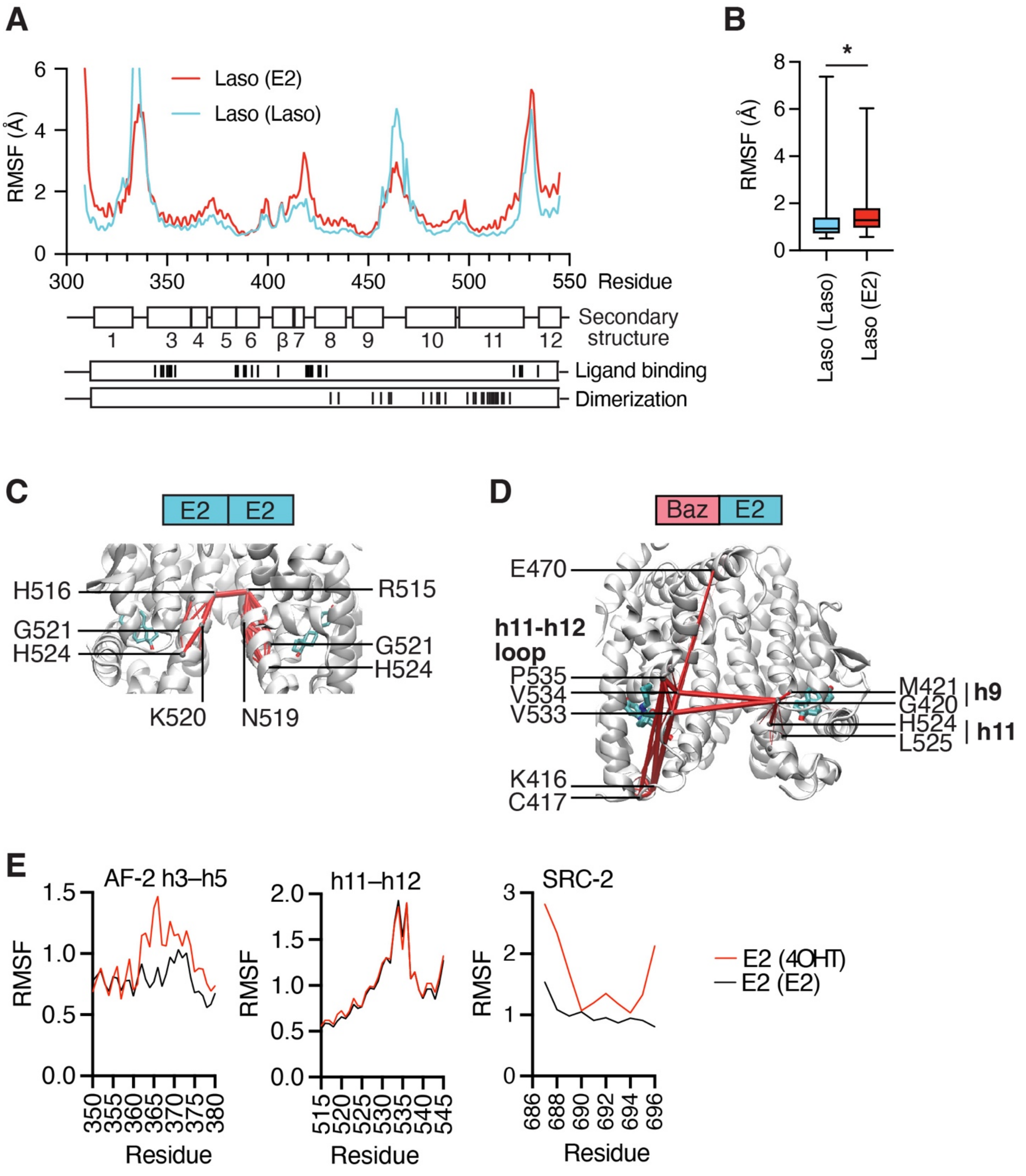

**Figure S6. MDS with E2 and lasofoxifene bound ER.**

**A–B)** RMSF of MDS with lasofoxifene (Laso) ER LBD dimers with either E2 or Laso bound to the dimer partner. The Laso/E2 dimer was generated by superimposing the E2/E2 bound ER with the Las bound ER.

**B)** Box blots with max/min whiskers of the data from **A)**. *Student's T test, $p = 10^{-7}$.

**C–D)** Pathways of correlated motion (suboptimal pathway analyses) between the ligands in the ER dimer were calculated from MDS. Baz, bazedoxifene

**E)** The heterodimer of ER bound to E2 and 4OHT (3ert.pdb) was subject to MDS. The E2-bound monomer was compared to the E2/E2 dimer for RMSF of the indicated regions. This shows that 4OHT binding altered the AF2 surface and SRC-2 dynamics of the E2 bound dimer partner.

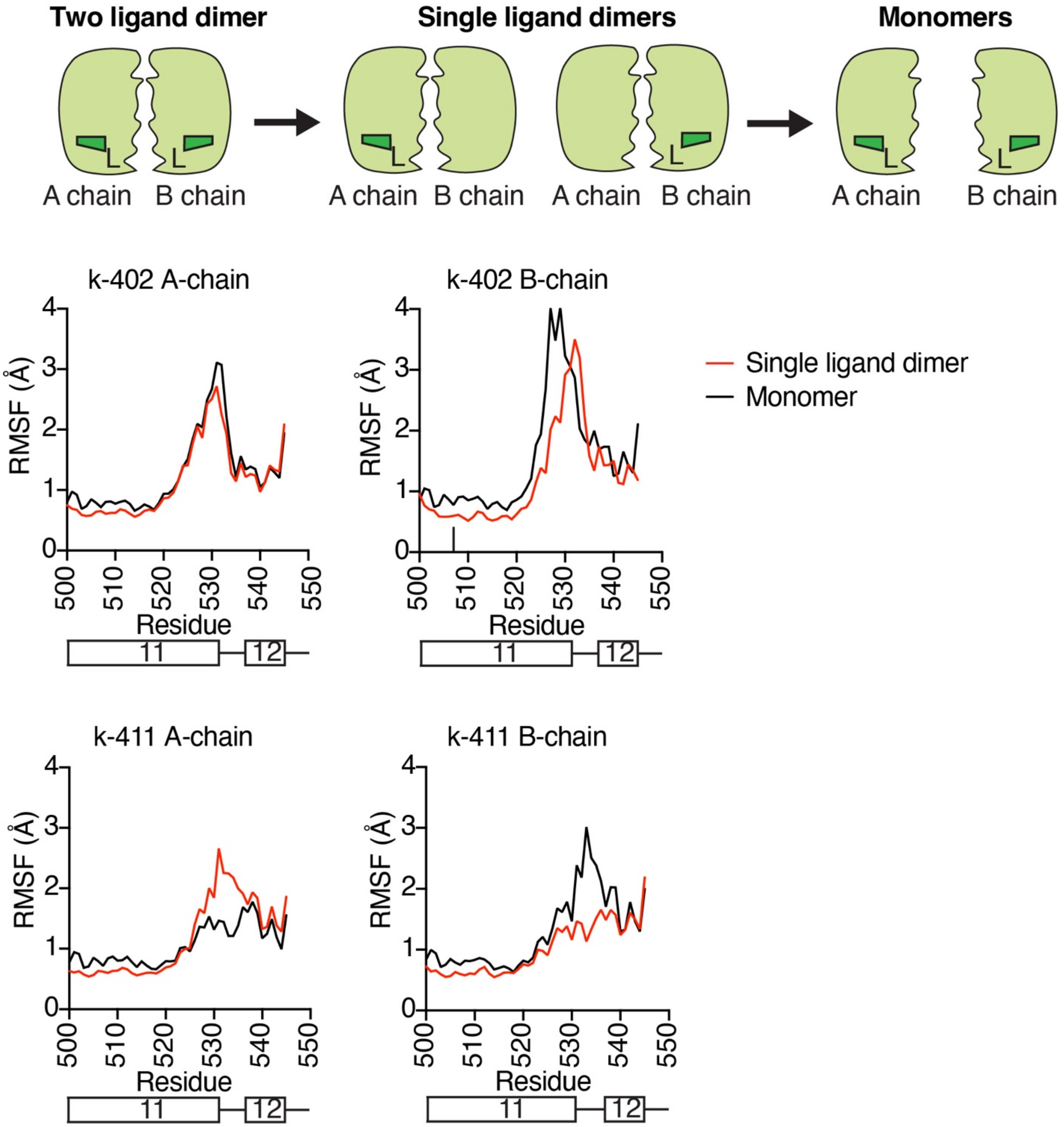

**Figure S7. RMSF of ER MDS with monomeric and single liganded dimers.**
MDS of ER monomers or single-liganded dimers with the ligand in the indicated monomer**.** RMSF showing h11 and h12 residues for the indicated monomers or single liganded dimers, with the chain designating which subunit has the ligand.

| | | Proliferation MCF-7 WT | | Proliferation MCF-7 D538G | | Proliferation MCF-7 Y537S | | 3xERE Luciferase | |
|---|---|---|---|---|---|---|---|---|---|
| Compound | RBA* (Ki, μM) | Emax | LogIC /EC50 ($r^2$) | Emax | LogIC /EC50 ($r^2$) | Emax | LogIC /EC50 ($r^2$) | Emax | LogIC /EC50 ($r^2$) |
| E2 | | 172 | -11.0 (0.98) | 115 | -11.1 (0.83) | 108 | - | 97 | -9.7 (0.96) |
| 4OHT | | 69 | -8.1 (0.93) | 38. | -8.1 (1.00) | 46 | -7.7 (1.00) | 1.1 | -7.7 (0.63) |
| Fulv | | 289 | -9.5 (0.97) | 27 | -8.5 (1.00) | 27 | -8.9 (1.00) | -5.89 | -10.1 (0.91) |
| k-402 | 0.95 (0.021) | 14 | -6.9 (0.97) | 63 | -6.3 (0.99) | 53 | -5.8 (1.00) | -7.03 | -7.5 (0.98) |
| k-400 | 1.76 (0.011) | 178 | -6.0 (0.98) | 77 | -6.5 (0.98) | 66 | -5.9 (0.96) | -7.0 | -6.9 (0.86) |
| k-406 | 1.33 (0.015) | 26 | -6.3 (0.97) | 67 | -6.0 (0.98) | 55 | -5.4 (0.98) | -6.3 | -7.3 (0.86) |
| k-424 | 3.73 (0.005) | 33 | -6.3 (0.95) | 63 | -6.5 (0.98) | 60 | -5.7 (0.96) | -8.4 | -7.5 (0.95) |
| k-422 | 1.43 (0.014) | 29 | -6.3 (0.92) | 60 | -5.7 (0.99) | 72 | -5.6 (0.96) | -8.70 | -7.0 (0.99) |
| k-4 | 1.22 (0.016) | 33 | -6.4 (0.87) | 61 | -6.0 (0.79) | 72 | - | -6.6 | -6.5 (0.83) |
| k-5 | 0.29 (0.069) | 28 | -6.0 (0.72) | 59 | -6.0 (0.75) | 68 | - | -7.9 | -6.9 (0.85) |
| k-408 | 1.86 (0.011) | 43 | -6.7 (0.96) | 71 | -6.4 (0.98) | 71 | -5.7 (0.86) | -6.8 | -7.2 (0.98) |
| k-11-67 | 0.404 (0.050) | 26 | -5.3 (0.98) | 74 | -6.0 (0.98) | 64 | -6.0 (0.99) | -6.6 | -6.1 (0.94) |
| k-403 | 0.09 (0.217) | 169 | -7.2 (0.93) | 101 | - | 109 | - | 3.7 | -8.9 (0.86) |
| k-411 | 1.38 (0.014) | 125 | -7.8 (0.98) | 109 | - | 98 | - | 10 | -7.6 (0.98) |
| k-409 | 0.39 (0.051) | 120 | -7.6 (0.77) | 98 | - | 86 | - | -0.7 | -3.59 (0.63) |
| k-410 | 0.33 (0.060) | 106 | - | 89 | -6.1 (0.99) | 86 | -5.6 (0.94) | 0.4 | - |

**Table S1. Ligand binding and cellular activity**
*Relative binding affinity E2 = 100

<table>
<tr><th></th><th>Chemical Structure</th><th>Proliferation Curves (WT, D538G, Y537S)</th><th>Luciferase Curves</th><th>(2Fo-Fc)[1] and (Fo-Fc)[2] difference maps</th></tr>
<tr><td>E2</td><td>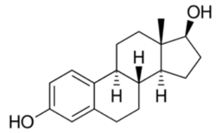
</td><td>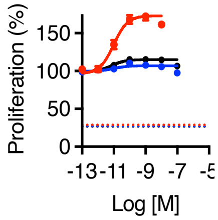
</td><td>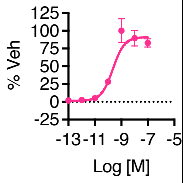
</td><td></td></tr>
<tr><td>4OHT</td><td>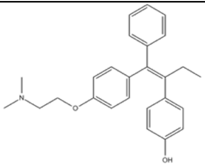</td><td>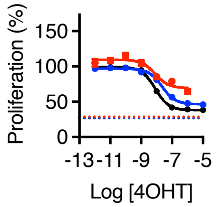
</td><td>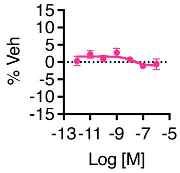
</td><td></td></tr>
<tr><td>Fulv</td><td>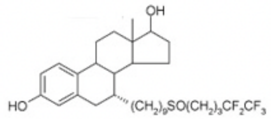
</td><td>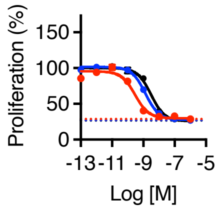
</td><td>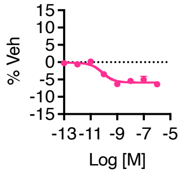
</td><td></td></tr>
<tr><td>k-402</td><td>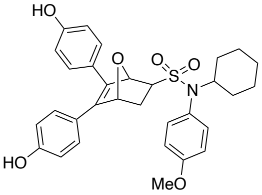
</td><td>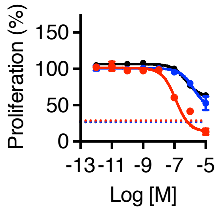
</td><td>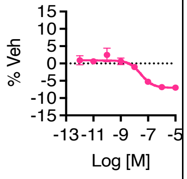
</td><td>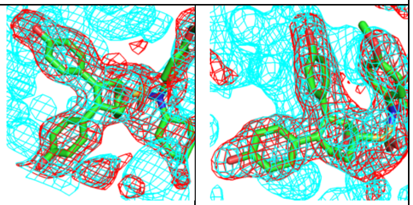</td></tr>
<tr><td>k-400</td><td>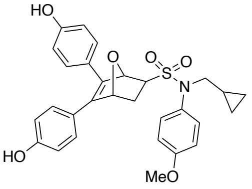
</td><td>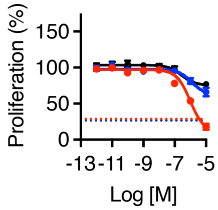
</td><td>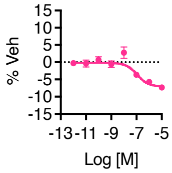
</td><td>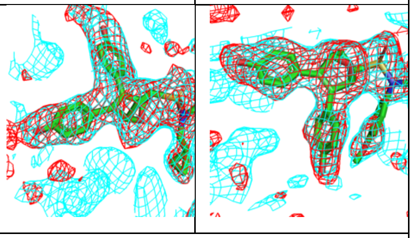</td></tr>
<tr><td>k-406</td><td>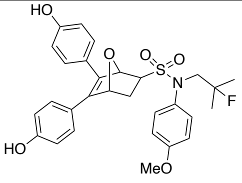
</td><td>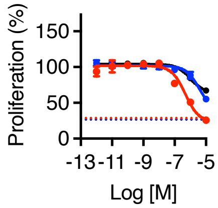
</td><td>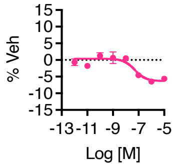
</td><td>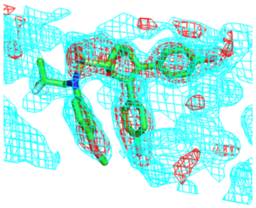</td></tr>
</table>

**Table S2. Compound structures and dose response curves.**

1. All (Fo-Fc) maps were generated after refinement of protein in absence of ligand. Maps are colored red and trimmed to 2.5Å RMSD of the ligand.
2. Refined protein-ligand and final corresponding (2Fo-Fc) maps are contoured at 0.7Å RMSD and colored cyan.

| | Chemical Structure | Proliferation Curves (WT, D538G, Y537S) | Luciferase Curves |
|---|---|---|---|
| k-424 | 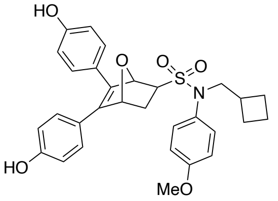  | 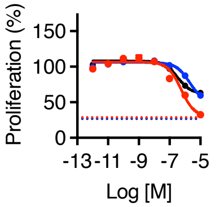  | 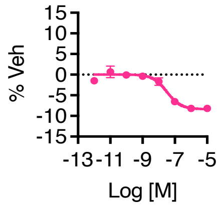  |
| k-422 | 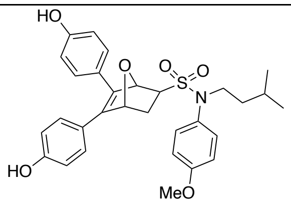  | 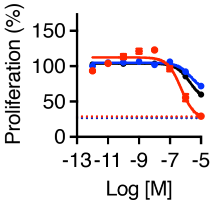  | 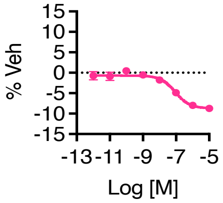  |
| k-4 | 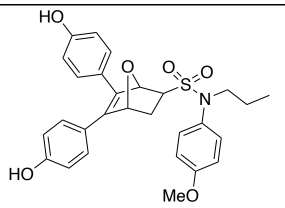  | 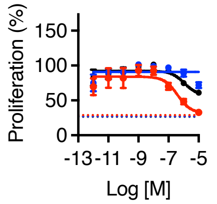  | 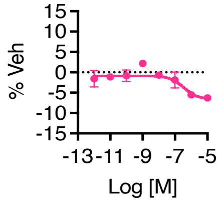  |
| k-5 | 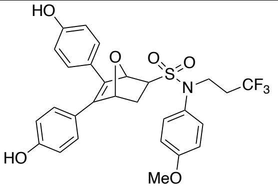  | 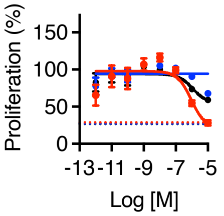  | 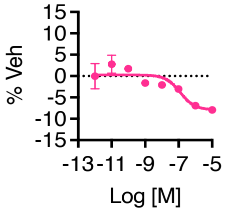  |
| k-408 | 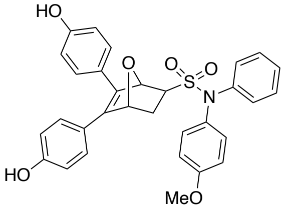  | 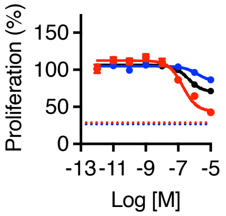  | 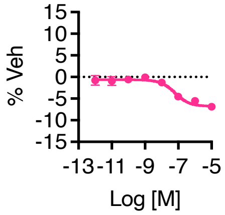  |
| k-11-67 | 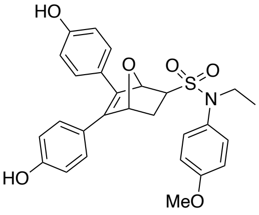  | 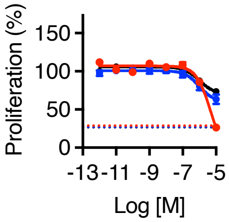  | 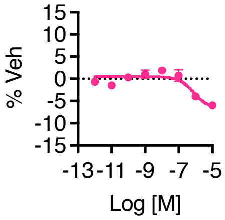  |

**Table S2. Compound structures and dose response curves.**

| | Chemical Structure | Proliferation Curves (WT, **D538G**, Y537S) | Luciferase Curves | $(2Fo\text{-}Fc)^1$ and $(Fo\text{-}Fc)^2$ difference maps |
|---|---|---|---|---|
| k-403 | 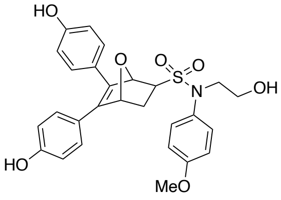 | 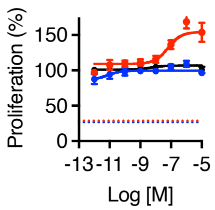 | 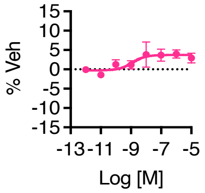 | 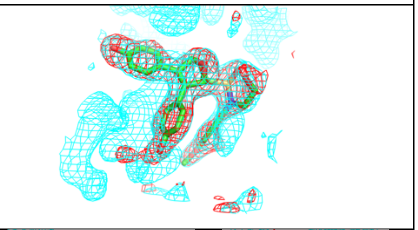 |
| k-411 | 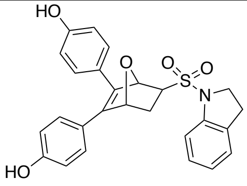 | 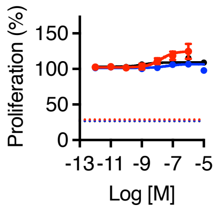 | 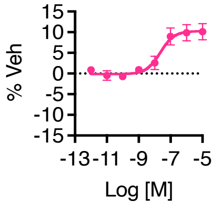 | 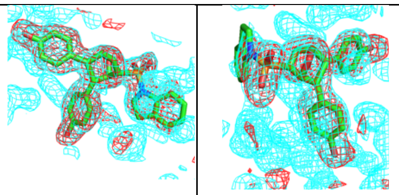 |
| k-409 | 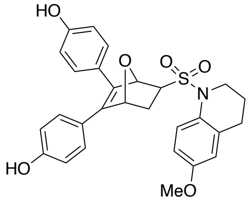 | 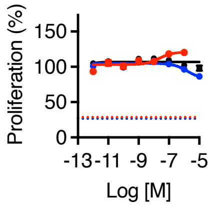 | 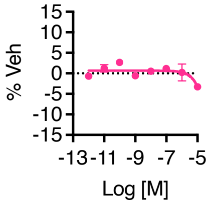 | 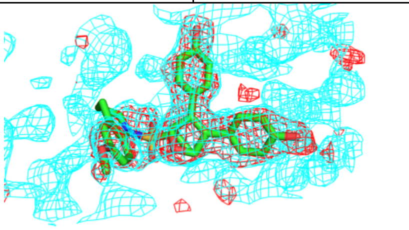 |
| k-410 | 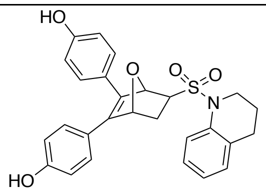 | 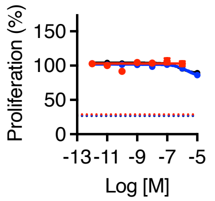 | 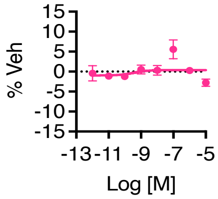 | 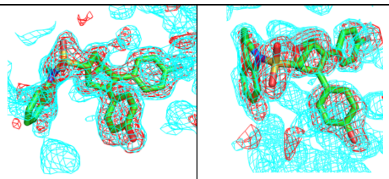 |

**Table S2. Compound structures and dose response curves.**

1. All (Fo-Fc) maps were generated after refinement of protein in absence of ligand. Maps are colored red and trimmed to 2.5Å RMSD of the ligand.
2. Refined protein-ligand and final corresponding (2Fo-Fc) maps are contoured at 0.7Å RMSD and colored cyan.

| Crystal name | BSK400_A2A3i | BSK402_A12e | BSK403_F10a | BSK406_A11f |
|---|---|---|---|---|
| **Compound #** | k-400 | k-402 | k-403 | k-406 |
| **PDB** | 8VZ0 | 8W07 | 8VZP | 8VZQ |
| **Data collection** | | | | |
| Wavelength (Å) | 1 | 1 | 1 | 1 |
| Space group | P 1 | P 1 | P 1 | P 1 |
| Cell dimensions | | | | |
| *a*, *b*, *c* (Å) | 53.259 58.886 93.0939 | 53.583 59.027 93.754 | 53.558 58.906 93.244 | 53.173 58.694 93.528 |
| *α*, *β*, *γ* (°) | 80.10 75.02 63.17 | 86.59 75.01 63.14 | 86.76 74.95 63.05 | 86.80 75.08 63.25 |
| Resolution (Å) | 33.052-1.860 (1.926-1.860) | 37.538-1.832 (1.897-1.832) | 37.204-1.716 (1.778-1.716) | 33.024-1.751 (1.814-1.751) |
| *R*merge | 0.284 | 0.12 | 0.171 | 0.121 |
| $I / \sigma I$ | 6.1 (1.9) | 9.7 (1.8) | 8.2 (1.7) | 9.7 (1.8) |
| Completeness (%) | 59.55 (2.53) | 71.40 (4.79) | 59.33 (2.47) | 62.95 (0.99) |
| Redundancy | 7.6 (7.6) | 7.8 (7.7) | 7.7 (7.6) | 7.7 (7.6) |
| CC(1/2) | 0.782 | 0.815 | 0.813 | 0.834 |
| **Refinement** | | | | |
| Resolution (Å) | 33.052-1.860 (1.926-1.860) | 37.538-1.832 (1.897-1.832) | 37.204-1.716 (1.778-1.716) | 33.024-1.751 (1.814-1.751) |
| No. reflections | 48681 (206) | 62018 (418) | 62076 (260) | 61711 (97) |
| *R*work / *R*free | 0.2063 / 0.2488 | 0.1787 / 0.2176 | 0.1809 / 0.2217 | 0.1825 / 0.2223 |
| No. atoms | | | | |
| Protein | 7011 | 7208 | 7103 | 7119 |
| Ligand/ion | 133 | 156 | 139 | 136 |
| Water | 168 | 330 | 295 | 427 |
| *B*-factors | | | | |
| Protein | 44.73 | 42.97 | 40.26 | 37.73 |
| Ligand/ion | 50.9 | 48.93 | 39.69 | 44.53 |
| Water | 44.81 | 43.12 | 40.3 | 38.1 |
| R.m.s. deviations | | | | |
| Bond lengths (Å) | 0.012 | 0.011 | 0.011 | 0.01 |
| Bond angles (°) | 1.97 | 1.37 | 1.77 | 0.912 |
| Clashscore, all atoms | 11.11 | 9 | 8.28 | 7.03 |
| Ramachandran outliers (%) | 0.23 | 0.34 | 0.11 | 0.34 |
| Ramachandran favored (%) | 97.74 | 98.54 | 98.7 | 98.77 |
| Side chain outliers (%) | 0.13 | 0 | 0 | 0 |

**Table S3. X-ray crystallography statistics**

*Highest-resolution shell is shown in parentheses.

| **Crystal name** | BSK409_E2c | BSK410_E10E11b | BSK411_F10c | KWN1154 |
|---|---|---|---|---|
| **Compound #** | k-409 | k-410 | k-411 | k-1154 |
| **PDB ID** | 8VZ1 | 8VYX | 8VYT | 8W03 |
| **Data collection** | | | | |
| Wavelength (Å) | 1 | 1 | 1 | 1 |
| Space group | P 1 | P 1 | P 1 | P1 |
| Cell dimensions | | | | |
| *a*, *b*, *c* (Å) | 53.529 58.971 93.745 | 53.809 58.867 94.127 | 53.394 58.696 93.356 | 53.489 58.771 93.208 |
| *α*, *β*, *γ* (°) | 86.51 75.05 63.02 | 86.75 74.98 62.91 | 86.88 74.94 63.05 | 86.607 75.165 62.959 |
| Resolution (Å) | 35.394-1.819 (1.884-1.819) | 33.101-1.692 (1.753-1.692) | 37.102-1.613 (1.671-1.613) | 38.359-1.681 (1.741-1.681) |
| *R*merge | 0.173 | 0.349 | 0.116 | 0.1 |
| *I* / σ*I* | 6.3 (1.7) | 6.2 (1.8) | 11.2 (1.7) | 11.4 (1.6) |
| Completeness (%) | 66.81 (3.61) | 68.28 (6.01) | 61.88 (4.09) | 59.79 (2.35) |
| Redundancy | 7.7 (7.6) | 7.1 (7.1) | 7.7 (7.7) | 7.3 (7.7) |
| CC(1/2) | 0.796 | 0.837 | 0.846 | 0.812 |
| **Refinement** | | | | |
| Resolution (Å) | 35.394-1.819 (1.884-1.819) | 33.101-1.692 (1.753-1.692) | 37.102-1.613 (1.671-1.613) | 38.359-1.681 (1.741-1.681) |
| No. reflections | 59112 (319) | 75444 (662) | 77528 (513) | 66313 (260) |
| *R*work / *R*free | 0.2248 / 0.2745 | 0.2086 / 0.2462 | 0.1796 / 0.2119 | 0.2016/0.2430 |
| No. atoms | | | | |
| Protein | 7161 | 7169 | 7291 | 7385 |
| Ligand/ion | 141 | 136 | 132 | 128 |
| Water | 298 | 541 | 608 | 426 |
| *B*-factors | | | | |
| Protein | 36.52 | 24.72 | 28.67 | 35.34 |
| Ligand/ion | 51.91 | 29.62 | 32.53 | 56.97 |
| Water | 36.8 | 25.18 | 29.39 | 35.81 |
| R.m.s. deviations | | | | |
| Bond lengths (Å) | 0.017 | 0.02 | 0.02 | 0.01 |
| Bond angles (°) | 1.83 | 2.08 | 2.04 | 1.99 |
| Clashscore, all atoms | 12.62 | 7.3 | 11.79 | 10.57 |
| Ramachandran outliers (%) | 0.34 | 0.11 | 0.11 | 0.11 |
| Ramachandran favored (%) | 97.19 | 98.87 | 98.2 | 98.23 |
| Side chain outliers (%) | 0 | 0 | 0 | 0.55 |

**Table S3. X-ray crystallography statistics**

*Highest-resolution shell is shown in parentheses.

| | Chain A | Chain B | Chain C | Chain D |
|---|---|---|---|---|
| k-400 | NA | down | out | out |
| k-402 | back | out | out | out |
| k-403 | out | out | out | out |
| k-406 | out | out | out | out |
| k-409 | down | down | down | down |
| k-410 | down | back | back | back |
| k-411 | down | back | down | down |
| **13** | NA | back | back | back |

**Table S4. Position of the F-ring relative to the ER structure.**

NA = poor electron density for the ligand

* The two substitutions from the nitrogen were cyclized into one R group

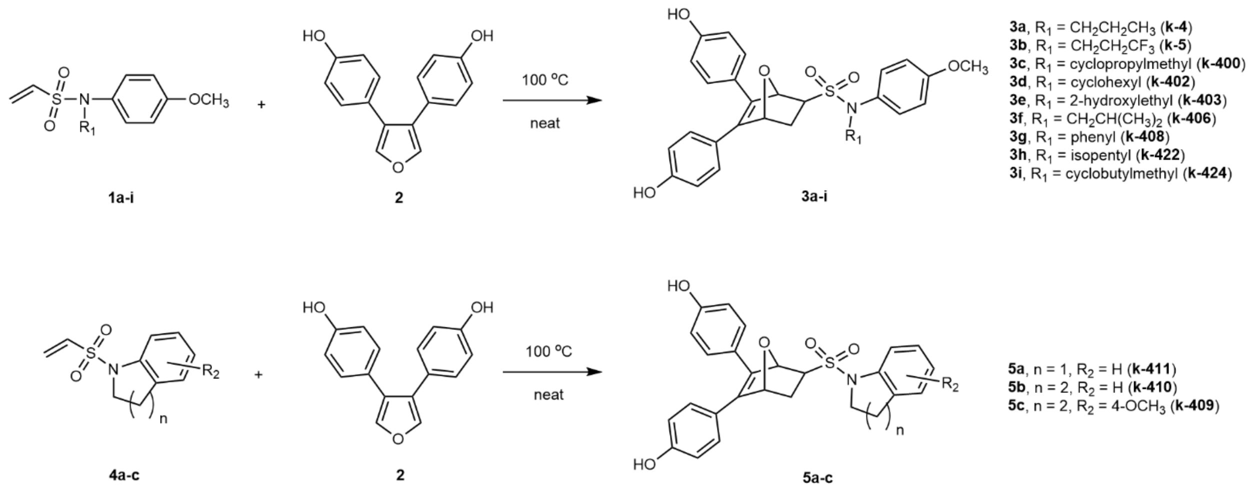

Scheme S1. Synthesis of OBHSN ligands

**Scheme S1. Chemical synthesis of OBHSN analogs**

### Materials and Methods

#### Cell culture

MCF7-ERα-Y537S and MCF7-ERα-D538G cells were a gift from Steffi Oesterreich, University of Pittsburgh Medical Center. Primary skeletal muscle stem cells were isolated from the leg muscles of 5-week-old female C57BL/6 mice and cultured as previously described (1). MCF7, MCF7-ERα-Y537S, and MCF7-ERα-D538G cells were maintained in Dulbecco's modified eagle medium (DMEM) supplemented with 10% fetal bovine serum (FBS) (Millipore Sigma, cat. no. F0926). All cell lines were cultured with 1X penicillin-streptomycin-neomycin antibiotics mixture (ThermoFisher Scientific, cat. no. 15640055), 1X MEM nonessential amino acids (ThermoFisher Scientific, cat no. 11140-050), 1X GlutaMAX (Gibco™ by Thermo Fisher Scientific, cat. no. 35050061) and 2.5 µg/ml plasmocin (Invivogen, cat. no. ant-mp). Mouse myoblast stem cells were differentiated to myotubes in steroid-free DMEM supplemented with insulin (Millipore-Sigma, cat. no. I9278) and 2% charcoal-stripped FBS. Myotube size was measured by high-content imaging as previously described(1). Cells were maintained at 37°C in a 5% $CO_2$ incubator.

#### Relative binding affinity

Radiolabeled E2 ([$^3$H]E2) ([6,7-$^3$H]estra-1,3,5, (10)-triene-3,17-β-diol), 54 Ci/mmol, was obtained from Amersham Pharmacia Biotech Biosciences (Piscataway, NJ). The RBA of the ligands was determined in wild-type ERα-LBD and baculovirus expressed full-length ERα (PanVera) by displacement of [$^3$H]E2 with unlabeled E2 or test compound and affinity expressed relative to E2.

#### Luciferase reporter assay

Steroid-deprived MCF7 cells were transfected with a 3xERE-luc reporter, treated with compounds and assayed for luciferase activity as previously described (2).

#### Cell Proliferation Assay

Cells were suspended in steroid-free media supplemented with 10% charcoal-stripped FBS and passed through a 30-micron strainer (Miltenyl Biotec, cat. no. 130-110-915). 1,000 cells in 25 µl of cell suspension were placed in each well of 384-well white, flat-bottom microplates (Greiner Bio-One CellStar, cat no. 781080). Plates were sealed with permeable membranes (Diversified

Biotech, Cat no. BEM-1), covered with a stainless-steel specimen plate lid (U.S. Patent 6,534,014), and incubated overnight at 37°C and 5% $CO_2$. The next day (i.e., Day-0), cells were treated with test compounds using a Biomek NXP 100-nl pintool (Beckman Coulter). The plates were sealed again, covered, and incubated at 37°C and 5% $CO_2$. After 5 days (i.e., Day-5), 25 µl of CellTiter-Glo® assay reagent (Promega, cat no. G7573) was added to each well. The plates were shaken gently for 5 min on an orbital shaker. Luminescence was measured using an Envision plate reader (PerkinElmer). Proliferation data was normalized using the initial luciferase activity (Day-0) as 0%, and the final luciferase activity in vehicle (DMSO)-treated wells (Day-5) as 100%.

**RNA isolation and real-time PCR**

Total RNA was isolated using TRIzol (Invitrogen) and reverse transcribed using MMTV reverse transcriptase (New England BioLabs). Real-time PCR was performed using SYBRgreen PCR Master Mix (Quantabio) as described(3). Relative mRNA levels of genes were normalized to the housekeeping gene 36B4, and fold-change calculated relative to the vehicle treated samples. Primer sequences for the genes studied were obtained from the Harvard Primer Bank. Sequences are available on their website. Primary mouse myotube cDNA samples were analyzed by real-time PCR using the PowerUp™ SYBR™ Green master mix (ThermoFisher, cat. no. A25742) with *Greb1* primers (Fwd, TCCTGCTGTACCTCTGTGACT; Rev, TGGCAGATCACACACAAGGT), and normalized to TaqMan® gene expression assay (ThermoFisher, cat. no. 4331182) reactions for *Akt1* (Mm01331626_m1). Results are the average ± SD from at least two independent experiments carried out in triplicate.

**Macromolecular x-ray crystallography**

The ERα-L372S/L536S mutant LBD was purified as previously described(4). The LBD (amino acid residues 298 through 554 with an N-terminal his-tag) was expressed in BL21 (DE3) Escherichia coli cells and purified by immobilized metal affinity chromatography using a $Ni^{2+}$ column, dialyzed after elution, and treated with Tobacco Etch Virus protease, and purified by ion exchange and size exclusion chromatography to remove the His-tag. The purified LBD was cocrystallized with ligands through sitting drop vapor diffusion method using trial gradients of 20 to 25% (weight/ volume) PEG 3350, 200 mM $MgCl_2$, and pH 6.5 through 8.0. Data were collected at the Stanford Synchrotron Radiation Lightsource (Beamline: 12–2) and Advanced Photon Source

(Beamlines: SERCAT BM22, ID-22), both at a temperature of 100 K and wavelength of 1.0 Å and scaled using AutoPROC (5) with the application of STARANISO (Globalphasing) to accommodate anisotropic diffraction. The structures were solved by molecular replacement from the starting raloxifene-bound ER LBD (PDB: 2QXS), and then refined using the PHENIX software suite version 1.20(6). Ligand restraint parameters were generated from PHENIX electronic Ligand Builder and Optimization Workbench(7). The LigandFit module in PHENIX was used to dock ligands within Crystallographic Object-Oriented Toolkit (COOT)(8). Refined structures were submitted to the PDB-REDO server before final refinement in PHENIX(9). Structures were analyzed using COOT and imaged using PyMOL (Schrodinger).

**Classical and accelerated molecular dynamics simulations**

The lasofoxifene/E2 dimer was generated by superposing 2OUZ.pdb onto the B chain of GWR.pdb using COOT(8). Missing portions of ER LBD were modeled in COOT by using other ER crystal structures with the same conformation trapping mutation and crystal packing environment. The Modeller46 extension (10) within UCSF Chimera (11) was used to fill in all other missing parts. Ligands were parameterized using Antechamber to generate force modification files using parmchk2 for use in tleap to generate topology and coordinate files using the AmberTools20 package. The ff14SB force field (12) was used to describe the ER LBD. The structures were solvated in a truncated octahedral box of TIP3P water molecules with a 12Å spacing between the protein and neutralized with $Na^+$ and $Cl^-$ ions. Minimization, heating, and equilibration of the protein-ligand complexes were performed as described previously(13). Constant pressure production runs were carried out at a temperature of 310K with Langevin dynamics using a collision frequency of $3ps^{-1}$. Monte Carlo barostat was used to control the pressure with a pressure relaxation time of 2ps. Hydrogen mass repartitioned (14) parameter files were used to carry out the production runs to allow 4fs time steps. 500ns production simulations were run in triplicates for each complex described.

For accelerated MD, a short 10ns classical MD simulation was used to compute average dihedral and potential energies as reference for the aMD parameters. A dual-boost approach was used by applying independent energy thresholds to enhance sampling(15). The aMD parameters were defined in terms of $E$ and $\alpha$, where $E$ represents the total boost energy level and $\alpha$ is a tuning parameter for the acceleration potential. $E_{total}$, $E_{dihedral}$, $\alpha_{total}$, and $\alpha_{dihedral}$ were calculated using the average $E_{total}$ and $E_{dihedral}$ from the 10ns classical MD simulation, as detailed in AMBER Advanced Tutorial 22 (https://ambermd.org/tutorials/advanced/tutorial22/).

**Simulation analysis**

Trajectory postprocessing was performed with CPPTRAJ(16). The autoimage command was used to recenter the trajectory coordinates and the strip command was used to remove water and ion molecules. The resulting trajectories were analyzed using CPPTRAJ and Bio3D(17). Distance distributions along the receptors (CPPTRAJ dist command) and receptor conformational heterogeneity (CPPTRAJ rmsd command) were measured. In Bio3D, the root mean square fluctuation (RMSF) was calculated by using the Cα's of each protein residue from every frame of the trajectories. Principal component analysis (PCA) was performed to understand the nature of the conformation differences of the trajectories by diagonalizing the fluctuation of the Cα's from the MD inputs. Bio3D's dccm function was used to generate a matrix of all atom-wise pairwise cross-correlation coefficients. Correlation matrices from different simulation inputs were subtracted to generate difference matrices. Correlation networks were constructed using Bio3D's cna() function. Correlation network analysis was carried out by selecting the Cα atoms of the protein residues and the C25 atom of the ligands to become the nodes for the network construction. Edges within the network were constructed based on Cα- Cα correlation values between all residues in the complex. A network edge was drawn if the pairwise correlation value was greater than 0.3 and the Cα- Cα distance was less than 10Å for at least 75% of the simulation. Suboptimal paths between designated "source" and "sink" nodes were drawn with the Bio3D cnapath() function, as previously described(18). The ligands on both sides of the ER-LBD dimer were designated as the source and sink nodes of the path construction. One hundred paths were collected for each pair of nodes that were analyzed. Suboptimal paths between residue sites were visualized as edges in VMD. Betweenness centrality was calculated with Bio3D.

**Potential binding energy calculation.**

The initial structures used in the calculations were built from our crystal structures and Protein Data Bank (PDB) 2QXS (Raloxifene-bound), 6VGH (Lasofoxifene-bound) and 3UUD (Estradiol-bound). Preparing of ligand-protein complex structures with protein preparation workflow in Schrodinger (Schrodinger, LLC, New York, NY). Splitting the structure by chain and splitting structure each chain by ligands, waters and protein. Calculating the potential binding energy of each monomer with MM-GBSA module in Schrodinger without flexible residues.

**RNA-seq**

MCF7 cells maintained in phenol red-free DMEM supplemented with 10% charcoal-stripped fetal bovine serum for 72 hours were treated with DMSO vehicle, 1 nM 17β-estradiol (E2), 100 nM fulvestrant, or a combination of E2 and fulvestrant. After 24 hours, total RNA was isolated using the RNeasy kit (QIAGEN) with on-column DNase digest. The experiment was repeated with 3 separate cell passages as biological replicates. RNA was quantified using the Qubit 2.0 Fluorometer (Invitrogen) and analyzed on the Agilent 4200 TapeStation RNA tape (Agilent Technologies) to assess quality. Only samples with RNA integrity number ≥ 9.5 were used. Messenger RNA (mRNA) was selected using the NEBNext poly(A) mRNA magnetic isolation module (Cat. #: E7490, New England Biolabs (NEB)) and libraries were prepared with NEBNext Ultra II Directional RNA kit (Cat. # E7760, NEB). The libraries were sequenced on the Illumina NextSeq 500 platform. Sequencing reads were further analyzed on the Galaxy web platform (v2.2.1) (19), including alignment to the human reference genome hg38 (female) using Hisat2 (v2.2.1) (20). Aligned reads were counted using featureCounts in the Subread (v2.0.1) package (21). Differential expression analysis was performed using DESeq2 (v1.36.0) (22). The RNA-seq dataset was deposited in the gene expression omnibus (GEO), with accession no. GSE231397.

### Chemistry

### Chemical Synthesis

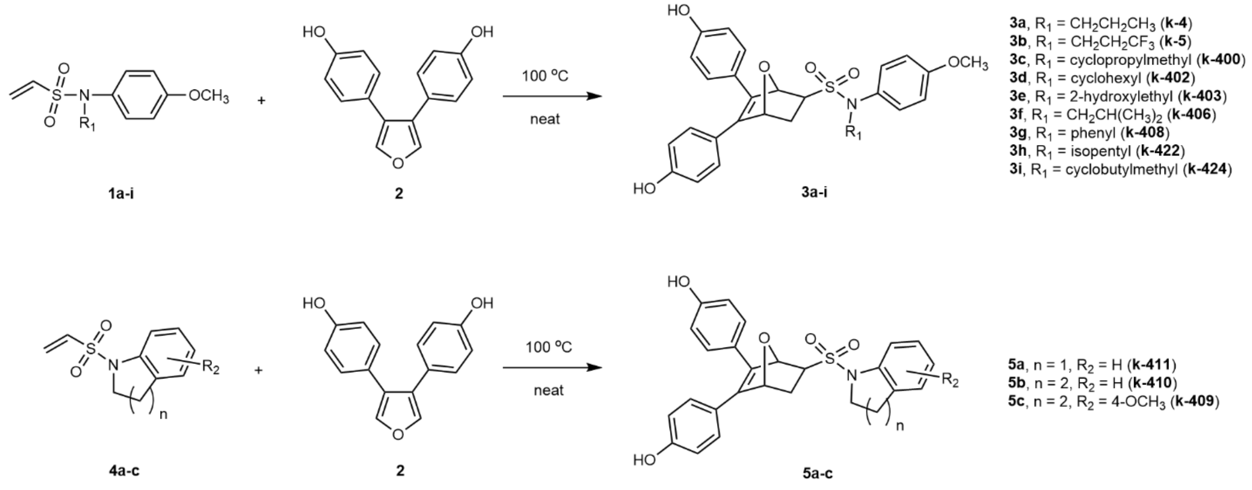

Scheme S1. Synthesis of OBHSN ligands

*General procedure for the Diels-Alder reaction to prepare OBHSN compounds.* The synthesis of OBHSN ligands is shown in Scheme S1. The Diels-Alder reaction of 3,4-bis(4-hydroxyphenyl)furan **2** (1.0 equiv) with corresponding dienophiles (1.2 equiv) gave the final OBHSN compounds directly. The Diels–Alder cycloaddition was generally run using ca. 0.5 mmol of furan and 0.6 mmol dienophile, and it proceeded well at 100 °C without any solvent or catalysts. The products were almost exclusively *exo* diastereomers. Purification was done by silica gel column with 1~3% MeOH in DCM, and the average yield of the purified *exo* product was ca. 50%.

The synthesis of k-**11-67** has been reported as compound **11j** in our previous paper(23).

*5,6-bis(4-hydroxyphenyl)-N-(4-methoxyphenyl)-N-propyl-7-oxabicyclo[2.2.1]hept-5-ene-2-sulfonamide (**k-4**).* Following the general procedure for Diels-Alder reaction between **1a** and **2**, compound **k-4** was obtained as yellow solid (Yield 59%). $^1$H NMR (500 MHz, acetone-$d_6$) δ 7.38 – 7.31 (m, 2H), 7.25 – 7.15 (m, 4H), 6.86 (s, 4H), 6.82 – 6.77 (m, 2H), 5.55 (s, 1H), 5.35 (d, J = 1.0 Hz, 1H), 4.52 (q, J = 8.6 Hz, 2H), 4.13 – 4.02 (m, 2H), 3.79 (s, 3H), 3.58 (s, 1H), 2.26 – 2.17 (m, 1H), 2.13 – 2.05 (m, 1H), 1.25 – 1.17 (m, 3H). $^{13}$C NMR (125 MHz, acetone-$d_6$) δ 159.67, 158.30, 157.83, 141.04, 138.39, 132.07, 130.61, 129.57, 128.65, 126.26, 123.58, 115.94, 114.95, 114.51, 84.55, 82.88, 68.19, 61.80, 55.10, 30.69, 30.32, 29.65, 29.50, 29.35, 29.19, 29.04, 28.88,

28.73, 20.17, 13.84. HRMS (*m/z*): [M + Na]$^+$ calcd. for $C_{28}H_{29}NO_6NaS$: 530.1613; found, 530.1620.

HPLC chromatogram report of **k-4**

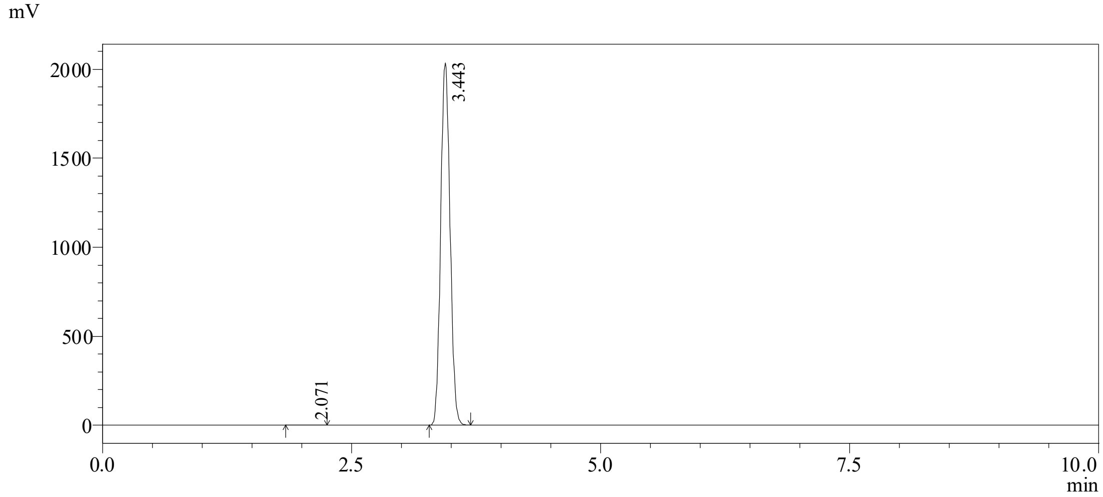

Peak list

| Peak# | RetTime [min] | Area [mV*s] | Height [mV] | Area % | Height % |
|---|---|---|---|---|---|
| 1 | 2.071 | 1307 | 186 | 0.010 | 0.010 |
| 2 | 3.443 | 13320578 | 2035422 | 99.990 | 99.990 |
| Total | | 13321885 | 2035608 | 100.000 | 100.000 |

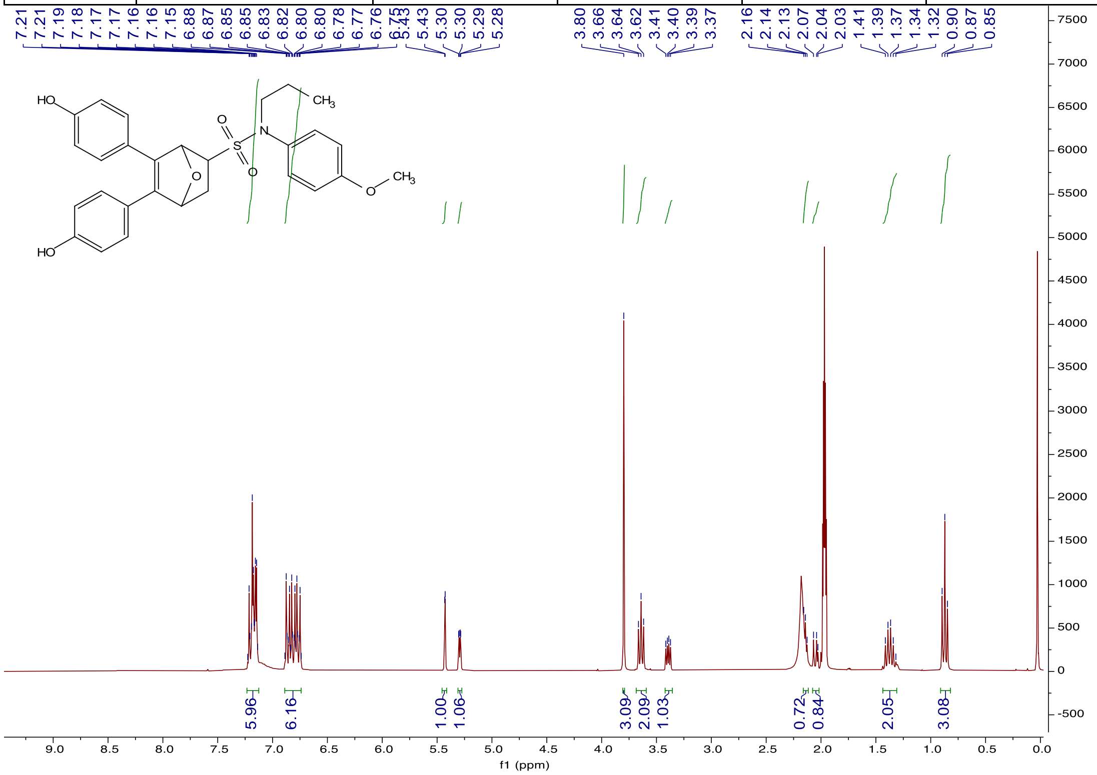

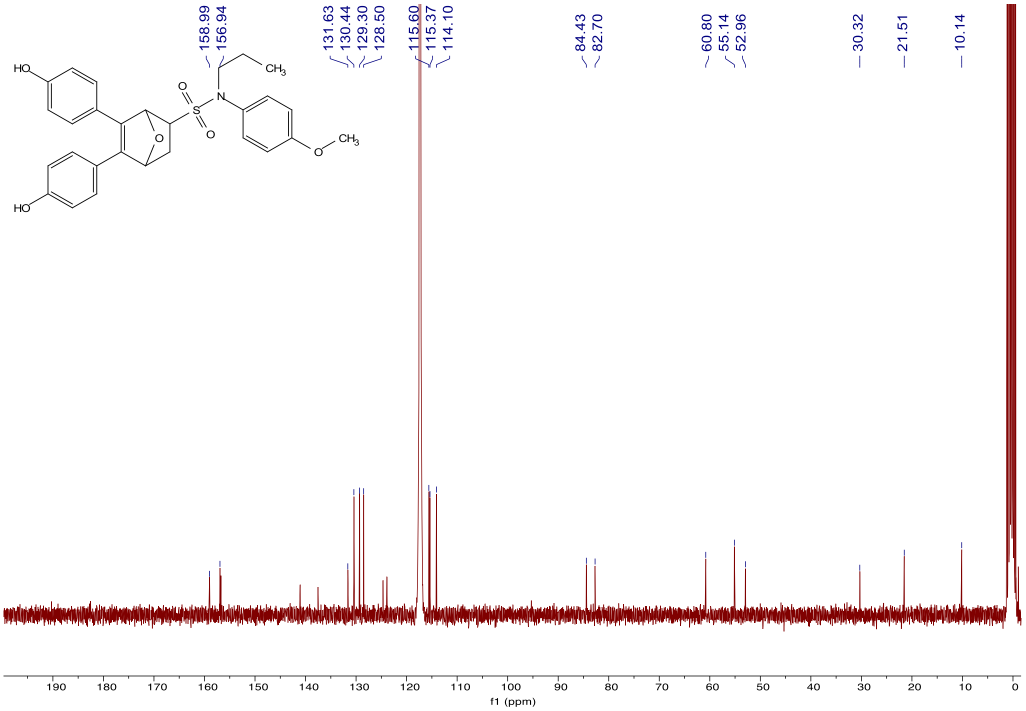

*5,6-bis(4-hydroxyphenyl)-N-(4-methoxyphenyl)-N-(3,3,3-trifluoropropyl)-7-oxabicyclo[2.2.1] hept-5-ene-2-sulfonamide (**k-5**)*. Following the general procedure for Diels-Alder reaction between **1b** and **2**, compound **k-5** was obtained as yellow solid (Yield 62%). $^{1}$H NMR (500 MHz, acetone-$d_6$) δ 7.34 (d, J = 8.9 Hz, 2H), 7.19 (t, J = 8.7 Hz, 4H), 6.85 (t, J = 9.1 Hz, 4H), 6.78 (d, J = 8.5 Hz, 2H), 5.52 (s, 1H), 5.33 (d, J = 4.4 Hz, 1H), 4.51 (q, J = 8.6 Hz, 2H), 3.79 (s, 3H), 3.55 (dd, J = 8.3, 4.4 Hz, 1H), 2.22 – 2.15 (m, 1H), 2.10 – 2.05 (m, 3H). $^{13}$C NMR (126 MHz, acetone-$d_6$) δ 159.67, 158.30, 157.83, 141.04, 138.39, 132.07, 130.61, 129.57, 128.65, 126.26, 123.58, 115.94, 114.95, 114.51, 84.55, 82.88, 68.19, 61.80, 55.10, 30.69, 30.32, 29.65, 29.50, 29.35, 29.19, 29.04, 28.88, 28.73, 20.17, 13.84. HRMS (*m/z*): $[M + Na]^+$ calcd. for $C_{28}H_{26}NO_6NaSF_3$: 584.1331; found, 584.1335.

HPLC chromatogram report of **k-5**

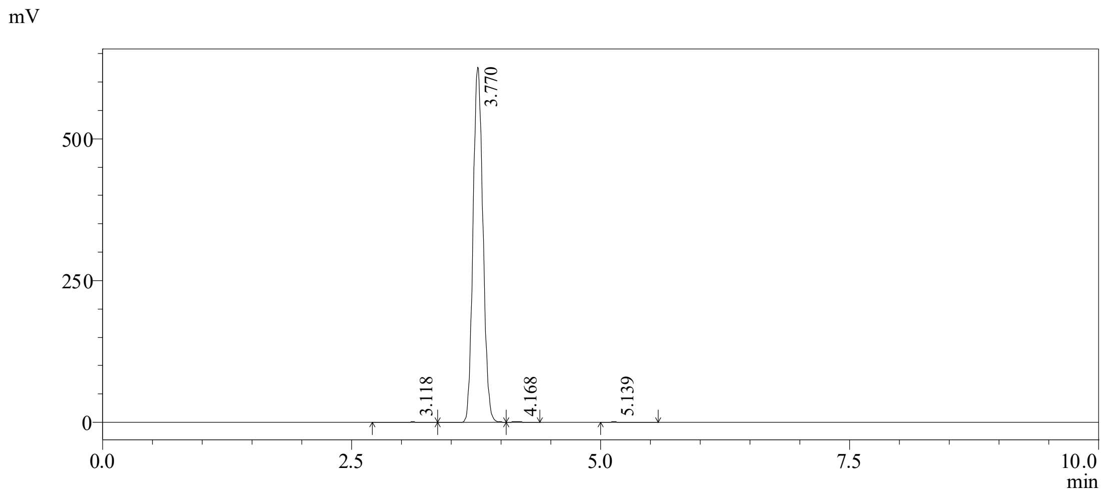

Peak list

| Peak# | RetTime [min] | Area [mV*s] | Height [mV] | Area % | Height % |
|---|---|---|---|---|---|
| 1 | 3.118 | 7371 | 661 | 0.169 | 0.169 |
| 2 | 3.770 | 4326651 | 626508 | 99.470 | 99.470 |
| 3 | 4.168 | 7834 | 735 | 0.180 | 0.180 |
| 4 | 5.139 | 7861 | 604 | 0.181 | 0.181 |
| Total |  | 4349717 | 628508 | 100.000 | 100.000 |

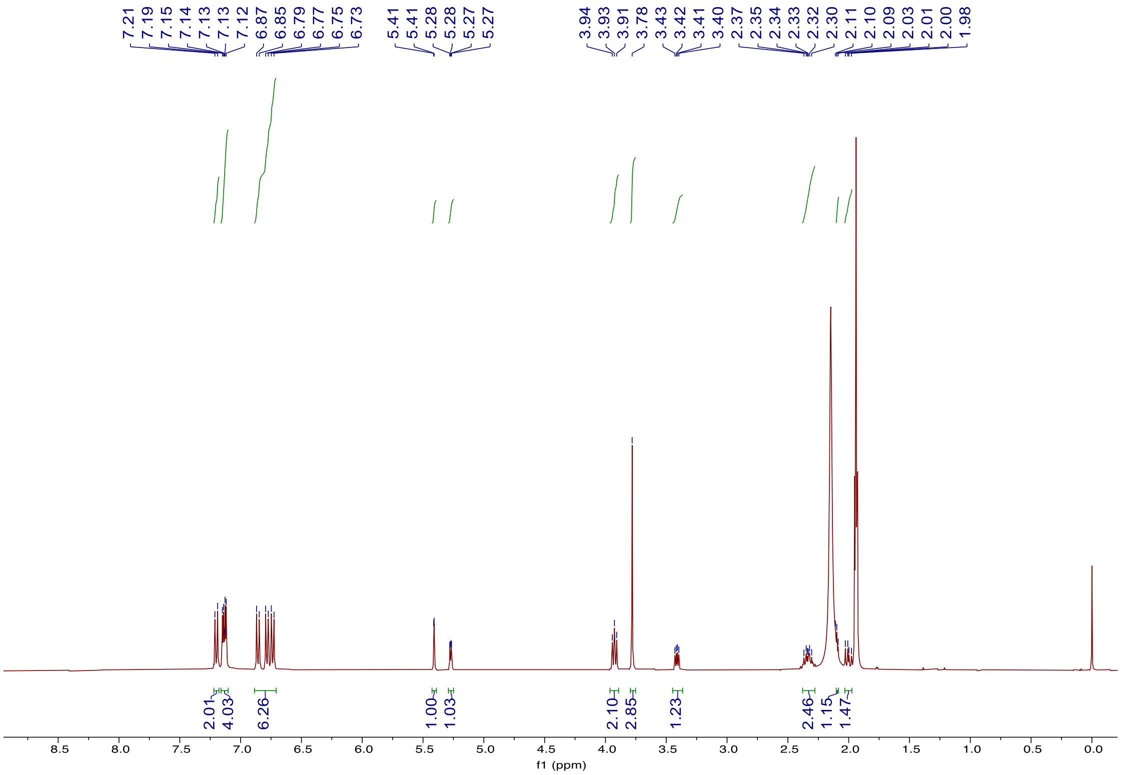

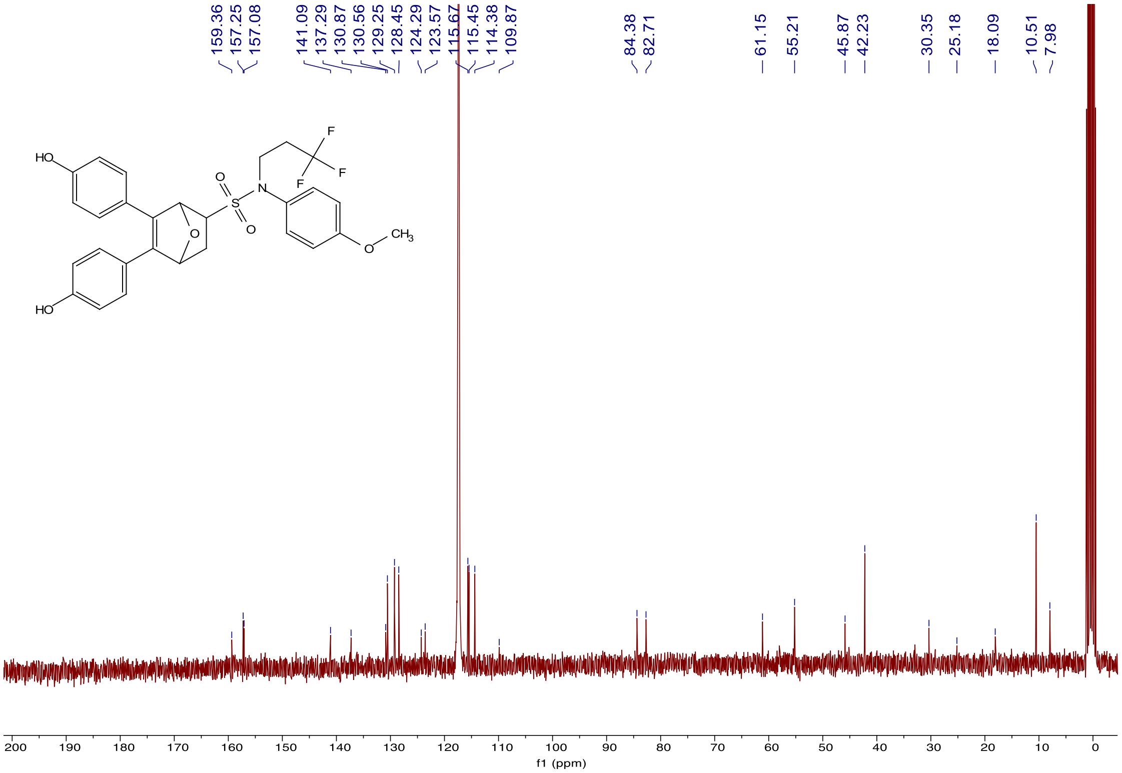

*N-(cyclopropylmethyl)-5,6-bis(4-hydroxyphenyl)-N-(4-methoxyphenyl)-7-oxabicyclo[2.2.1] hept-5-ene-2-sulfonamide (**k-400**)*. Following the general procedure for Diels-Alder reaction between **1c** and **2**, compound **k-400** was obtained as grey solid (Yield 32%). $^{1}H$ NMR (400 MHz, Acetonitrile-$d_3$) δ 7.12 – 7.09 (m, 4H), 7.05 – 7.01 (m, 2H), 6.83 – 6.77 (m, 2H), 6.75 – 6.68 (m, 4H), 5.39 (d, J = 1.2 Hz, 1H), 5.27 (dd, J = 4.5, 1.3 Hz, 1H), 3.74 (s, 3H), 3.30 (dd, J = 8.5, 4.5 Hz, 1H), 2.20 (t, J = 4.5 Hz, 1H), 1.99 – 1.96 (m, 1H). $^{13}C$ NMR (101 MHz, Acetonitrile-$d_3$) δ 159.64, 149.85, 132.84, 131.54, 129.88, 129.04, 116.16, 115.93, 114.60, 85.00, 83.27, 61.80, 57.14, 55.69, 30.84, 11.05, 3.79, 3.72. HRMS (m/z): $[M + Na]^+$ calcd for $C_{29}H_{29}NO_6SNa$, 542.1608; found, 542.1612.

HPLC chromatogram report of **k-400**

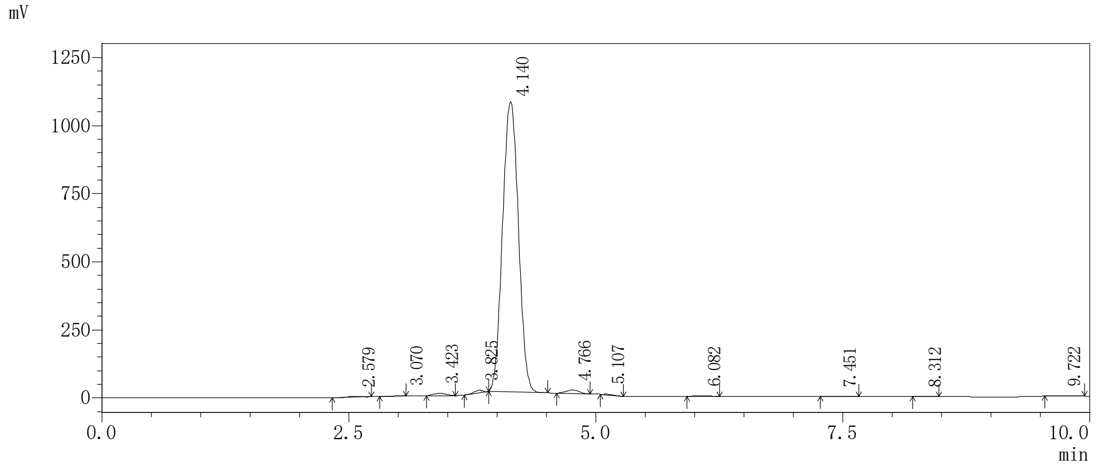

Peak list

| Peak# | RetTime [min] | Area [mV*s] | Height [mV] | Area % | Height % |
|---|---|---|---|---|---|
| 1 | 2.579 | 21795 | 3079 | 0.179 | 0.179 |
| 2 | 3.070 | -4783 | 215 | -0.039 | -0.039 |
| 3 | 3.423 | 76450 | 8096 | 0.627 | 0.627 |
| 4 | 3.825 | 50469 | 8391 | 0.414 | 0.414 |
| 5 | 4.140 | 11890672 | 1064624 | 97.461 | 97.461 |
| 6 | 4.766 | 115951 | 12303 | 0.950 | 0.950 |
| 7 | 5.107 | 10045 | 2106 | 0.082 | 0.082 |
| 8 | 6.082 | 13634 | 1208 | 0.112 | 0.112 |
| 9 | 7.451 | 9471 | 649 | 0.078 | 0.078 |
| 10 | 8.312 | 4449 | 455 | 0.036 | 0.036 |
| 11 | 9.722 | 12248 | 783 | 0.100 | 0.100 |
| Total | | 12200399 | 1101909 | 100.000 | 100.000 |

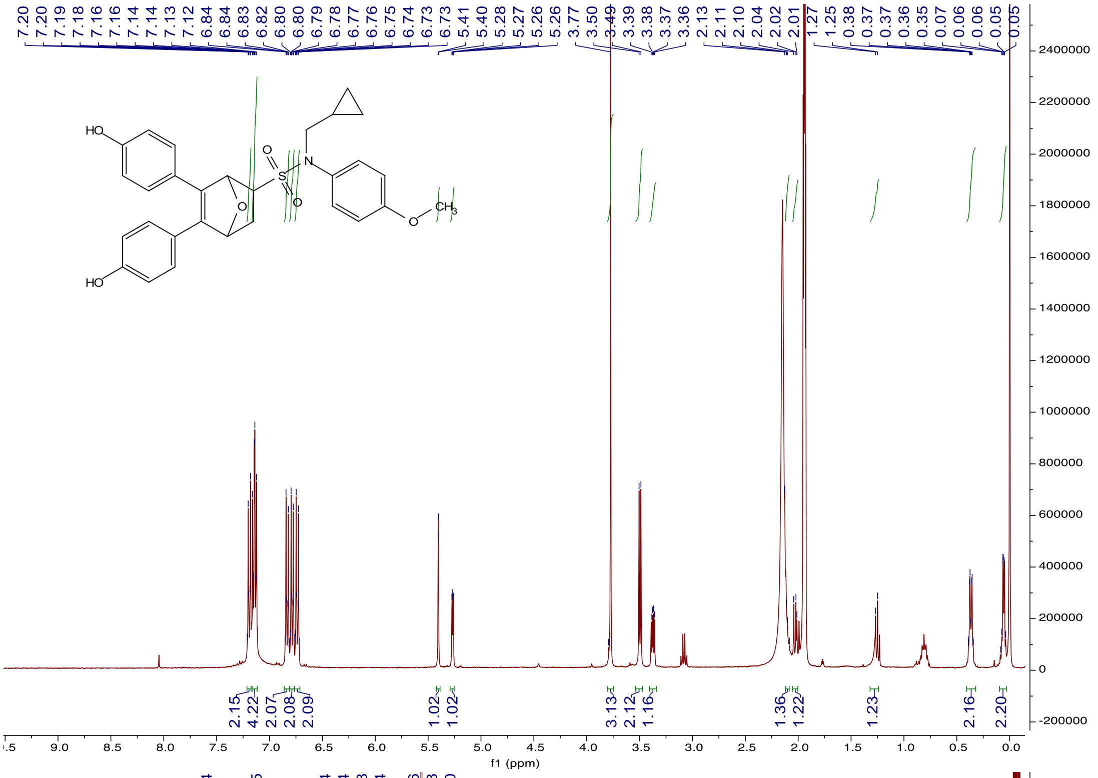
7.20 7.20 7.19 7.18 7.16 7.16 7.14 7.14 7.13 7.12 6.84 6.84 6.83 6.82 6.80 6.80 6.79 6.78 6.77 6.76 6.75 6.74 6.73 6.73 5.41 5.40 5.28 5.27 5.26 5.26 3.77 3.50 3.49 3.39 3.38 3.37 3.36 2.13 2.11 2.10 2.04 2.02 2.01 1.27 1.25 0.38 0.37 0.37 0.36 0.35 0.07 0.06 0.06 0.05 0.05
HO
HO
O
S
O
N
O
CH3
2.15 4.22 2.07 2.08 2.09 1.02 1.02 3.13 2.12 1.16 1.36 1.22 1.23 2.16 2.20
f1 (ppm)

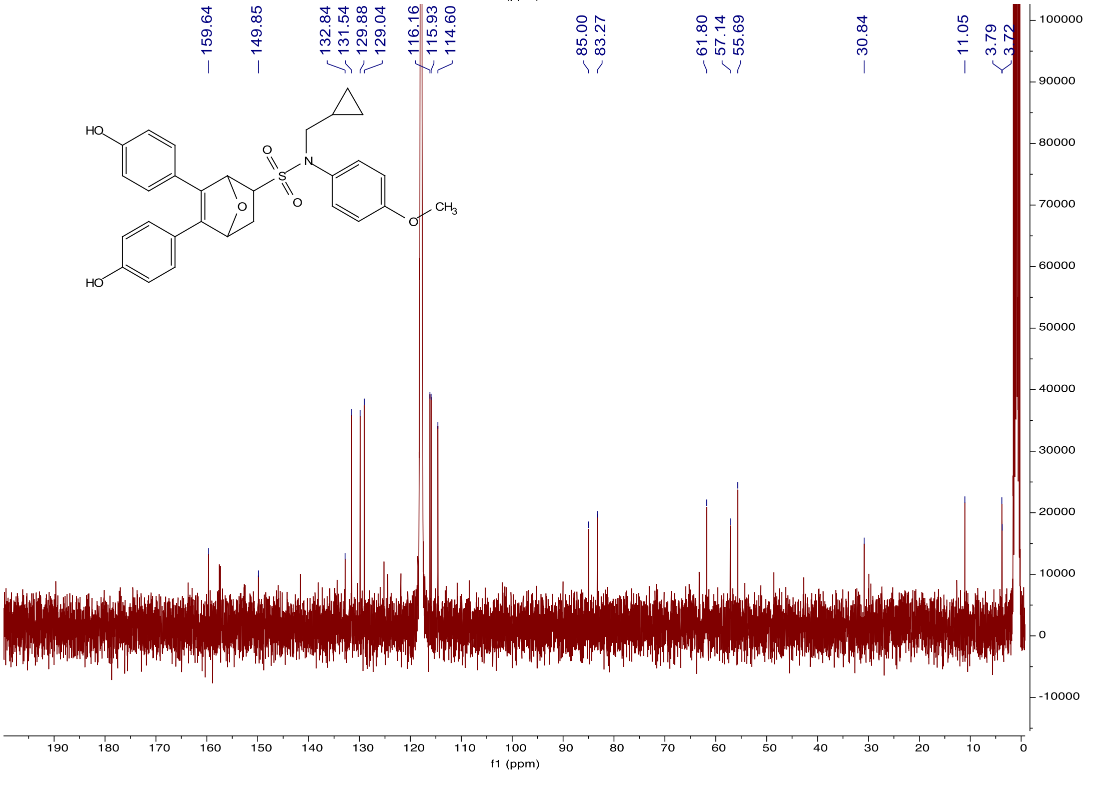
159.64 149.85 132.84 131.54 129.88 129.04 116.16 115.93 114.60 85.00 83.27 61.80 57.14 55.69 30.84 11.05 3.79 3.72
HO
HO
O
S
O
N
O
CH3
f1 (ppm)

*N-cyclohexyl-5,6-bis(4-hydroxyphenyl)-N-(4-methoxyphenyl)-7-oxabicyclo[2.2.1]hept-5-ene-2-sulfonamide (**k-402**)*. Following the general procedure for Diels-Alder reaction between **1d** and **2**, compound **k-402** was obtained as yellow solid (Yield 32%). $^{1}$H NMR (400 MHz, Acetone-$d_6$) δ 8.69 (s, 1H), 8.61 (s, 1H), 7.25 – 7.17 (m, 4H), 7.10 (d, J = 4.5 Hz, 2H), 6.83 (d, J = 6.7 Hz, 4H), 6.80 (s, 3H), 5.41 (s, 1H), 5.33 (d, J = 5.6 Hz, 1H), 3.79 (s, 3H), 3.61 – 3.54 (m, 1H), 3.43 (dd, J = 8.4, 4.7 Hz, 1H), 2.27 (dt, J = 11.9, 4.6 Hz, 1H), 2.12 (dd, J = 11.8, 8.3 Hz, 1H), 1.77 – 1.65 (m, 3H), 1.43 – 1.26 (m, 3H), 1.07 (dd, J = 9.6, 6.2 Hz, 2H), 0.92 – 0.78 (m, 2H). $^{13}$C NMR (101 MHz, Acetone-*$d_6$*) δ 159.58, 157.46, 157.24, 140.85, 137.76, 134.03, 129.25, 128.40, 124.51, 123.77, 115.68, 115.44, 113.59, 84.64, 82.68, 62.31, 59.00, 56.87, 54.86, 33.22, 30.43, 25.75, 24.97, 19.96, 18.00, 13.62. HRMS (m/z): $[M + Na]^+$ calcd for $C_{31}H_{33}NO_6SNa$, 570.1920; found, 570.1925.

HPLC chromatogram report of **k-402**

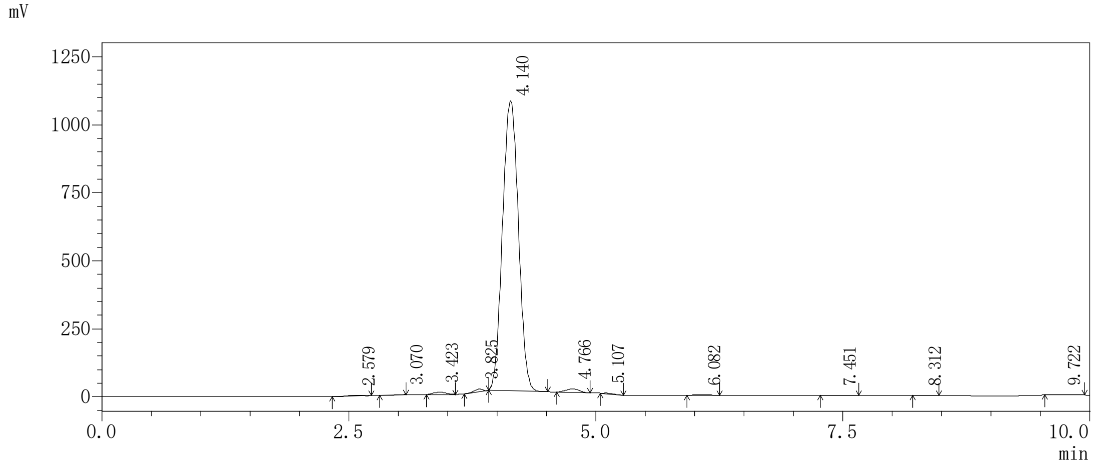

Peak list

| Peak# | RetTime [min] | Area [mV*s] | Height [mV] | Area % | Height % |
|---|---|---|---|---|---|
| 1 | 2.579 | 21795 | 3079 | 0.179 | 0.179 |
| 2 | 3.070 | -4783 | 215 | -0.039 | -0.039 |
| 3 | 3.423 | 76450 | 8096 | 0.627 | 0.627 |
| 4 | 3.825 | 50469 | 8391 | 0.414 | 0.414 |
| 5 | 4.140 | 11890672 | 1064624 | 97.461 | 97.461 |
| 6 | 4.766 | 115951 | 12303 | 0.950 | 0.950 |
| 7 | 5.107 | 10045 | 2106 | 0.082 | 0.082 |
| 8 | 6.082 | 13634 | 1208 | 0.112 | 0.112 |
| 9 | 7.451 | 9471 | 649 | 0.078 | 0.078 |
| 10 | 8.312 | 4449 | 455 | 0.036 | 0.036 |
| 11 | 9.722 | 12248 | 783 | 0.100 | 0.100 |
| Total | | 12200399 | 1101909 | 100.000 | 100.000 |

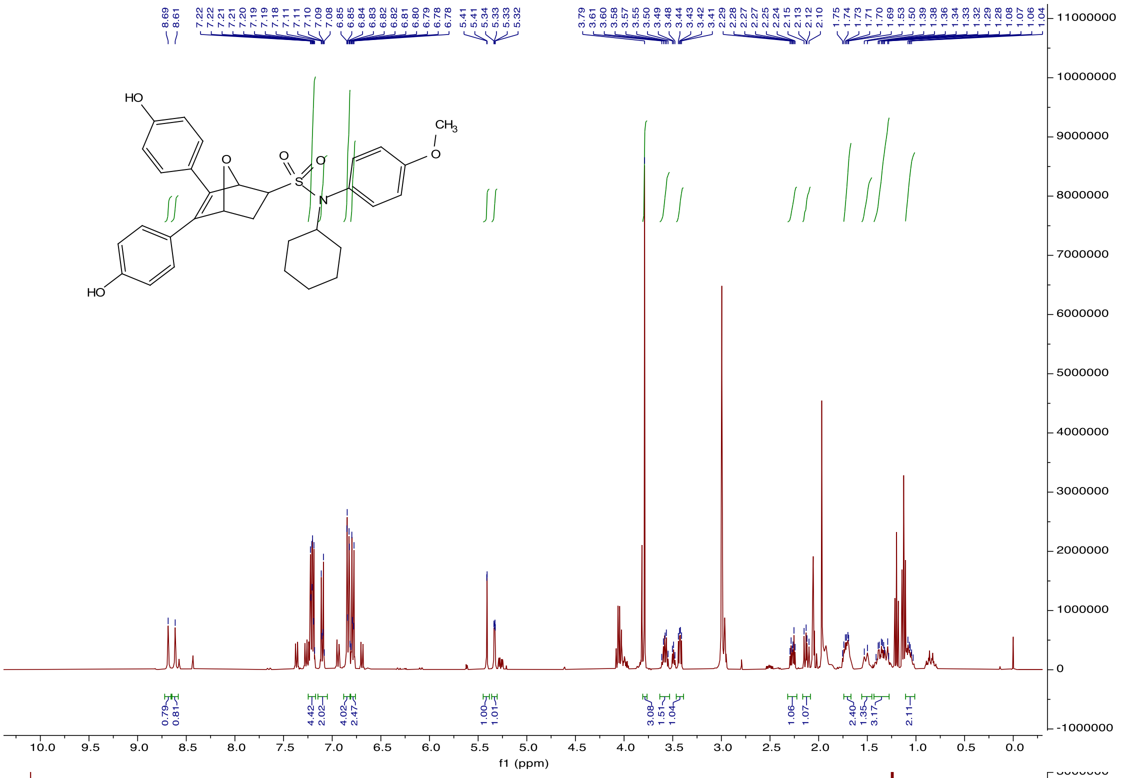

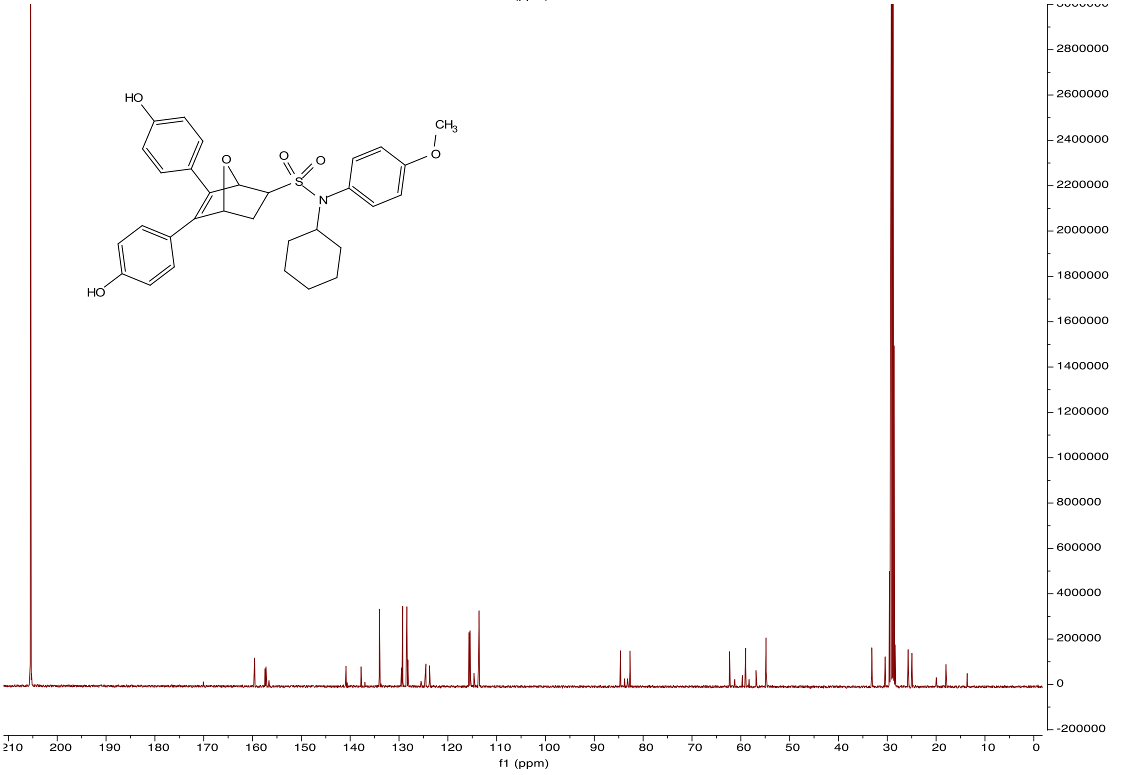

*N-(2-hydroxyethyl)-5,6-bis(4-hydroxyphenyl)-N-(4-methoxyphenyl)-7-oxabicyclo[2.2.1]hept-5-ene-2-sulfonamide (**k-403**)*. Following the general procedure for Diels-Alder reaction between **1e**

and **2**, compound **k-403** was obtained as yellow solid (Yield 21%). $^{1}$H NMR (501 MHz, Acetone-$d_6$) δ 8.65 (s, 1H), 8.59 (s, 1H), 7.29 (d, J = 8.7 Hz, 2H), 7.20 (t, J = 8.8 Hz, 4H), 6.84 (t, J = 8.5 Hz, 4H), 6.79 (d, J = 8.3 Hz, 2H), 5.51 (s, 1H), 5.31 (d, J = 3.5 Hz, 1H), 3.85 (dt, J = 12.8, 6.2 Hz, 1H), 3.79 (s, 5H), 3.54 (dq, J = 14.7, 4.3 Hz, 3H), 2.21 (dt, J = 11.3, 4.2 Hz, 1H). $^{13}$C NMR (101 MHz, Acetone-*$d_6$*) δ 205.35, 159.06, 157.40, 157.27, 140.94, 137.59, 132.31, 130.78, 129.14, 128.50, 124.45, 123.82, 115.64, 115.55, 115.45, 115.36, 114.03, 84.48, 82.68, 61.51, 59.47, 56.85, 54.83, 53.89, 30.51, 29.53, 29.34, 29.15, 28.96, 28.76, 28.57, 28.38, 18.00. HRMS (m/z): [M + Na]$^+$ calcd for $C_{27}H_{27}NO_7SNa$, 532.1400; found, 532.1404.

HPLC chromatogram report of **k-403**

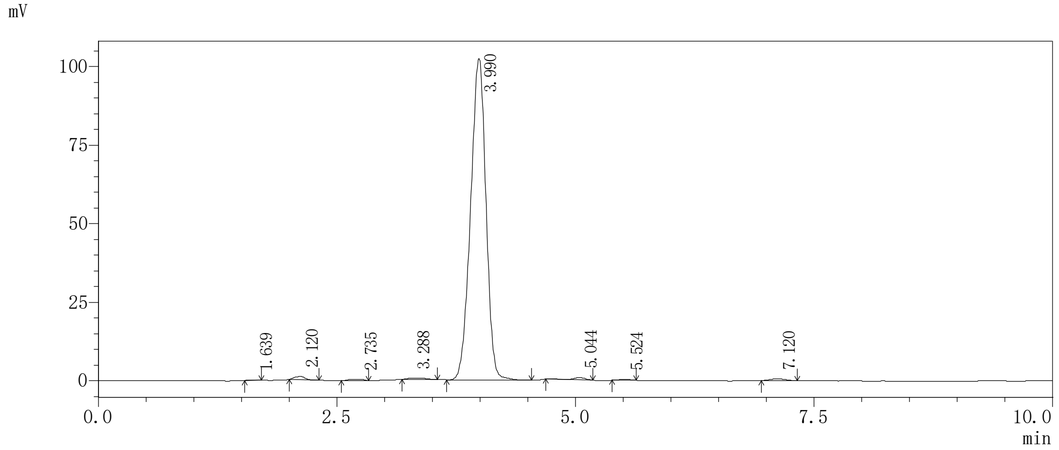

Peak list

| Peak# | RetTime [min] | Area [mV*s] | Height [mV] | Area % | Height % |
|---|---|---|---|---|---|
| 1 | 1.639 | 759 | 106 | 0.064 | 0.064 |
| 2 | 2.120 | 7647 | 1013 | 0.646 | 0.646 |
| 3 | 2.735 | 4742 | 444 | 0.401 | 0.401 |
| 4 | 3.288 | 4806 | 417 | 0.406 | 0.406 |
| 5 | 3.990 | 1149587 | 102377 | 97.162 | 97.162 |
| 6 | 5.044 | 6368 | 722 | 0.538 | 0.538 |
| 7 | 5.524 | 2450 | 264 | 0.207 | 0.207 |
| 8 | 7.120 | 6811 | 590 | 0.576 | 0.576 |
| Total | | 1183171 | 105933 | 100.000 | 100.000 |

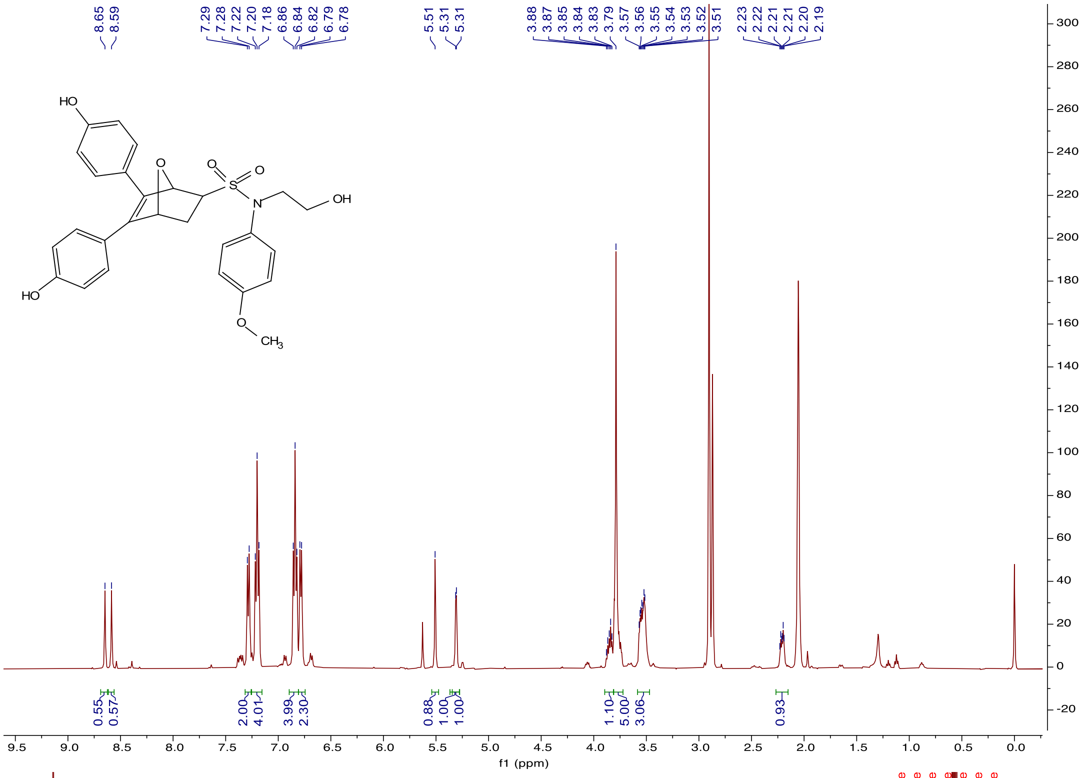
8.65
8.59
7.29
7.28
7.22
7.20
7.18
6.86
6.84
6.82
6.79
6.78
5.51
5.31
5.31
3.88
3.87
3.85
3.84
3.83
3.79
3.57
3.56
3.55
3.54
3.53
3.52
3.51
2.23
2.22
2.21
2.21
2.20
2.19
HO
O
O
S
O
N
OH
HO
O
CH3
0.55
0.57
2.00
4.01
3.99
2.30
0.88
1.00
1.00
1.10
5.00
3.06
0.93
f1 (ppm)

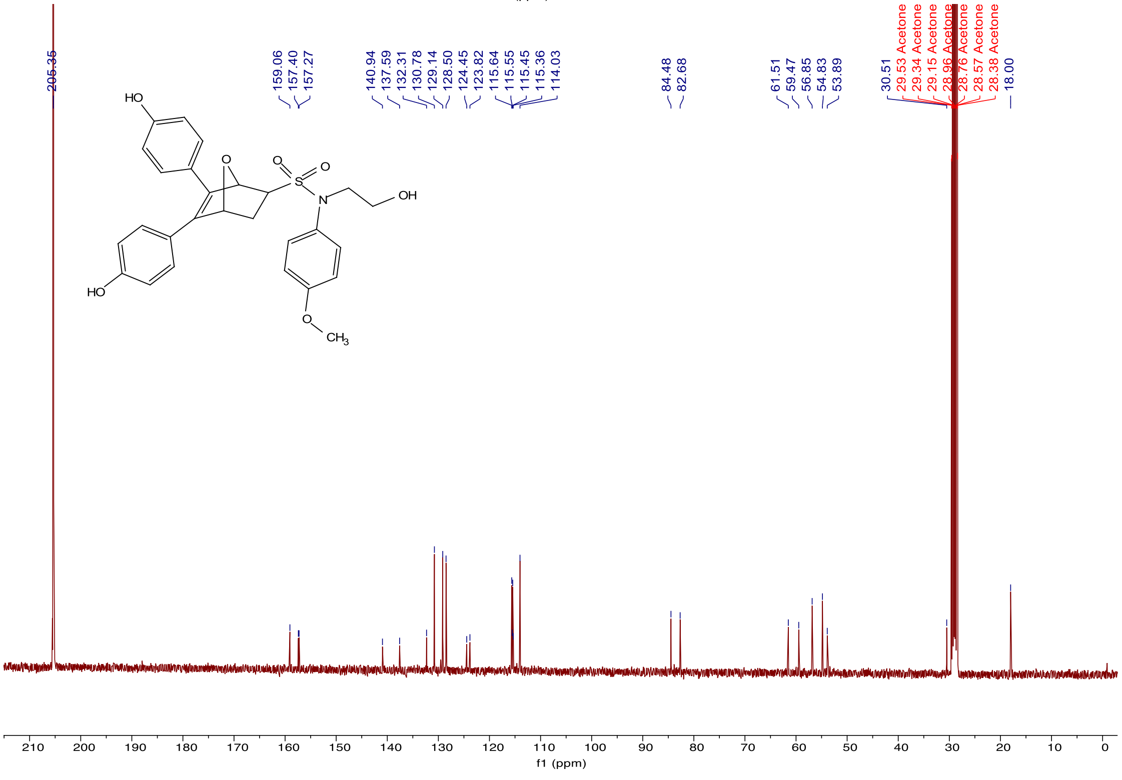
205.35
159.06
157.40
157.27
140.94
137.59
132.31
130.78
129.14
128.50
124.45
123.82
115.64
115.55
115.45
115.36
114.03
84.48
82.68
61.51
59.47
56.85
54.83
53.89
30.51
29.53 Acetone
29.34 Acetone
29.15 Acetone
28.96 Acetone
28.76 Acetone
28.57 Acetone
28.38 Acetone
18.00
HO
O
O
S
O
N
OH
HO
O
CH3
f1 (ppm)

*5,6-bis(4-hydroxyphenyl)-N-isobutyl-N-(4-methoxyphenyl)-7-oxabicyclo[2.2.1]hept-5-ene-2-sulfonamide (**k-406**)*. Following the general procedure for Diels-Alder reaction between **1f** and **2**, compound **k-406** was obtained as yellow solid (Yield 55%). $^1$H NMR (500 MHz, $CDCl_3$) δ 7.24 (d, J = 8.9 Hz, 2H), 7.13 (t, J = 8.4 Hz, 4H), 6.81 (d, J = 8.9 Hz, 2H), 6.75 (d, J = 7.3 Hz, 4H), 5.54 (s, 1H), 5.47 (d, J = 2.5 Hz, 1H), 3.79 (s, 3H), 3.55 (d, J = 7.3 Hz, 2H), 3.41 (dd, J = 8.4, 4.4 Hz, 1H), 2.35 – 2.26 (m, 1H), 2.00 – 1.90 (m, 1H), 1.57 (dt, J = 13.6, 6.7 Hz, 1H), 0.91 (dd, J = 10.5, 6.7 Hz, 6H). $^{13}$C NMR (126 MHz, $CDCl_3$) δ 159.88, 159.11, 155.79, 137.69, 131.74, 130.92, 129.27, 128.87, 125.33, 125.11, 124.41, 115.92, 115.06, 114.97, 84.62, 83.16, 77.52, 77.26, 77.01, 67.15, 62.60, 55.70, 33.63, 30.75, 29.66, 28.08. HRMS (m/z): $[M + Na]^+$ calcd. for $C_{29}H_{31}NO_6SNa$: 544.1770; found, 544.1763.

HPLC chromatogram report of **k-406**

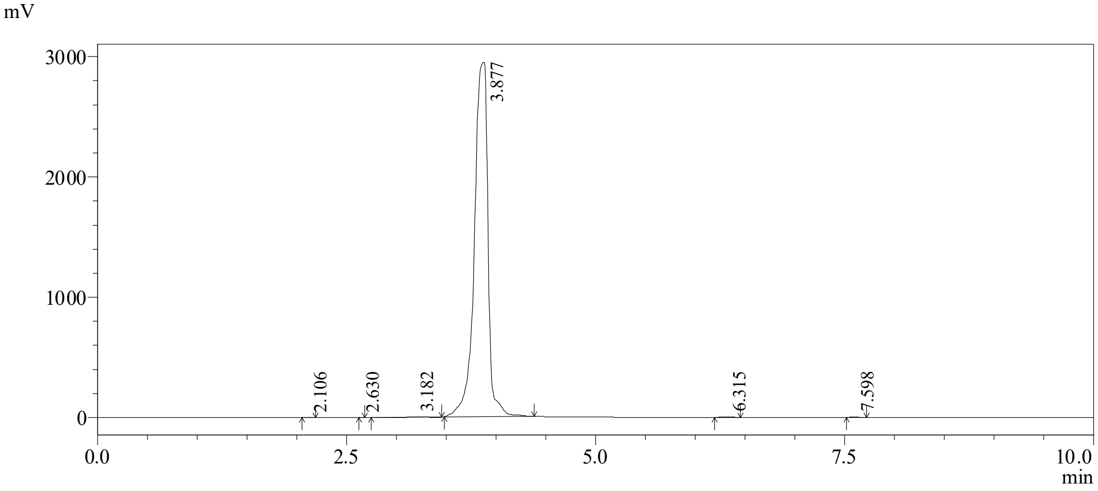

Peak list

| Peak# | RetTime [min] | Area [mV*s] | Height [mV] | Area % | Height % |
|---|---|---|---|---|---|
| 1 | 2.106 | 876 | 237 | 0.003 | 0.003 |
| 2 | 2.630 | 35 | 18 | 0.000 | 0.000 |
| 3 | 3.182 | 18757 | 2449 | 0.067 | 0.067 |
| 4 | 3.877 | 28008470 | 2943902 | 99.755 | 99.755 |
| 5 | 6.315 | 42398 | 5766 | 0.151 | 0.151 |
| 6 | 7.598 | 6805 | 950 | 0.024 | 0.024 |
| Total | | 28077341 | 2953321 | 100.000 | 100.000 |

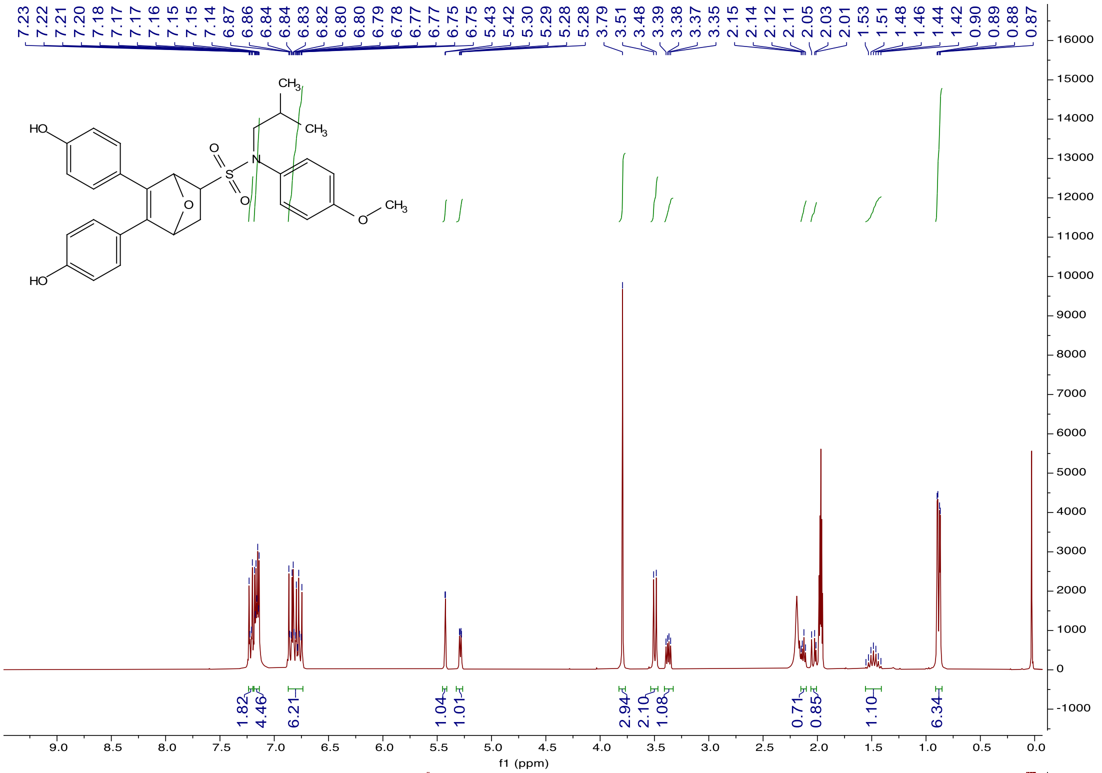
7.23 7.22 7.21 7.20 7.18 7.17 7.17 7.16 7.15 7.15 7.14 6.87 6.86 6.84 6.83 6.82 6.80 6.80 6.79 6.78 6.77 6.77 6.75 6.75 5.43 5.42 5.30 5.29 5.28 3.79 3.51 3.48 3.39 3.38 3.37 3.35 2.15 2.14 2.12 2.11 2.05 2.03 2.01 1.53 1.51 1.48 1.46 1.44 1.42 0.90 0.89 0.88 0.87
HO
HO
O
S
N
O
CH3
CH3
O
CH3
1.82
4.46
6.21
1.04
1.01
2.94
2.10
1.08
0.71
0.85
1.10
6.34
f1 (ppm)

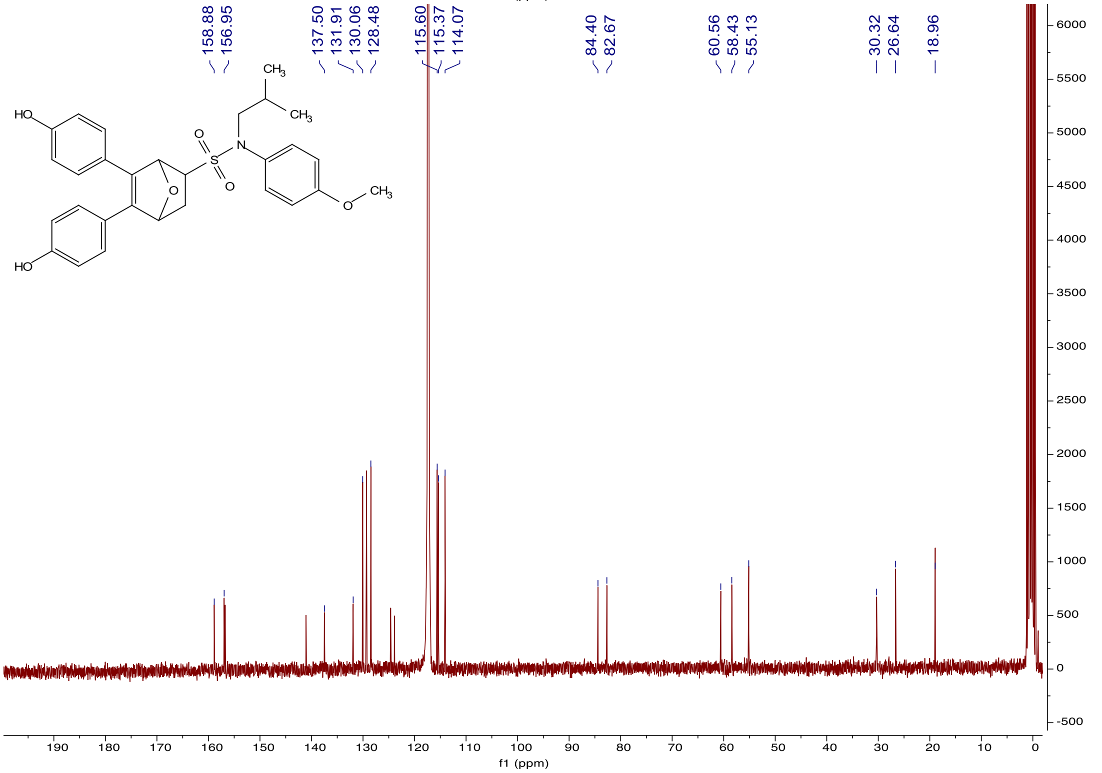
158.88 156.95 137.50 131.91 130.06 128.48 115.60 115.37 114.07 84.40 82.67 60.56 58.43 55.13 30.32 26.64 18.96
HO
HO
O
S
N
O
CH3
CH3
O
CH3
f1 (ppm)

*5,6-bis(4-hydroxyphenyl)-N-(4-methoxyphenyl)-N-phenyl-7-oxabicyclo[2.2.1]hept-5-ene-2-sulfonamide (**k-408**)*. Following the general procedure for Diels-Alder reaction between **1g** and **2**, compound **k-408** was obtained as white solid (Yield 51%). $^{1}$H NMR (400 MHz, Acetonitrile-d$_3$) δ 7.43 – 7.38 (m, 2H), 7.37 – 7.31 (m, 4H), 7.29 – 7.24 (m, 1H), 7.18 – 7.09 (m, 4H), 6.90 – 6.84 (m, 2H), 6.80 – 6.72 (m, 4H), 5.46 (d, J = 1.3 Hz, 1H), 5.29 (dd, J = 4.3, 1.3 Hz, 1H), 3.76 (s, 3H), 3.64 (dd, J = 8.3, 4.6 Hz, 1H), 2.25 – 2.22 (m, 1H), 2.10 (d, J = 2.6 Hz, 1H). $^{13}$C NMR (101 MHz, Acetonitrile-*d*$_3$) δ 157.53, 157.39, 142.75, 137.90, 134.61, 131.18, 129.97, 129.80, 129.10, 128.83, 127.92, 115.94, 115.09, 85.25, 83.29, 63.23, 55.75, 31.18. HRMS (m/z): [M + Na]$^+$ calcd for $C_{31}H_{27}NO_6SNa$, 541.1451; found, 564.1458.

HPLC chromatogram report of **k-408**

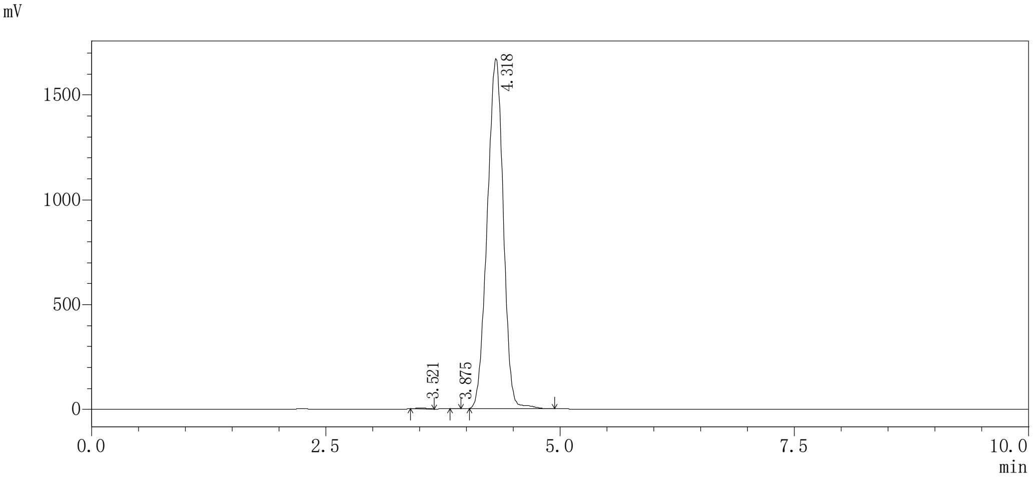

Peak list

| Peak# | RetTime [min] | Area [mV*s] | Height [mV] | Area % | Height % |
|---|---|---|---|---|---|
| 1 | 3.521 | 30181 | 3989 | 0.152 | 0.152 |
| 2 | 3.875 | 2196 | 490 | 0.011 | 0.011 |
| 3 | 4.318 | 19838217 | 1667262 | 99.837 | 99.837 |
| Total | | 19870594 | 1671740 | 100.000 | 100.000 |

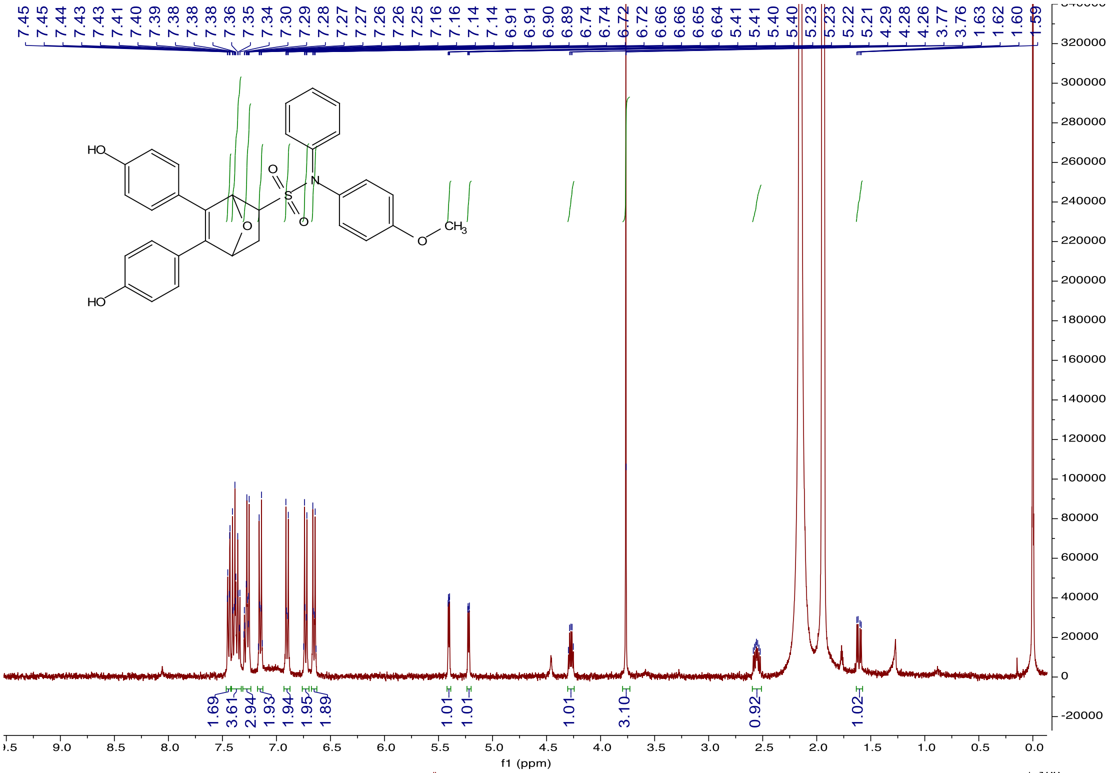

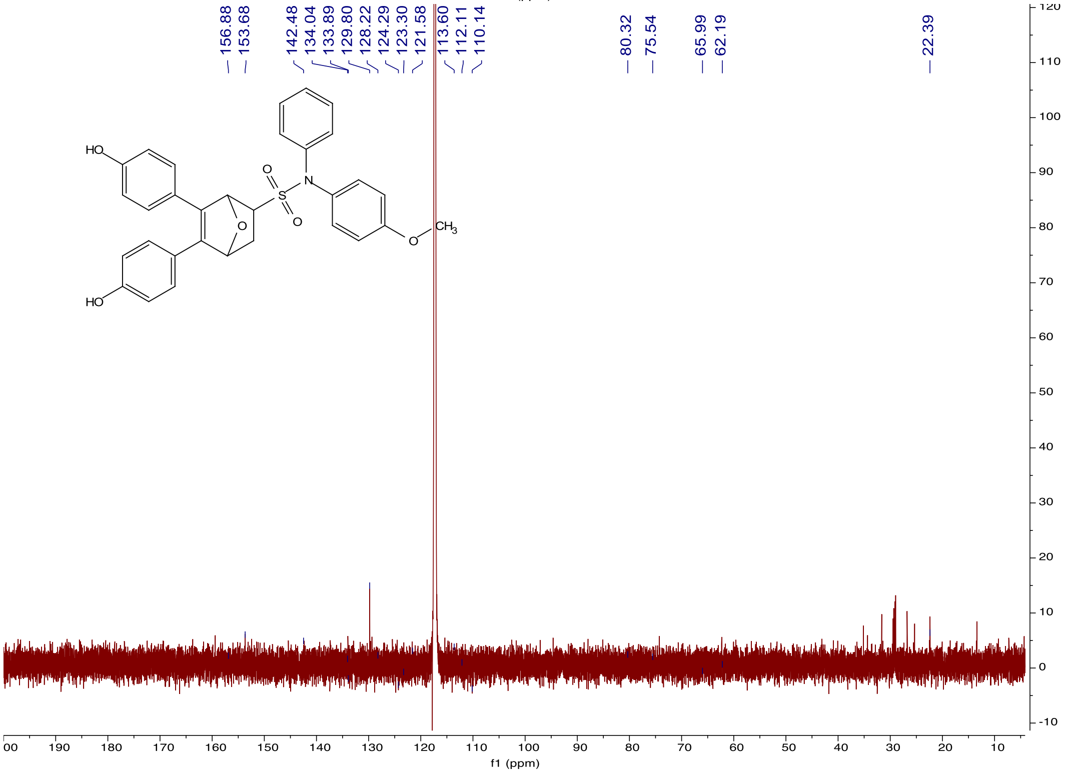

*4,4'-(5-((6-methoxy-3,4-dihydroquinolin-1(2H)-yl)sulfonyl)-7-oxabicyclo[2.2.1]hept-2-ene-2,3-diyl)diphenol (**k-409**).* Following the general procedure for Diels-Alder reaction between **4c** and **2**, compound **k-409** was obtained as yellow solid (Yield 45%). $^{1}$H NMR (400 MHz, Acetone-$d_6$) δ 8.67 (d, J = 16.8 Hz, 2H), 7.64 (d, J = 9.0 Hz, 1H), 7.20 – 7.12 (m, 2H), 7.06 – 6.97 (m, 2H), 6.80 – 6.72 (m, 5H), 6.69 (d, J = 3.0 Hz, 1H), 5.34 – 5.24 (m, 2H), 3.85 – 3.78 (m, 1H), 3.77 (s, 3H), 3.76 – 3.68 (m, 1H), 3.57 (dd, J = 8.4, 4.5 Hz, 1H), 2.71 (dtd, J = 23.3, 16.8, 6.7 Hz, 2H), 2.28 (dt, J = 11.8, 4.4 Hz, 1H), 2.03 – 1.97 (m, 1H), 1.89 (dtd, J = 13.9, 7.0, 4.8 Hz, 2H).$^{13}$C NMR (101 MHz, Acetone-*$d_6$*) δ157.31, 157.30, 156.57, 140.97, 137.27, 131.59, 130.42, 128.78, 128.63, 124.35, 124.28, 123.62, 115.58, 115.47, 114.03, 112.24, 84.55, 82.55, 61.94, 54.78, 54.73, 46.50, 30.38, 26.80, 22.45. HRMS (m/z): $[M + Na]^+$ calcd for $C_{28}H_{27}NO_6SNa$, 528.1451; found, 528.1461.

HPLC chromatogram report of **k-409**

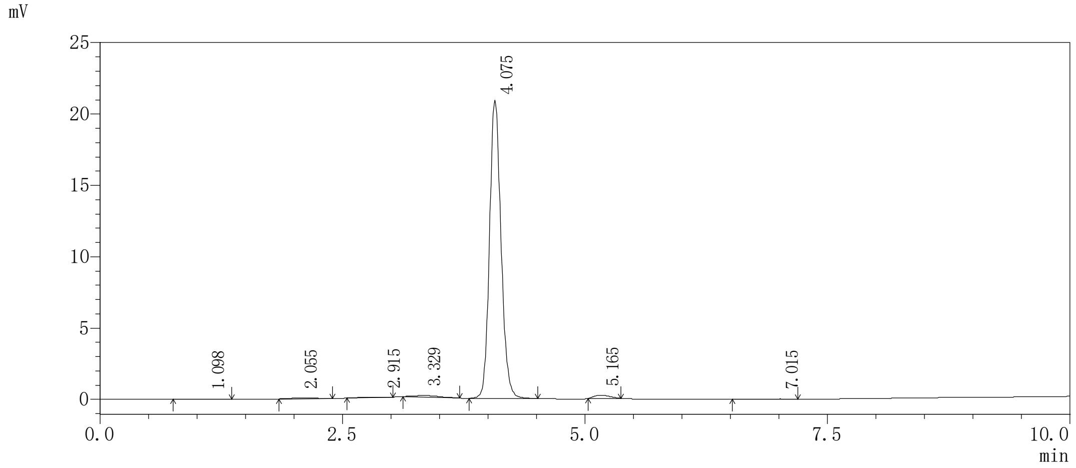

Peak list

| Peak# | RetTime [min] | Area [mV*s] | Height [mV] | Area % | Height % |
|---|---|---|---|---|---|
| 1 | 1.098 | 381 | 20 | 0.212 | 0.212 |
| 2 | 2 | 2.055 | 1207 | 0.671 | 0.671 |
| 3 | 3 | 2.915 | 385 | 0.214 | 0.214 |
| 4 | 4 | 3.329 | 2016 | 1.120 | 1.120 |
| 5 | 5 | 4.075 | 173193 | 96.214 | 96.214 |
| 6 | 6 | 5.165 | 2533 | 1.407 | 1.407 |
| 7 | 7 | 7.015 | 291 | 0.162 | 0.162 |
| Total | 总计 | | 180007 | 100.000 | 100.000 |

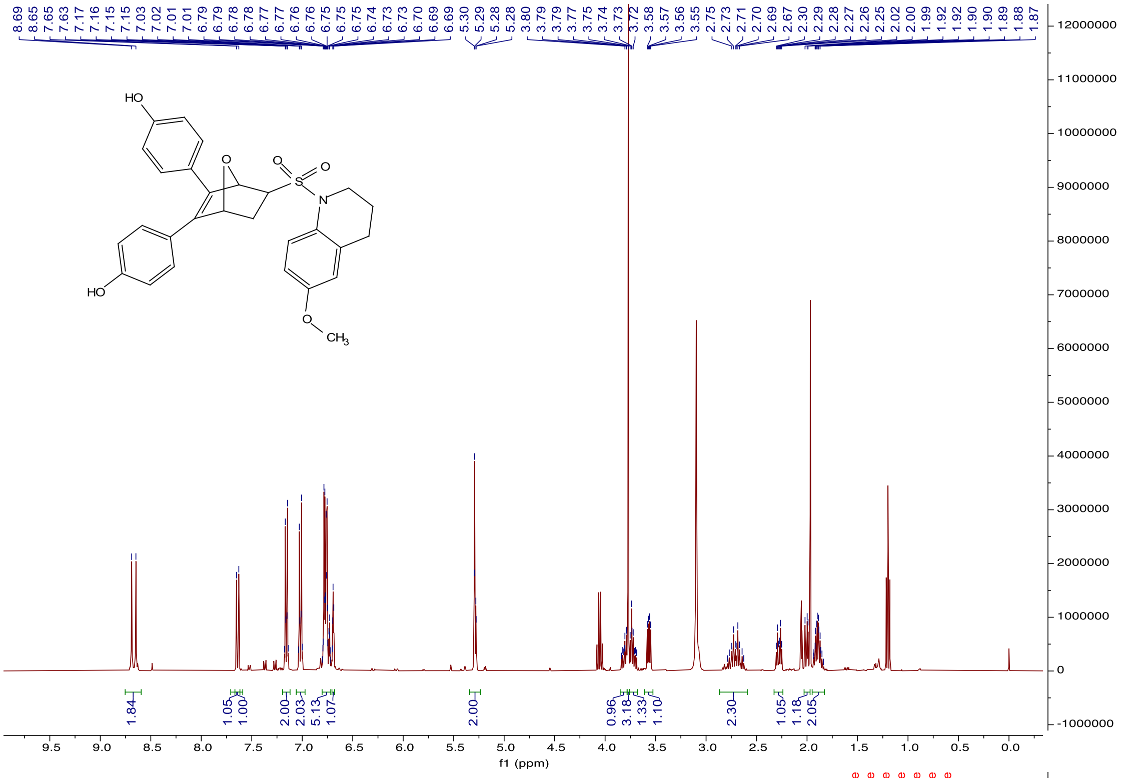

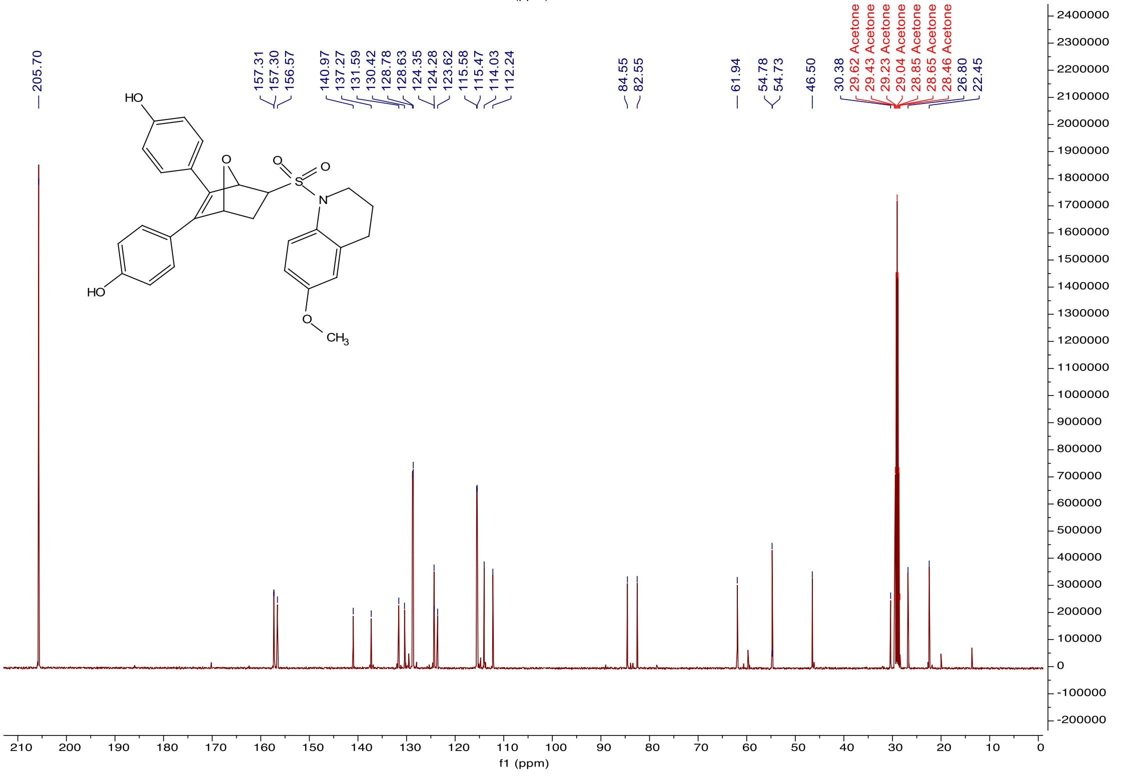

*4,4'-(5-((3,4-dihydroquinolin-1(2H)-yl)sulfonyl)-7-oxabicyclo[2.2.1]hept-2-ene-2,3-diyl) diphenol (**k-410**)*. Following the general procedure for Diels-Alder reaction between **4b** and **2**, compound **k-410** was obtained as yellow solid (Yield 53%). $^{1}$H NMR (400 MHz, Acetone-$d_6$) δ 8.61 (s, 1H), 8.58 (s, 1H), 7.64 (d, J = 8.3 Hz, 1H), 7.19 – 7.13 (m, 2H), 7.05 – 6.97 (m, 2H), 6.94 (dd, J = 11.2, 2.4 Hz, 2H), 6.78 (q, J = 2.5 Hz, 2H), 6.75 (q, J = 2.5 Hz, 2H), 5.32 (d, J = 1.2 Hz, 1H), 5.29 (dd, J = 4.4, 1.1 Hz, 1H), 3.86 – 3.70 (m, 2H), 3.61 (dd, J = 8.3, 4.5 Hz, 1H), 2.81 – 2.61 (m, 2H), 2.31 (dt, J = 11.8, 4.5 Hz, 1H), 2.00 (dd, J = 11.9, 8.4 Hz, 1H), 1.89 (pd, J = 7.1, 4.6 Hz, 2H). $^{13}$C NMR (101 MHz, Acetone-*$d_6$*) δ 157.30, 157.27, 141.00, 137.27, 134.97, 133.51, 130.16, 129.55, 129.51, 128.71, 128.68, 127.12, 124.27, 123.63, 122.33, 115.56, 115.50, 84.49, 82.60, 61.97, 59.75, 46.66, 30.31, 26.71, 22.55, 19.87, 13.67. HRMS (m/z): $[M + Na]^+$ calcd for $C_{27}H_{25}NO_5SNa$, 498.1345; found, 498.1346.

HPLC chromatogram report of **k-410**

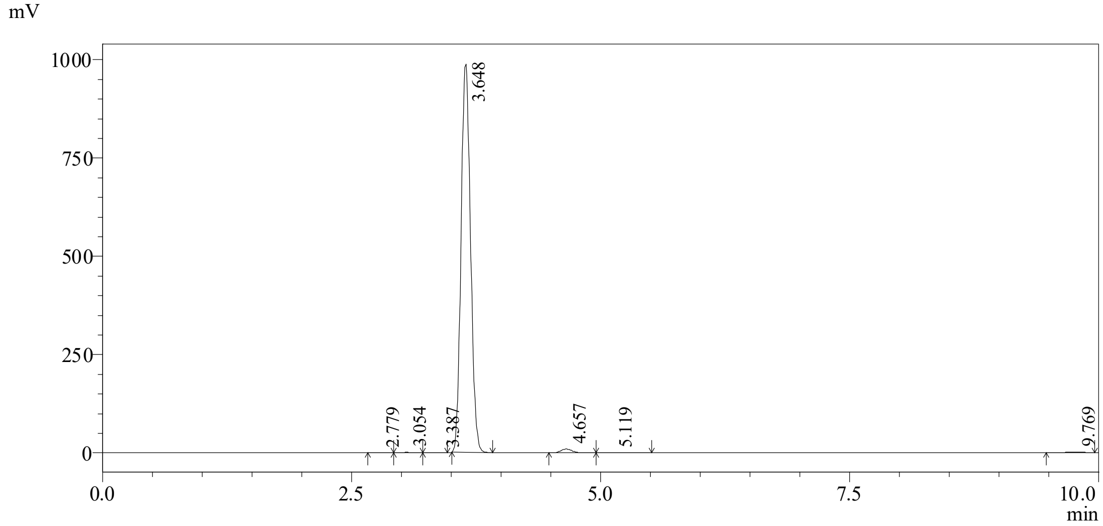

Peak list

| Peak# | RetTime [min] | Area [mV*s] | Height [mV] | Area % | Height % |
|---|---|---|---|---|---|
| 1 | 2.779 | 10486 | 1435 | 0.157 | 0.157 |
| 2 | 3.054 | 14620 | 1507 | 0.219 | 0.219 |
| 3 | 3.387 | 9493 | 954 | 0.142 | 0.142 |
| 4 | 3.648 | 6508650 | 988119 | 97.686 | 97.686 |
| 5 | 4.657 | 79793 | 9918 | 1.198 | 1.198 |
| 6 | 5.119 | 11602 | 927 | 0.174 | 0.174 |
| 7 | 9.769 | 28211 | 2018 | 0.423 | 0.423 |
| Total | | 6662855 | 1004878 | 100.000 | 100.000 |

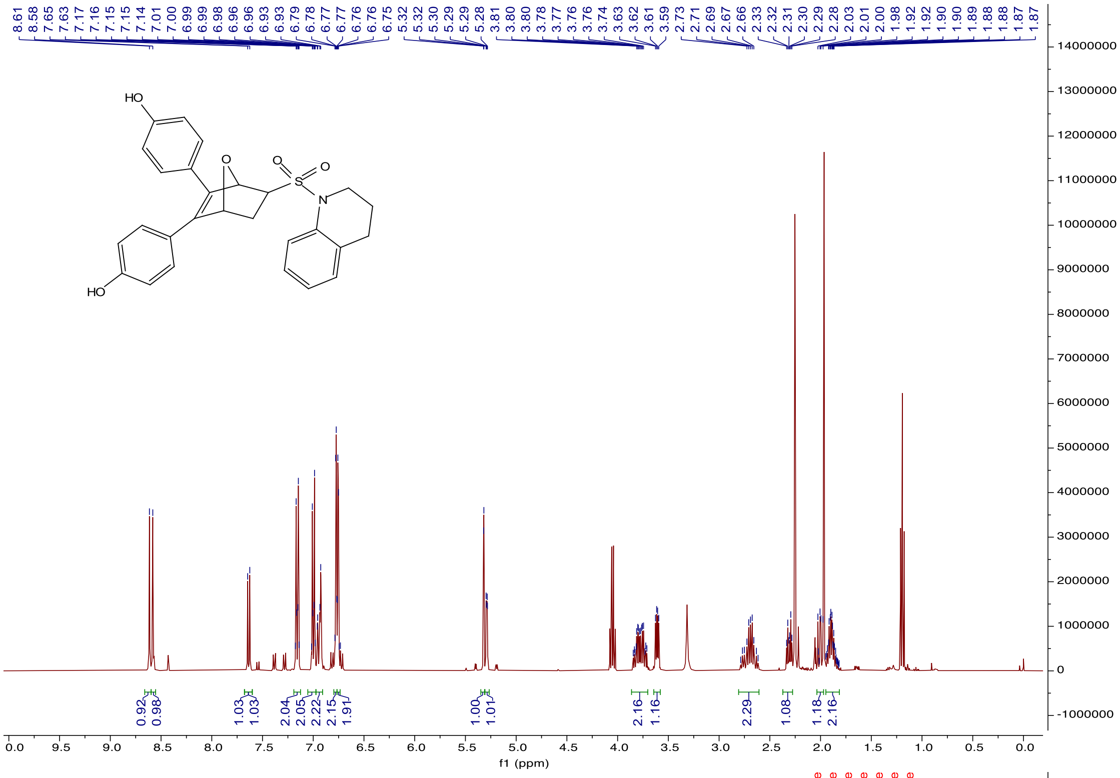
8.61 8.58 7.65 7.63 7.17 7.16 7.15 7.15 7.14 7.01 7.00 6.99 6.99 6.98 6.96 6.96 6.93 6.93 6.79 6.78 6.77 6.77 6.76 6.76 6.75 5.32 5.32 5.30 5.29 5.29 5.28 3.81 3.80 3.80 3.78 3.77 3.76 3.76 3.74 3.63 3.62 3.61 3.59 2.73 2.71 2.69 2.67 2.66 2.33 2.32 2.31 2.30 2.29 2.28 2.03 2.01 2.00 1.98 1.92 1.92 1.90 1.89 1.88 1.88 1.87 1.87
HO
O
O
S
O
N
HO
0.92 0.98 1.03 1.03 2.04 2.05 2.22 2.15 1.91 1.00 1.01 2.16 1.16 2.29 1.08 1.18 2.16
f1 (ppm)

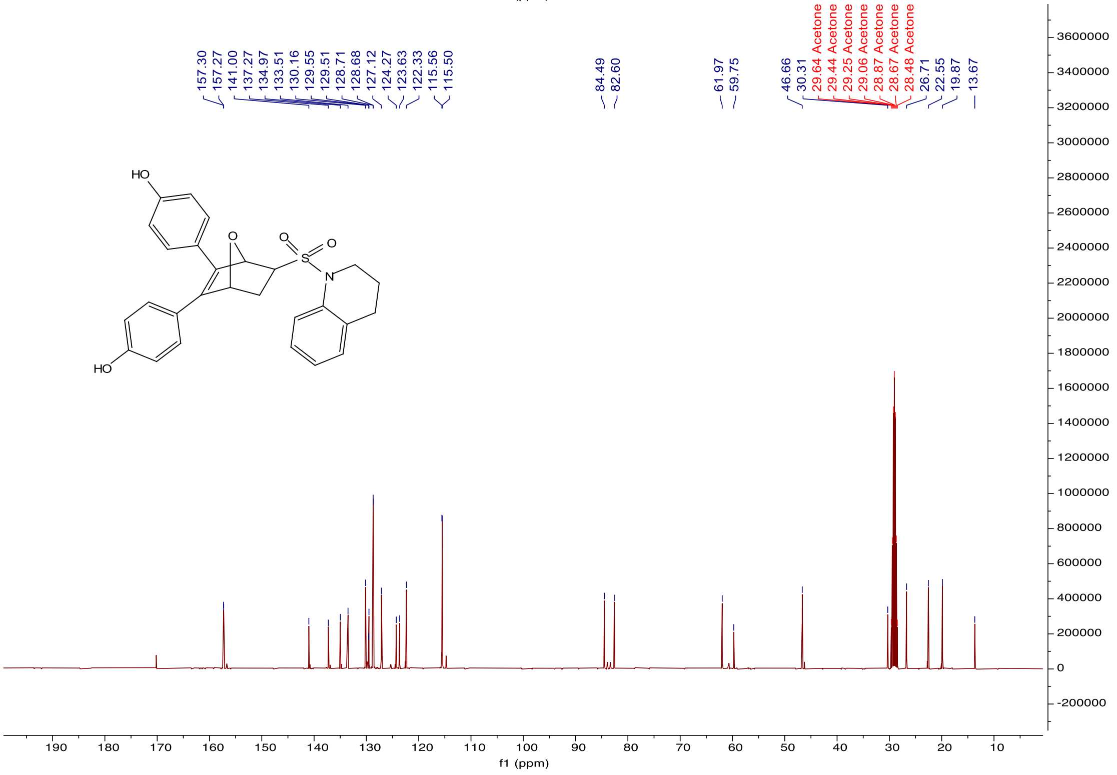
157.30 157.27 141.00 137.27 134.97 133.51 130.16 129.55 129.51 128.71 128.68 127.12 124.27 123.63 122.33 115.56 115.50
84.49 82.60
61.97 59.75
46.66 30.31 29.64 Acetone 29.44 Acetone 29.25 Acetone 29.06 Acetone 28.87 Acetone 28.67 Acetone 28.48 Acetone 26.71 22.55 19.87 13.67
HO
O
O
S
O
N
HO
f1 (ppm)

*4,4'-(5-(indolin-1-ylsulfonyl)-7-oxabicyclo[2.2.1]hept-2-ene-2,3-diyl)diphenol* ***(k-411)***. Following the general procedure for Diels-Alder reaction between **4a** and **2**, compound **k-411** was obtained as yellow solid (Yield 56%). $^{1}$H NMR (400 MHz, Acetone-d$_6$) δ 8.63 (d, J = 1.2 Hz, 2H), 7.46 (d, J = 8.0 Hz, 1H), 7.24 (d, J = 7.4 Hz, 1H), 7.20 – 7.13 (m, 3H), 7.04 – 6.98 (m, 3H), 6.81 – 6.70 (m, 4H), 5.48 (d, J = 1.2 Hz, 1H), 5.31 (dd, J = 4.4, 1.1 Hz, 1H), 4.14 – 4.07 (m, 2H), 3.67 (dd, J = 8.4, 4.4 Hz, 1H), 3.12 (td, J = 8.2, 5.0 Hz, 2H), 2.40 (dt, J = 11.9, 4.5 Hz, 1H), 2.01 (dd, J = 11.9, 8.4 Hz, 1H). $^{13}$C NMR (101 MHz, Acetone-*d$_6$*) δ170.19, 157.36, 157.23, 142.62, 140.91, 137.12, 132.00, 128.85, 128.45, 127.53, 125.56, 124.19, 123.62, 123.33, 115.57, 115.51, 113.83, 84.29, 82.79, 61.19, 59.75, 50.34, 30.12, 27.84, 20.02, 17.98, 13.66. HRMS (m/z): [M + Na]$^+$ calcd for $C_{26}H_{23}NO_5SNa$, 484.1189; found, 484.1192.

HPLC chromatogram report of **k-411**

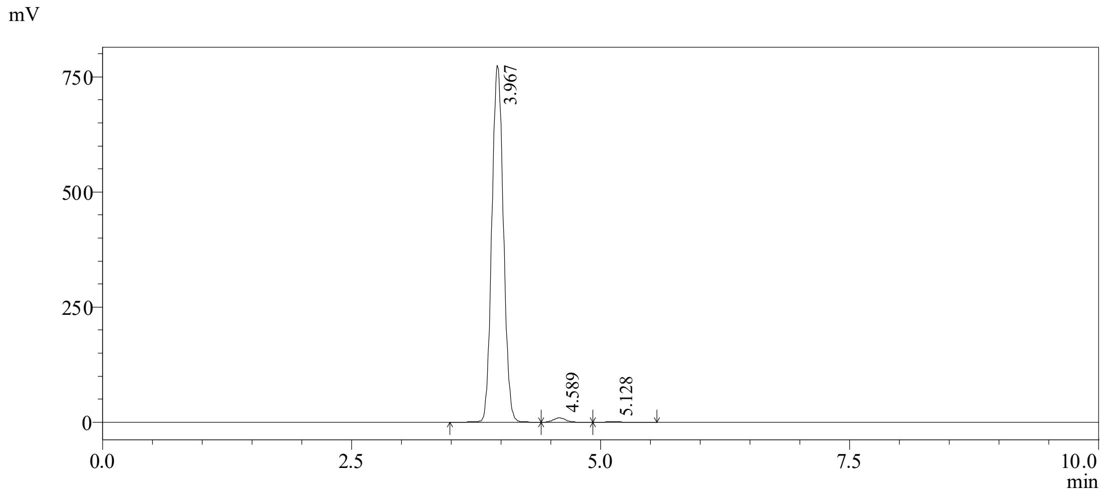

Peak list

| Peak# | RetTime [min] | Area [mV*s] | Height [mV] | Area % | Height % |
|---|---|---|---|---|---|
| 1 | 3.967 | 6269823 | 774923 | 98.488 | 98.488 |
| 2 | 4.589 | 83565 | 9360 | 1.313 | 1.313 |
| 3 | 5.128 | 12709 | 1020 | 0.200 | 0.200 |
| Total | | 6366097 | 785303 | 100.000 | 100.000 |

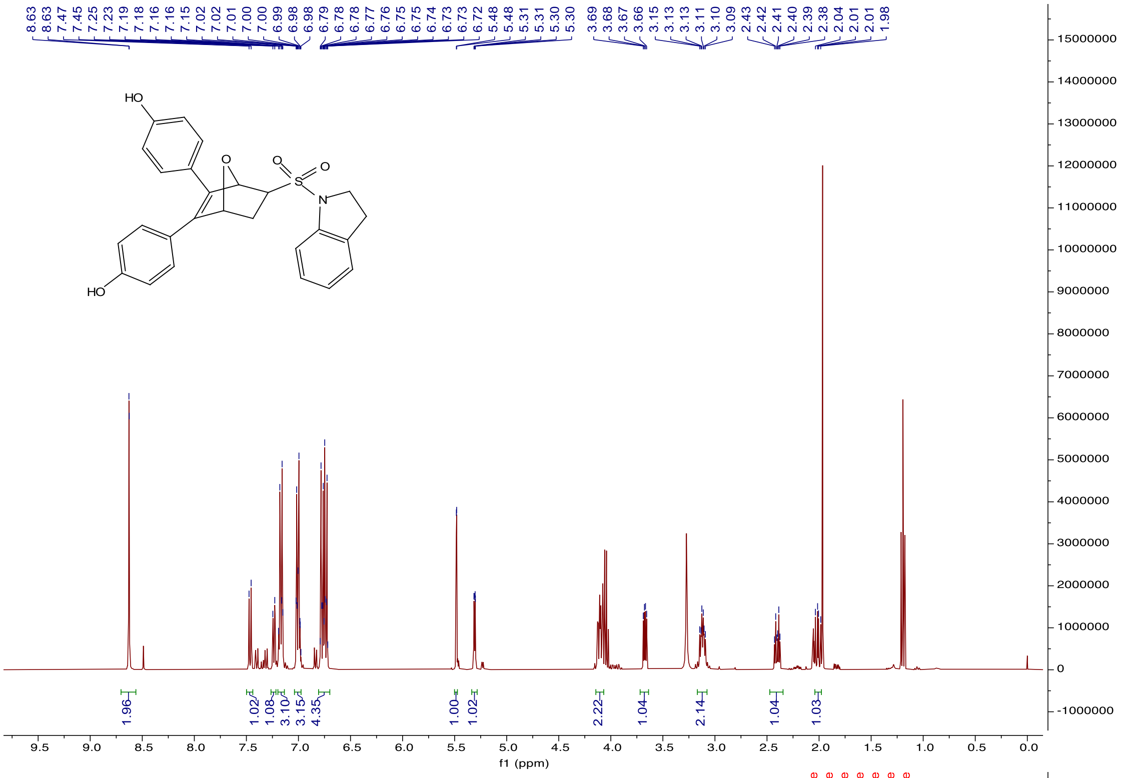
8.63
8.63
7.47
7.45
7.25
7.23
7.19
7.18
7.16
7.16
7.15
7.02
7.02
7.01
7.00
7.00
6.99
6.98
6.98
6.79
6.78
6.78
6.77
6.76
6.75
6.74
6.73
6.73
6.72
5.48
5.48
5.31
5.31
5.30
5.30
3.69
3.68
3.67
3.66
3.15
3.13
3.13
3.11
3.10
3.09
2.43
2.42
2.41
2.40
2.39
2.38
2.04
2.01
2.01
1.98
HO
O
O
S
O
N
HO
1.96
1.02
1.08
3.10
3.15
4.35
1.00
1.02
2.22
1.04
2.14
1.04
1.03
f1 (ppm)

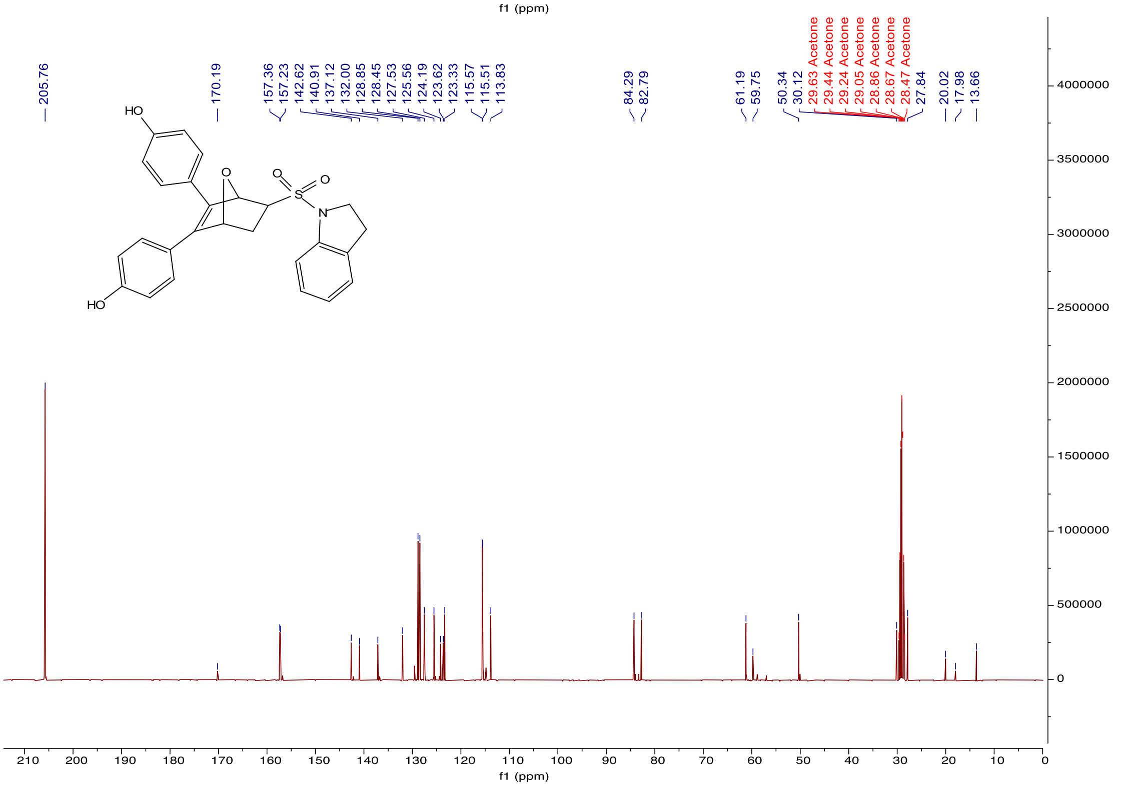
205.76
170.19
157.36
157.23
142.62
140.91
137.12
132.00
128.85
128.45
127.53
125.56
124.19
123.62
123.33
115.57
115.51
113.83
84.29
82.79
61.19
59.75
50.34
30.12
29.63 Acetone
29.44 Acetone
29.24 Acetone
29.05 Acetone
28.86 Acetone
28.67 Acetone
28.47 Acetone
27.84
20.02
17.98
13.66
HO
O
O
S
O
N
HO
f1 (ppm)

*5,6-bis(4-hydroxyphenyl)-N-isopentyl-N-(4-methoxyphenyl)-7-oxabicyclo[2.2.1]hept-5-ene-2-sulfonamide (**k-422**)*. Following the general procedure for Diels-Alder reaction between **1h** and **2**, compound **k-422** was obtained as white solid (Yield 45%). $^{1}$H NMR (400 MHz, Acetonitrile-$d_3$) δ 7.19 – 7.10 (m, 6H), 6.84 – 6.72 (m, 6H), 5.40 (d, J = 1.3 Hz, 1H), 5.26 (dd, J = 4.4, 1.3 Hz, 1H), 3.77 (s, 3H), 3.71 – 3.63 (m, 2H), 3.36 (dd, J = 8.3, 4.5 Hz, 1H), 2.11 (s, 1H), 2.01 (dd, J = 12.0, 8.4 Hz, 1H), 1.59 (dd, J = 13.4, 6.7 Hz, 1H), 1.22 (q, J = 6.9 Hz, 2H), 0.83 (dd, J = 6.7, 1.0 Hz, 6H). $^{13}$C NMR (101 MHz, Acetonitrile-$d_3$) δ 159.54, 157.33, 132.16, 130.99, 129.87, 129.04, 116.16, 115.93, 114.64, 85.00, 83.25, 61.27, 55.69, 50.15, 37.76, 30.87, 25.44, 22.13. HRMS (m/z): $[M + Na]^+$ calcd for $C_{30}H_{33}NO_6SNa$, 558.1921; found, 558.1926.

HPLC chromatogram report of **k-422**

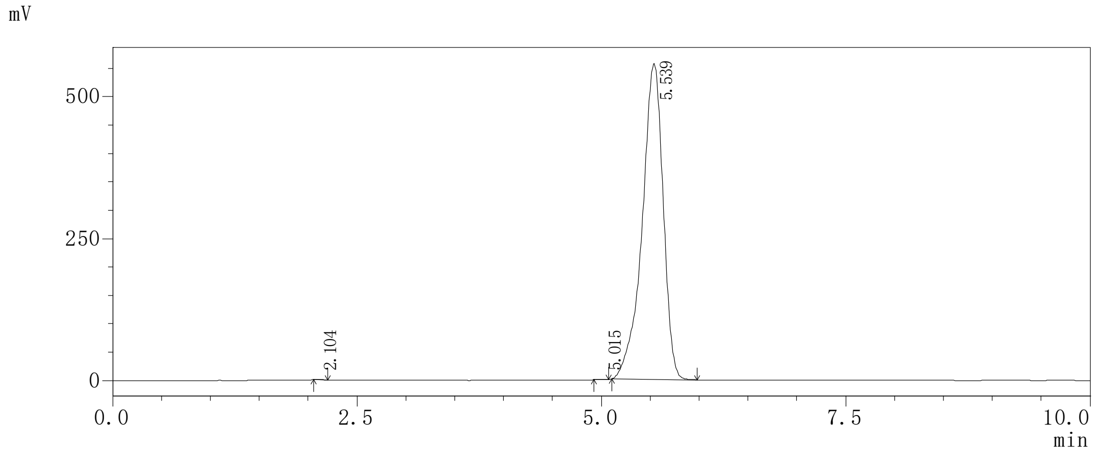

Peak list

| Peak# | RetTime [min] | Area [mV*s] | Height [mV] | Area % | Height % |
|---|---|---|---|---|---|
| 1 | 2.104 | 2619 | 563 | 0.031 | 0.031 |
| 2 | 5.015 | 1866 | 328 | 0.022 | 0.022 |
| 3 | 5.539 | 8331009 | 556338 | 99.946 | 99.946 |
| Total | | 8335493 | 557229 | 100.000 | 100.000 |

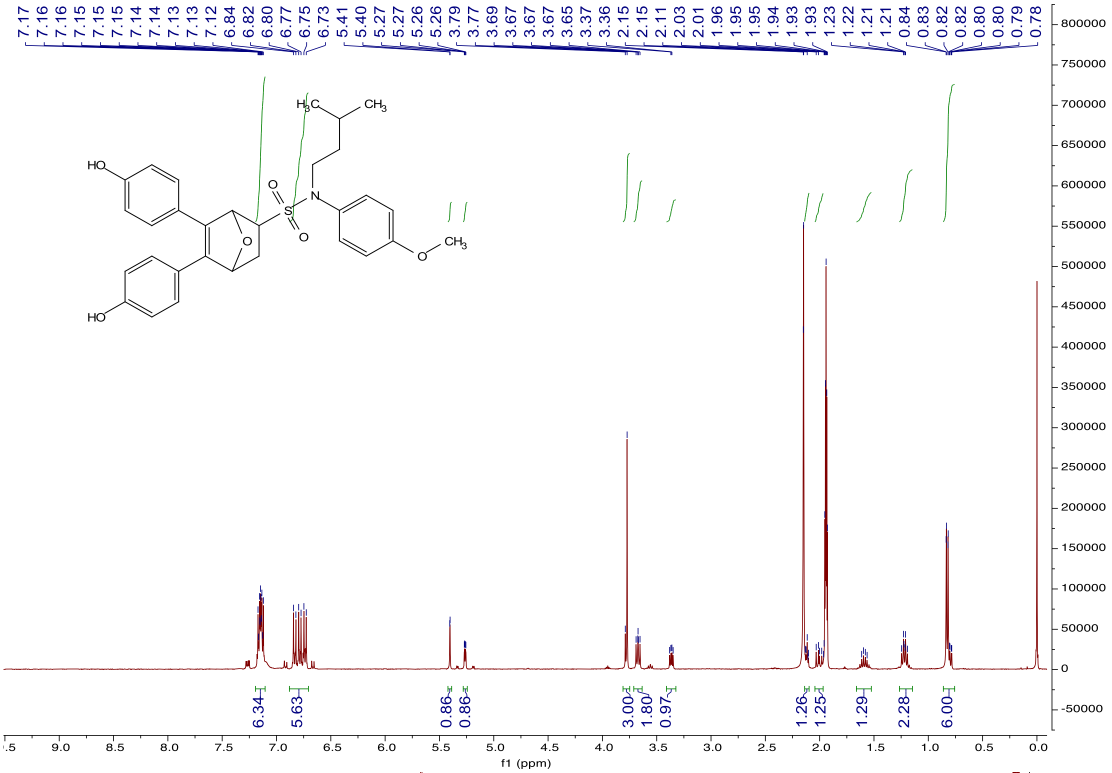

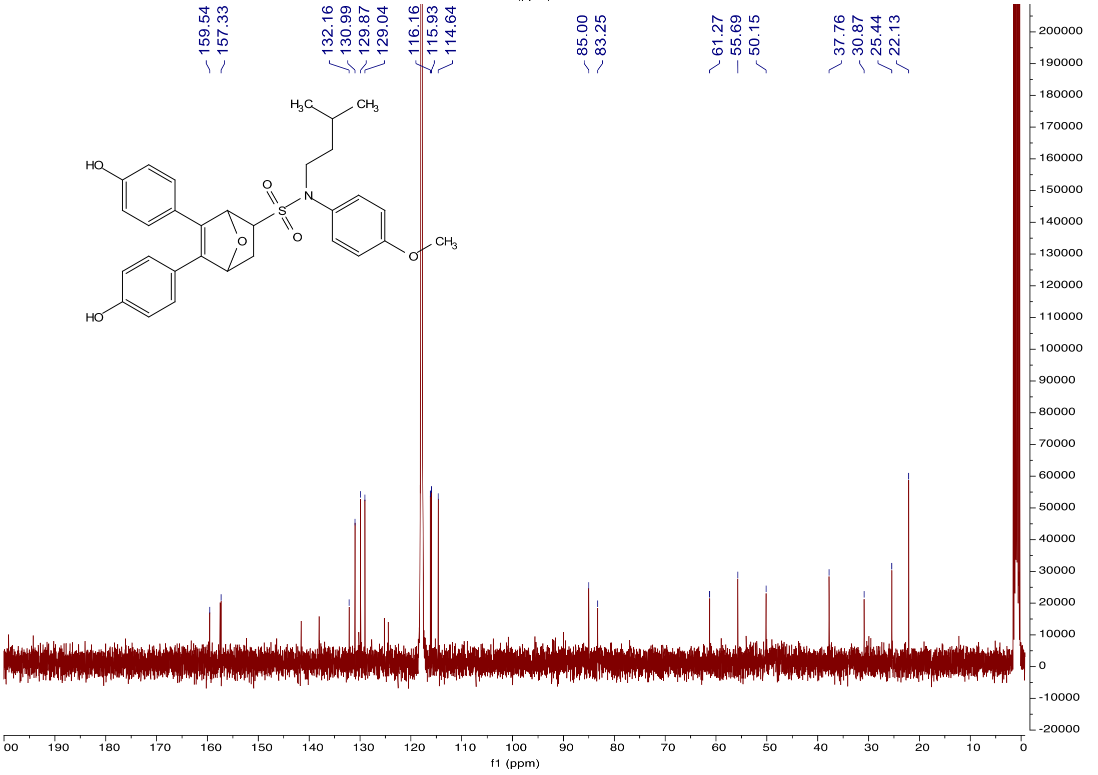

*N-(cyclobutylmethyl)-5,6-bis(4-hydroxyphenyl)-N-(4-methoxyphenyl)-7-oxabicyclo[2.2.1]hept-5-ene-2-sulfonamide (**k-424**)*. Following the general procedure for Diels-Alder reaction between **1i**

and **2**, compound **k-424** was obtained as white solid (Yield 48%). [1]H NMR (400 MHz, Acetonitrile-$d_3$) δ 7.13 (qd, J = 6.4, 2.1 Hz, 6H), 6.89 – 6.66 (m, 6H), 5.41 (d, J = 1.3 Hz, 1H), 5.27 (dd, J = 4.4, 1.3 Hz, 1H), 3.77 (s, 3H), 3.66 (d, J = 7.5 Hz, 2H), 3.36 (dd, J = 8.3, 4.5 Hz, 1H), 2.28 – 2.20 (m, 1H), 2.12 (d, J = 4.4 Hz, 1H), 2.01 (dd, J = 12.0, 8.4 Hz, 1H), 1.90 – 1.82 (m, 2H), 1.80 – 1.74 (m, 2H), 1.63 (ddd, J = 11.0, 6.9, 2.8 Hz, 2H). [13]C NMR (151 MHz, Acetonitrile-$d_3$) δ 159.02, 156.75, 141.02, 137.51, 130.50, 128.46, 115.60, 115.36, 114.01, 84.42, 82.67, 60.80, 56.49, 55.12, 34.22, 30.28, 25.38, 17.74. HRMS (m/z): $[M + Na]^+$ calcd for $C_{30}H_{31}NO_6SNa$, 556.1764; found, 556.1765.

HPLC chromatogram report of **k-424**

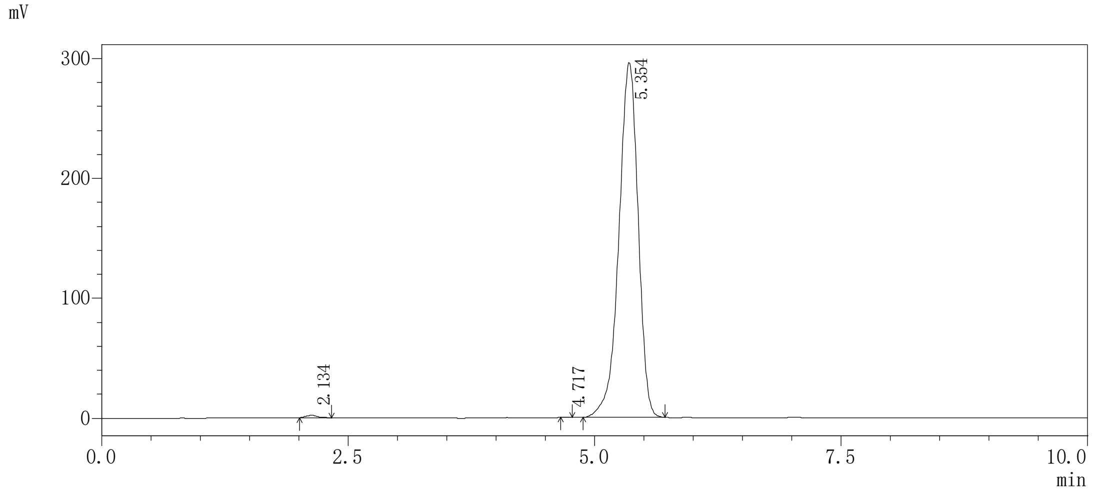

Peak list

| Peak# | RetTime [min] | Area [mV*s] | Height [mV] | Area % | Height % |
|---|---|---|---|---|---|
| 1 | 2.134 | 14921 | 1868 | 0.370 | 0.370 |
| 2 | 4.717 | 718 | 158 | 0.018 | 0.018 |
| 3 | 5.354 | 4014176 | 295579 | 99.612 | 99.612 |
| Total | | 4029815 | 297606 | 100.000 | 100.000 |

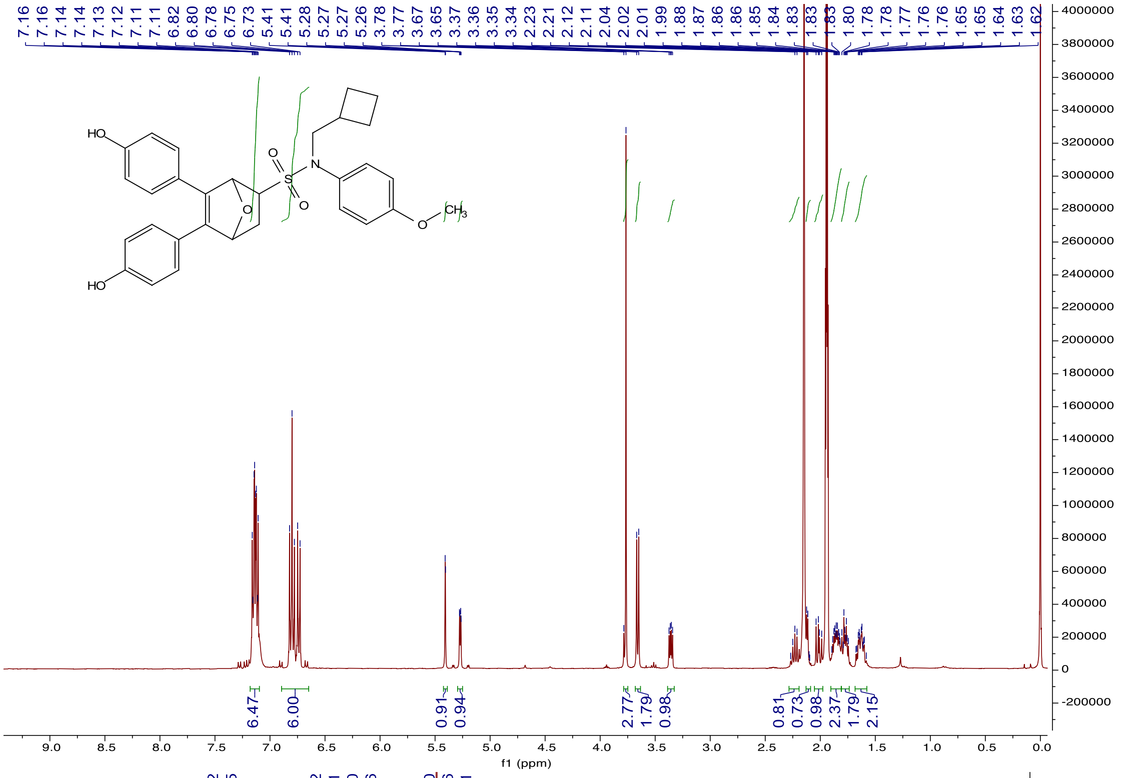

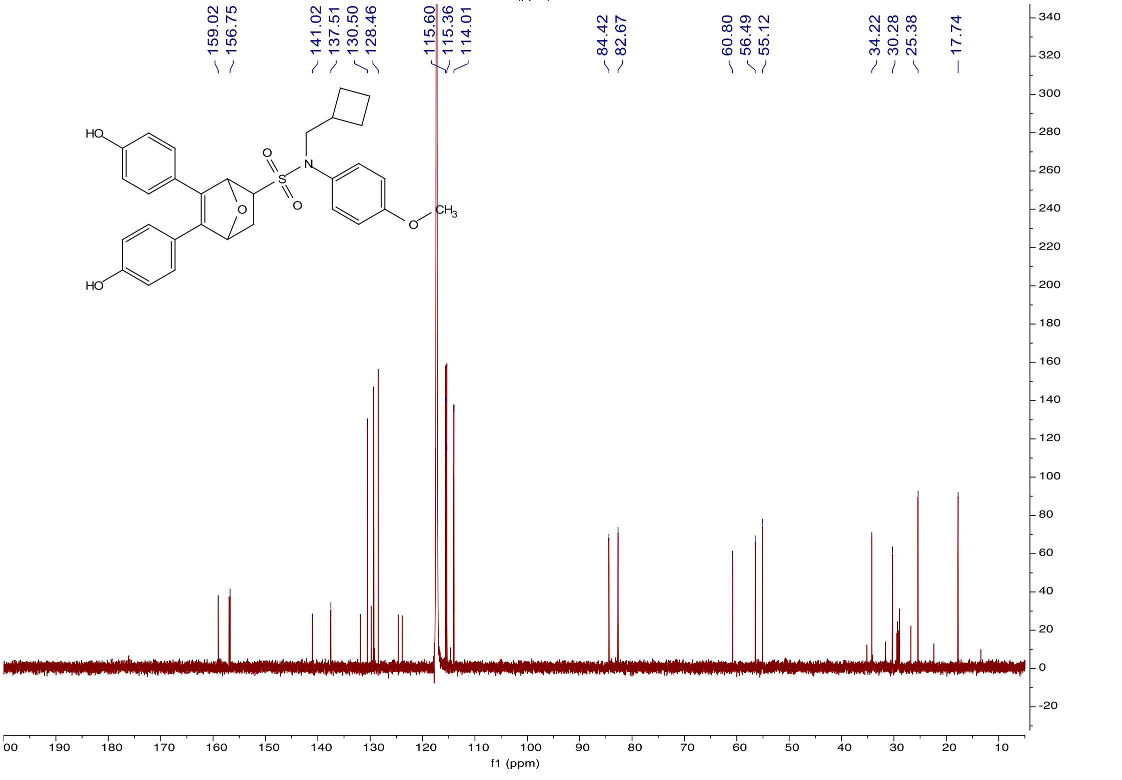

**SI References**

1. N. E. Bruno *et al.*, Activation of Crtc2/Creb1 in skeletal muscle enhances weight loss during intermittent fasting. *FASEB J* **35**, e21999 (2021).
2. J. C. Nwachukwu *et al.*, Predictive features of ligand-specific signaling through the estrogen receptor. *Mol Syst Biol* **12**, 864 (2016).
3. A. Bergamaschi *et al.*, The forkhead transcription factor FOXM1 promotes endocrine resistance and invasiveness in estrogen receptor-positive breast cancer by expansion of stem-like cancer cells. *Breast Cancer Res* **16**, 436 (2014).
4. K. W. Nettles *et al.*, NFkappaB selectivity of estrogen receptor ligands revealed by comparative crystallographic analyses. *Nat Chem Biol* **4**, 241-247 (2008).
5. C. Vonrhein *et al.*, Data processing and analysis with the autoPROC toolbox. *Acta Crystallogr D Biol Crystallogr* **67**, 293-302 (2011).
6. P. D. Adams *et al.*, PHENIX: building new software for automated crystallographic structure determination. *Acta Crystallogr D Biol Crystallogr* **58**, 1948-1954 (2002).
7. N. W. Moriarty, R. W. Grosse-Kunstleve, P. D. Adams, electronic Ligand Builder and Optimization Workbench (eLBOW): a tool for ligand coordinate and restraint generation. *Acta Crystallogr D Biol Crystallogr* **65**, 1074-1080 (2009).
8. P. Emsley, K. Cowtan, Coot: model-building tools for molecular graphics. *Acta Crystallogr D Biol Crystallogr* **60**, 2126-2132 (2004).
9. R. P. Joosten, F. Long, G. N. Murshudov, A. Perrakis, The PDB_REDO server for macromolecular structure model optimization. *IUCrJ* **1**, 213-220 (2014).
10. B. Webb, A. Sali, Comparative Protein Structure Modeling Using MODELLER. *Curr Protoc Bioinformatics* **54**, 5 6 1-5 6 37 (2016).
11. E. F. Pettersen *et al.*, UCSF Chimera--a visualization system for exploratory research and analysis. *J Comput Chem* **25**, 1605-1612 (2004).
12. J. A. Maier *et al.*, ff14SB: Improving the Accuracy of Protein Side Chain and Backbone Parameters from ff99SB. *J Chem Theory Comput* **11**, 3696-3713 (2015).
13. R. Brust *et al.*, A structural mechanism for directing corepressor-selective inverse agonism of PPARgamma. *Nat Commun* **9**, 4687 (2018).
14. C. W. Hopkins, S. Le Grand, R. C. Walker, A. E. Roitberg, Long-Time-Step Molecular Dynamics through Hydrogen Mass Repartitioning. *J Chem Theory Comput* **11**, 1864-1874 (2015).
15. L. C. Pierce, R. Salomon-Ferrer, F. d. O. C. Augusto, J. A. McCammon, R. C. Walker, Routine Access to Millisecond Time Scale Events with Accelerated Molecular Dynamics. *J Chem Theory Comput* **8**, 2997-3002 (2012).
16. D. R. Roe, T. E. Cheatham, 3rd, PTRAJ and CPPTRAJ: Software for Processing and Analysis of Molecular Dynamics Trajectory Data. *J Chem Theory Comput* **9**, 3084-3095 (2013).
17. B. J. Grant, A. P. Rodrigues, K. M. ElSawy, J. A. McCammon, L. S. Caves, Bio3d: an R package for the comparative analysis of protein structures. *Bioinformatics* **22**, 2695-2696 (2006).
18. X. Q. Yao *et al.*, Dynamic Coupling and Allosteric Networks in the alpha Subunit of Heterotrimeric G Proteins. *J Biol Chem* **291**, 4742-4753 (2016).
19. C. Galaxy, The Galaxy platform for accessible, reproducible and collaborative biomedical analyses: 2022 update. *Nucleic Acids Res* **50**, W345-W351 (2022).

20. D. Kim, B. Langmead, S. L. Salzberg, HISAT: a fast spliced aligner with low memory requirements. *Nat Methods* **12**, 357-360 (2015).
21. Y. Liao, G. K. Smyth, W. Shi, featureCounts: an efficient general purpose program for assigning sequence reads to genomic features. *Bioinformatics* **30**, 923-930 (2014).
22. M. I. Love, W. Huber, S. Anders, Moderated estimation of fold change and dispersion for RNA-seq data with DESeq2. *Genome Biol* **15**, 550 (2014).
23. M. Zhu *et al.*, Bicyclic core estrogens as full antagonists: synthesis, biological evaluation and structure-activity relationships of estrogen receptor ligands based on bridged oxabicyclic core arylsulfonamides. *Org Biomol Chem* **10**, 8692-8700 (2012).
